# A comprehensive view of cell-type-specific temporal dynamics in human and mouse brains

**DOI:** 10.1101/2022.10.01.509820

**Authors:** Ziyu Lu, Melissa Zhang, Jasper Lee, Andras Sziraki, Sonya Anderson, Shaoyu Ge, Peter T. Nelson, Wei Zhou, Junyue Cao

**Author notes:** These authors contributed equally (Ziyu Lu, Melissa Zhang). Senior author (Wei Zhou, Junyue Cao). Correspondence (W.Z.), (J.C.).

## Abstract

Progenitor cells play fundamental roles in preserving optimal organismal functions under normal, aging, and disease conditions. However, progenitor cells are incompletely characterized, especially in the brain, partly because conventional methods are restricted by inadequate throughput and resolution for deciphering cell-type-specific proliferation and differentiation dynamics *in vivo*. Here, we developed *TrackerSci*, a new technique that combines *in vivo* labeling of newborn cells with single-cell combinatorial indexing to profile the single-cell chromatin landscape and transcriptome of rare progenitor cells and track cellular differentiation trajectories *in vivo*. We applied *TrackerSci* to analyze the epigenetic and gene expression dynamics of newborn cells across entire mouse brains spanning three age stages and in a mouse model of Alzheimer’s disease. Leveraging the dataset, we identified diverse progenitor cell types less-characterized in conventional single cell analysis, and recovered their unique epigenetic signatures. We further quantified the cell-type-specific proliferation and differentiation potentials of progenitor cells, and identified the molecular programs underlying their aging-associated changes (*e.g.,* reduced neurogenesis/oligodendrogenesis). Finally, we expanded our analysis to study progenitor cells in the aged human brain through profiling ∼800,000 single-cell transcriptomes across five anatomical regions from six aged human brains. We further explored the transcriptome signatures that are shared or divergent between human and mouse oligodendrogenesis, as well as the region-specific down-regulation of oligodendrogenesis in the human cerebellum. Together, the data provide an in-depth view of rare progenitor cells in mammalian brains. We anticipate *TrackerSci* will be broadly applicable to characterize cell-type-specific temporal dynamics in diverse systems.

## Introduction

New neurons and glial cells are continuously produced in the adult mammalian brains, a critical process associated with memory, learning, and stress (Lugert et al., 2010; Spalding et al., 2013). There is a consensus that adult neurogenesis and oligodendrogenesis decline with advancing ages and in neuropathological conditions (Galvan and Jin, 2007; Pollina and Brunet, 2011), but to what extent is debated (Mathews et al., 2017; Sorrells et al., 2018). The ambiguity stems partly from technical limitations - most studies rely upon the utilization of proxy markers, which may introduce bias for quantifying the dynamics of extremely rare progenitor cells, especially in aged tissues. Furthermore, the identity of progenitor cells is established as a result of tightly controlled epigenetic programs, driven in part by transcription factors that interact with cis-regulatory sequences in a cell-type-specific manner. While previous single-cell studies have provided critical insight into the gene expression signatures of progenitor cells in the adult brain (Franjic et al., 2022; Habib et al., 2016; Kalinina and Lagace, 2022), little is known about how the epigenetic landscape regulates the dynamics of rare progenitor cells *in vivo*. Therefore, novel approaches for quantitatively capturing newborn cells and tracking their transcriptome and chromatin state changes are critical to understanding cell population dynamics in development, aging, and disease states.

Here we describe a novel method, *TrackerSci*, to track the proliferation and differentiation dynamics of newborn cells in the mammalian brain. *TrackerSci* integrated protocols for labeling newly synthesized DNA with a thymidine analog 5-Ethynyl-2-deoxyuridine (EdU) (Salic and Mitchison, 2008) and single-cell combinatorial indexing sequencing for both transcriptome (Cao et al., 2019) and chromatin accessibility profiling (Cusanovich et al., 2015). As a demonstration, we applied *TrackerSci* to profiling the single-cell transcriptome or chromatin accessibility dynamics of 14,689 newborn cells from entire mouse brains spanning three age stages and two genotypes. With the resulting datasets, we recovered rare progenitor cell populations less represented in conventional single-cell analysis and tracked their cell-type-specific proliferation and differentiation dynamics across ages. Furthermore, we identified the genetic and epigenetic signatures associated with the alteration of cellular dynamics (*e.g.,* adult neurogenesis, oligodendrogenesis) that occurs in the aged mammalian brain. Finally, to compare rare progenitor cells across species, we generated a human brain cell atlas profiling ∼800,000 single-nucleus transcriptomes of the human brain across five anatomic regions. By integration analysis with the *TrackerSci* dataset, we identified region- and cell-type-specific signatures of rare progenitor cells in the aged human brain and recovered conserved and divergent molecular signatures of oligodendrogenesis cells between human and mouse. The experimental and computational methods described here could be broadly applied to track cellular regenerative capacity and differentiation potential across mammalian organs and other biological systems.

## Results

### Overview of *TrackerSci*

The optimized *TrackerSci* protocol follows these steps (**Figure 1A**): (i) Mice are labeled with 5-Ethynyl-2-deoxyuridine (EdU), a thymidine analog that can be incorporated into replicating DNA for labeling *in vivo* cellular proliferation (Lin et al., 2009; Salic and Mitchison, 2008). (ii) Brains are dissected, and nuclei are extracted, fixed, and then subjected to click chemistry-based *in situ* ligation (Clarke et al., 2017) to an azide-containing fluorophore, followed by fluorescence-activated cell sorting (FACS) to enrich the EdU+ cells (**Figure S1A**). (iii) Indexed reverse transcription or transposition is used to introduce the first round of indexing. Cells from all wells are pooled and then redistributed into multiple 96-well plates through FACS sorting to further purify the EdU+ cells (**Figure S1B**). (iv) We then follow library preparation protocols similar to sci-RNA-seq (Cao et al., 2019) for transcriptome profiling or sci-ATAC-seq (Cusanovich et al., 2015) for chromatin accessibility analysis. Most cells pass through a unique combination of wells, such that their contents are marked by a unique combination of barcodes that can be used to group reads derived from the same cell. Notably, the two sorting steps implemented in *TrackerSci* are essential for excluding contaminating cells and enriching extremely rare proliferating cell populations, especially in the aged brain (less than 0.1% of the total cell population are EdU+ cells).

**Figure 1.**
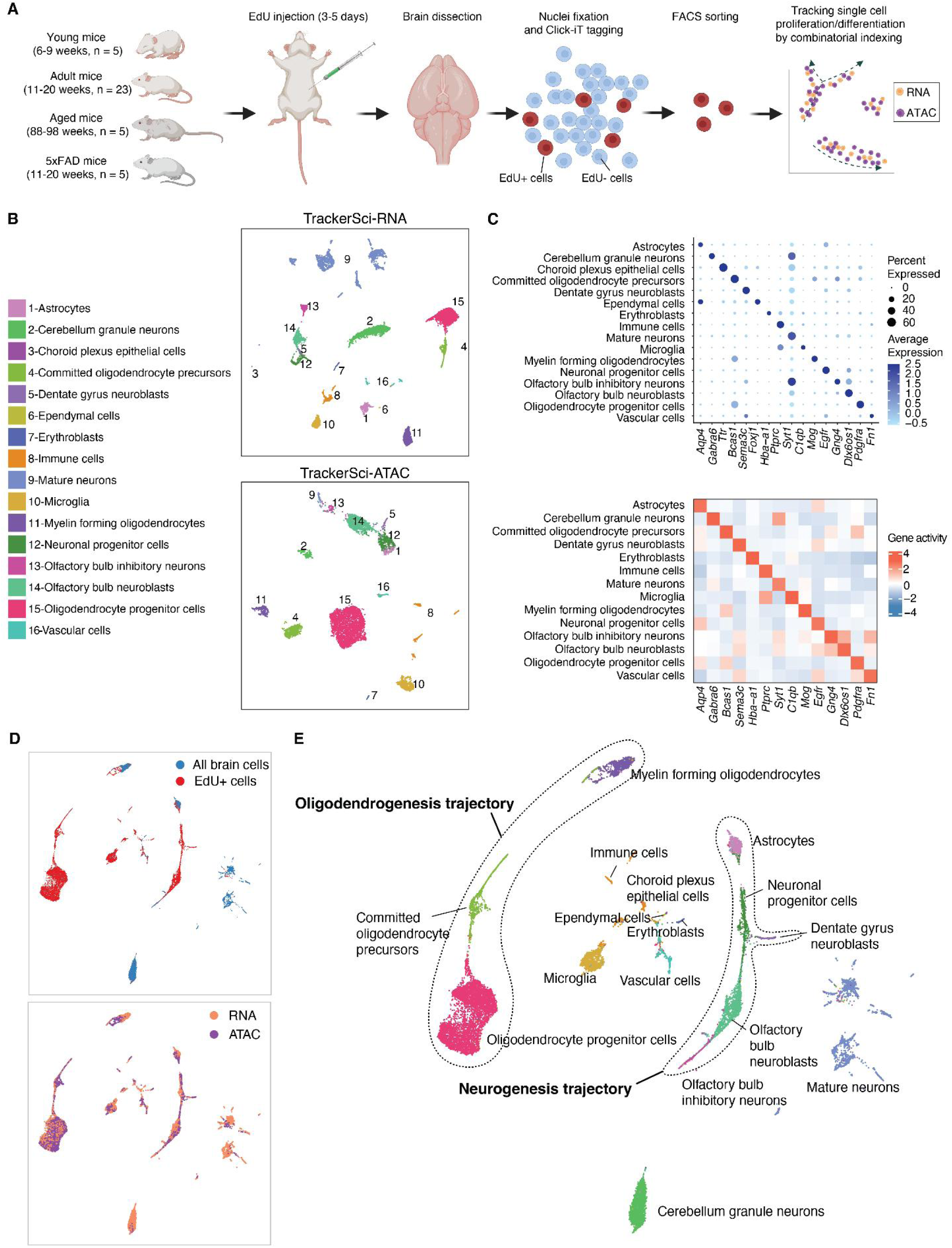
*TrackerSc*i enables single-cell transcriptome and chromatin accessibility profiling of rare proliferating cells in the mammalian brain. (A) *TrackerSci* workflow and experiment scheme. Key steps are outlined in the text. (B) UMAP visualization of single-cell transcriptomes (top) and single-cell chromatin accessibility profiles (bottom), including EdU+ cells (profiled by *TrackerSci*) and all brain cells (without enrichment of EdU+ cells), colored by main cell types. Dimension reduction analysis for scRNA-seq and scATAC-seq was performed independently. (C) Dotplot and heatmap showing gene expression and gene activity of known marker genes for each cluster defined by *TrackerSci-RNA* (top) and *TrackerSci-ATAC* (bottom), respectively. (D-E) UMAP visualization of mouse brain cells, integrating the single-cell transcriptome and chromatin accessibility profiles of EdU+ cells and DAPI singlets (representing the global brain cell population). Cells are colored by sources (D, top), molecular layers (D, bottom), and main cell types (D). The identified neurogenesis and oligodendrogenesis trajectories are both annotated in (E).

We extensively optimized the reaction conditions (*e.g.,* fixation, permeabilization, and click-chemistry reaction) to ensure the approach is fully compatible with FACS sorting and single-cell transcriptome and chromatin accessibility profiling (**Figure S2 and S3**). For instance, the active Cu(I) catalyst and additive included in the conventional click-chemistry reaction (Habib et al., 2016) significantly reduced the nuclei quality for single-cell gene expression analysis (**Figure S2A**). To solve this problem, we tested a commercialized click-chemistry method using picolyl azide dye and copper protectant, which resulted in a minimal defect on library complexity (**Figure S2B, Method**) or cell purity for single-cell RNA-seq analysis, as shown in an experiment profiling a mixture of human HEK293T and mouse NIH/3T3 cells (**Figure S1C and S1D**). As a quality control, we further compared the *TrackerSci* chromatin accessibility profile with the conventional sci-ATAC-seq profile in a mixture of human HEK293T and mouse NIH/3T3 cells. Both methods showed similar cellular purity (**Figure S3A**), fragment length distributions (**Figure S3B**), a comparable number of unique fragments per cell, and a similar ratio of reads overlapping with promoters in both cell lines and mouse brain nuclei (**Figure S3C and S3D**).

Additionally, the aggregated transcriptome and chromatin accessibility profiles derived from *TrackerSci* (both cultured cell lines and tissues) were highly correlated with conventional single-cell combinatorial indexing profiling (**Figure S2E and S3E**), suggesting that the labeling and conjugating reactions (*e.g.,* EdU labeling and click-chemistry) in *TrackerSci* do not substantially interfere with downstream single-cell transcriptome and chromatin accessibility profiling by combinatorial indexing.

### A global view of newborn cells across the mammalian brain

We next applied *TrackerSci* to capture rare newborn cells from mouse brains spanning three age stages and two genotypes. Briefly, following three to five days of continuous EdU labeling, we isolated nuclei from the whole brain of 38 sex-balanced C57BL/6 mice (**Figure 1A**; **Table S1**), including 33 wild-type mice across multiple development stages (Young: 6-9 weeks, Adult: 11-20 weeks, and Aged: 88-98 weeks) as well as five 5xFAD mutant mice (11-20 weeks) harboring multiple Alzheimer’s Disease (AD) mutations (Oakley et al., 2006). Following *TrackerSci* protocol, we obtained transcriptomic profiles for 5,715 newborn cells (median 2,909 UMIs) (**Figure S4A and S4B**) and chromatin accessibility profiles for 8,974 newborn cells (median 50,225 unique reads) (**Figure S5A and S5B**). In addition, to characterize the global brain cell population as a background control, we included DAPI singlets representing ‘all’ brain cells (*i.e.,* without enrichment of the EdU+ cells) and obtained transcriptomic profiles for 8,380 nuclei (median 1,553 UMIs) and chromatin accessibility profiles for 342 nuclei (median 24,521 unique reads). The EdU+ nuclei and DAPI singlets were collected from the same set of samples and processed in parallel to minimize any batch effect.

We first subjected the 14,095 *TrackerSci* transcriptome profiles, including both EdU+ nuclei and DAPI singlets, to Louvain clustering (Blondel et al., 2008) and UMAP visualization (McInnes et al., 2018) (**Figure 1B**; **Figure S4C and S4D**). Sixteen cell clusters were identified and annotated based on established markers (**Figure 1C**; **Table S2**), ranging in size from 25 cells (Choroid plexus epithelial cells) to 3,141 cells (Mature neurons). We next performed a semi-supervised clustering analysis of 9,316 *TrackerSci* chromatin accessibility profiles (8,974 EdU+ nuclei and 342 DAPI singlets), and identified fourteen clusters (**Figure 1B**; **Figure S5C and S5D; Methods**), which mapped 1:1 to the main cell types identified in the transcriptome analysis. Two rare cell types (*i.e.,* ependymal cells and choroid plexus epithelial cells) were only detected in the RNA dataset, mainly due to the low abundance of these cell types. As expected, the corresponding cell types defined by the two molecular layers overlapped well in the integration analysis (**Figure 1D**).

We observed a notably altered distribution of cell-type-specific fractions between ‘all’ brain cells and the EdU+ cells (**Figure 2A**). For example, in contrast to the ‘all’ brain cells that are dominated by mature neurons (*e.g.,* cerebellum granule neurons: 32.7% in DAPI singlets vs. 2.85% in EdU+ cells) and differentiated glial cells (*e.g.,* myelin-forming oligodendrocytes: 11.9 % in DAPI singlets vs. 0.75% in EdU+ cells), the EdU+ population showed prominent enrichment of progenitor cells such as immature neurons (*e.g.,* olfactory bulb neuroblasts: 0.14% in DAPI singlets vs. 13.4% in EdU+ cells) and glia progenitors (*e.g.,* oligodendrocyte progenitor cells: 1.11% in DAPI singlets vs. 45.4% in EdU+ cells). Intriguingly, we detected newly-generated erythroblasts (*Hbb-bt*+, *Hbb-bs*+) and immune cells (*Ptprc*+), which may correspond to newborn blood cells circulating in the brain, as they exclusively exist in the EdU+ nuclei. Of note, the cell-type-specific distribution of newborn cells was highly correlated between *TrackerSci* transcriptome and chromatin accessibility datasets (Spearman’s correlation r = 0.92; **Figure 3B**) and across conditions (**Figure S6**).

**Figure 2.**
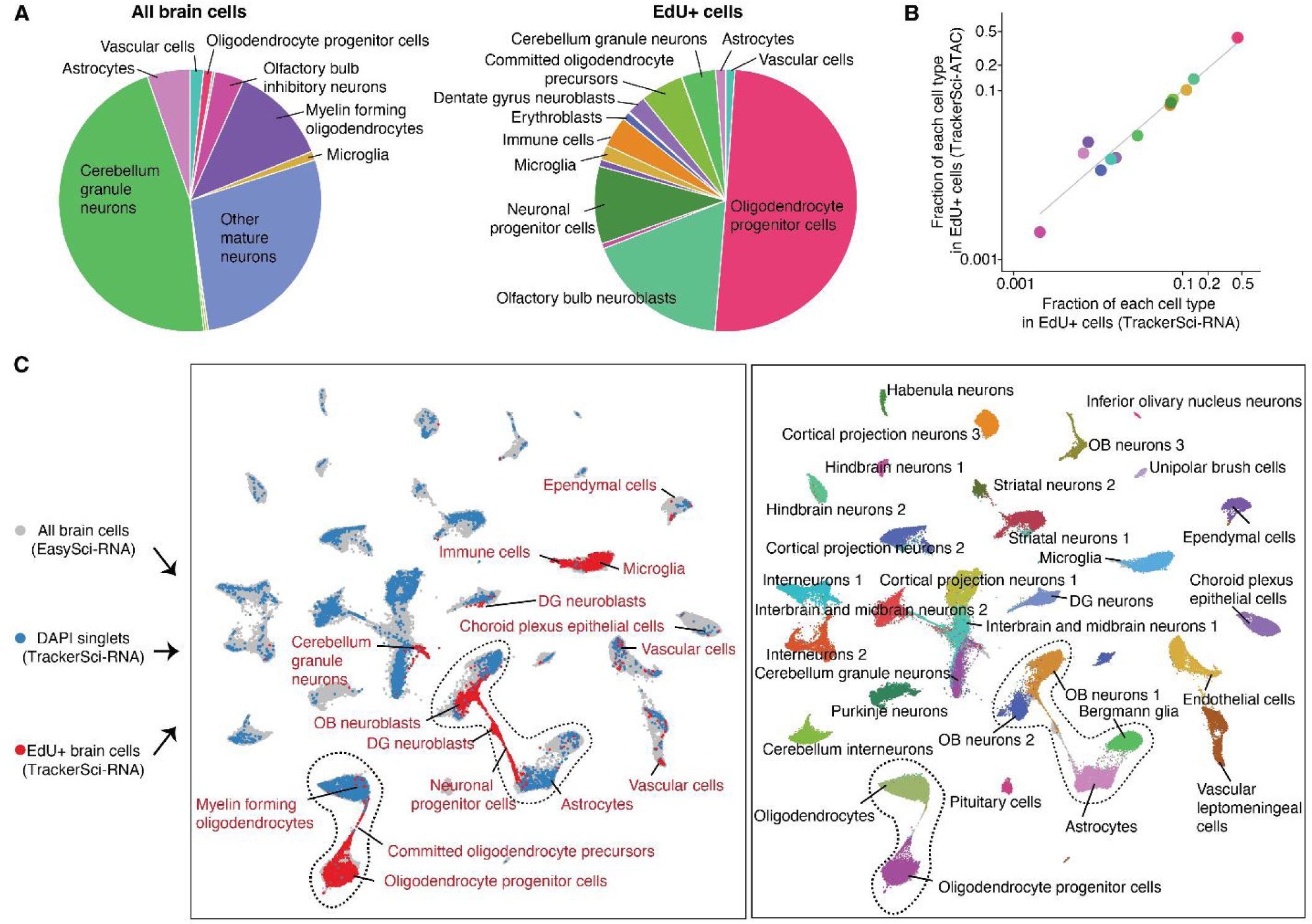
*TrackerSci* captures rare newborn cells that are less represented in conventional single-cell studies. (A) Pie plots showing the proportion of main cell types identified in the global cell population (left) and the enriched EdU+ cell population (right). (B) Scatter plot showing the fraction of each cell type in the enriched EdU+ cell population by single-cell transcriptome (x-axis) or chromatin accessibility analysis (y-axis) in *TrackerSci*, together with a linear regression line. (C) We integrated the *TrackerSci* dataset, including both EdU+ cells and DAPI singlets, with a large-scale brain cell atlas (Sziraki et al., 2022) comprising 1,469,111 cells. For the brain cell atlas, we sampled 5,000 cells of each cell type for the integration analysis. The UMAP plots show the integrated cells, colored by assay types (left, cell types from *TrackerSci* are annotated) or cell annotations from the brain cell atlas (right, cells from *TrackerSci* are colored in grey).

**Figure 3.**
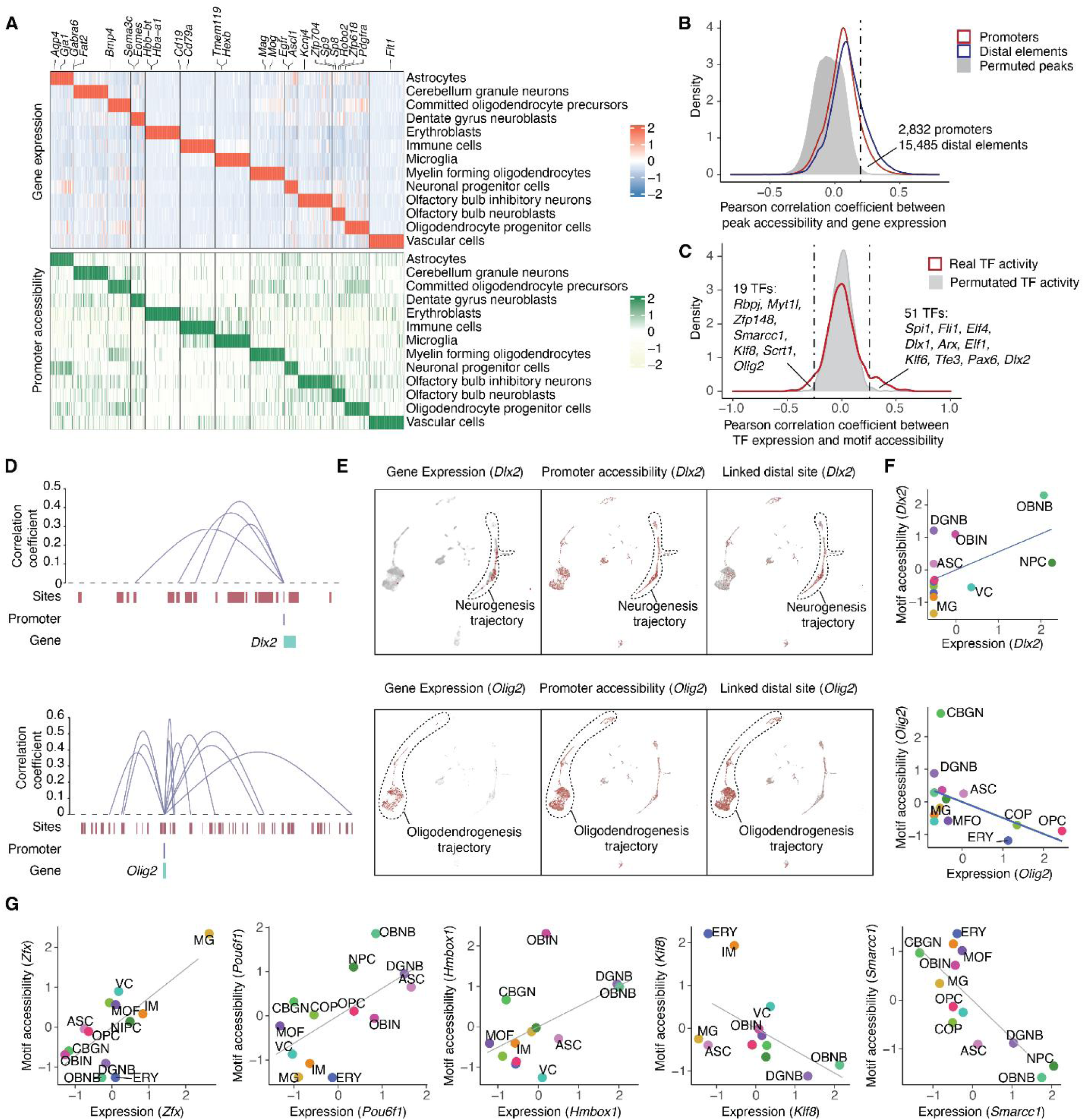
Identifying epigenetic elements and transcription factors associated with heterogeneous cellular states of newborn cells in the mouse brain. (A) Heatmap showing the relative expression (top) and chromatin accessibility (bottom) of cell-type-specific genes across cell types. The UMI count matrix (gene expression) and read count matrix (ATAC-seq) were normalized by the library size and then log-transformed, column centered, and scaled. The resulting values clamped to [-2, 2]. (B) Density plot showing the distribution of Pearson correlation coefficients between gene expression and the accessibility of promoter (colored in red) or nearby accessible elements (within ±500 kb of the promoter, colored in blue) across pseudo-cells. In addition, we plotted the background distribution of the Pearson correlation coefficient after permuting the accessibility of peaks across pseudo-cells. (C) Density plot showing the distribution of Pearson correlation coefficients between TF expression and their motif accessibility across pseudo-cells. The background distribution was calculated after permuting the motif accessibility of TFs across pseudo-cells. (D) Genome browser plot showing links between distal regulatory sites and genes for a neurogenesis marker (*Dlx2*, top) and an oligodendrogenesis marker (*Olig2*, bottom). (E) UMAP plots showing the cell-type-specific expression (left), the accessibility of promoter (middle), and linked distal site (right) for genes *Dlx2* (top) and *Olig2* (bottom). The single-cell expression data (UMI count) and ATAC-seq data (read count) were normalized first by library size and then log-transformed, column centered, and scaled. (F) Scatter plots showing the correlation between the scaled gene expression and motif accessibility across cell types for *Dlx2* (top) and *Olig2* (bottom), together with a linear regression line. ASC: astrocytes, CBGN: cerebellum granule neurons, COP: committed oligodendrocyte precursors, DGNB: dentate gyrus neuroblasts, ERY: erythroblasts, MFO: myelin-forming oligodendrocytes, MG: microglia, NPC: neuronal progenitor cells, OBNB: olfactory bulb neuroblasts, OBIN: olfactory bulb inhibitory neurons, OPC: oligodendrocyte progenitor cells, VC: vascular cells. (G) Scatter plots showing the correlation between the scaled gene expression and motif accessibility of less-characterized TF regulators, together with a linear regression line.

We next integrated *TrackerSci* datasets with a global brain cell atlas from our companion study (Sziraki et al., 2022), for which we profiled 1.5 million cells from entire mouse brains spanning three age groups and two mutants associated with AD. Briefly, we integrated EdU+ brain cells (5,715 single-cell transcriptomes from *TrackerSci*), ‘All’ brain cells (8,380 DAPI singlets from *TrackerSci*), and “All” brain cells from the global brain cell atlas (sampling 5,000 cells for each main cell type) into the same UMAP space. As expected, ‘All’ brain cells from the *TrackerSci* highly overlapped with cells from the global brain cell atlas in the integrated UMAP space (**Figure 2C**). Remarkably, the EdU+ cells (from *TrackerSci*) formed continuous cellular differentiation trajectories bridging several terminally differentiated cell types, including the oligodendrogenesis trajectory from the oligodendrocyte progenitor cells to differentiated oligodendrocytes, and the neurogenesis trajectory connecting astrocytes and OB neurons (**Figure 2C**). Of note, the bridge cells are validated by the expression of known progenitor markers, such as *Bmp4 and Enpp6* for committed oligodendrocyte precursors (Marques et al., 2018; Zhang et al., 2014) and *Mki67*, *Egfr* for neuronal progenitor cells (Pastrana et al., 2009) (**Figure S7A**). While the 1.5 million global brain cell atlas is one of the most extensive single-cell analyses of adult mouse brains, these “bridge” cells were still missing in the original trajectory analysis **(Figure S7B)**, highlighting the application of the *TrackerSci* method for recovering continuous cellular differentiation trajectories in adult tissues.

### Identify cell-type-specific epigenetic signatures and TF regulators of newborn cells

Toward a better understanding of the molecular signatures of newborn cells, we performed differential expression (DE) and differential accessibility (DA) analysis, yielding 5,610 DE genes (FDR of 5%, **Figure 3A**; **Table S3; Methods**) and 68,556 DA sites (FDR of 5%, **Table S4; Methods**) with significant changes across cell types. Notably, 1,744 (34.8%) of DE genes have DA promoters enriched in the same cell type (median Pearson r = 0.81, **Figure 3A**). While canonical gene markers were observed and used for our annotation of different cell types (**Figure S8**), we detected many novel markers that are highly cell-type-specific but have not been reported in prior research, including markers for neuronal progenitor cells (*e.g., Adgrv1* and *Rmi2*), DG neuroblasts (*e.g., Prdm8* and *Marchf4*), OB neuroblasts (*e.g., Zfp618* and *Sdk2*) and committed oligodendrocyte precursors (*e.g., Ccdc134* and *Mroh3*) (**Figure S8**). The cell type specificity of these markers were cross-validated by both gene expression and promoter accessibility. For comparison, some of the widely used neurogenesis markers, such as *Sox2* and *Dcx*, were found to be expressed across multiple cell types (*e.g.,* oligodendrocyte progenitor cells; **Figure S9**), which may affect their accuracy for labeling cells in neurogenesis (Hodge and Hevner, 2011).

To investigate the epigenetic landscape that shapes the transcriptome of newborn cells, we next sought to identify the cis-regulatory elements underlying the cell-type-specific expression of gene markers. We first computed the correlation between the expression of each gene marker and the accessibility of its nearby DA sites across 88 ‘pseudo-cells’ (a subset of cells with adjacent integrative UMAP coordinates grouped by k-means clustering, **Figure S10A; Methods**). To control for any potential artifacts of the analysis, we permuted the sample IDs of the data matrix followed by the same analysis pipeline. Altogether, we identified 15,485 positive links between genes and distal sites (plus 2,832 associations between genes and promoters) at an empirically defined significance threshold of FDR = 0.05 and based on their cell-type-specificity (**Figure 3B; Table S5; Methods**).

The identified distal site-gene linkages were significantly closer than all possible pairs tested (median 159 kb for identified links vs. 251 kb for all pairs tested; p-value < 5 × 10^−5^, unpaired permutation test based on 20,000 simulations, **Figure S10B**). Most genes were associated with a few links (median two distal sites per gene, out of a median of 94 distal sites within 500 kb of the TSS tested, **Figure S10B**). For example, *Dlx2*, a canonical neurogenesis marker (Petryniak et al., 2007), was significantly linked to four distal peaks, all exhibiting remarkable cell-type-specificity similar to its gene expression (**Figure 3D and 3E Figure S10C**). By contrast, a small subset of genes (3.5%) were linked with a large number of peaks (>= 10 peaks). For instance, *Olig2* was linked to 10 distal peaks (**Figure 3D**), all highly enriched in the oligodendrocyte progenitor cells (OPC) and committed oligodendrocyte precursors (COP) (**Figure 3E; Figure S10D**). Some genes (*e.g., Dlx2*) showed strong cell-type-specificity in their linked distal sites compared to their promoters (**Figure S10E**), indicating that long-range transcriptional control could play a key role in determining cell type specificities.

To further characterize transcription factors (TFs) that contribute to the cell type specification of progenitor cells, we computed the Pearson correlation coefficient between TF expression and motif accessibility across all afore-described “pseudo-cells”. We then performed the same analysis using the permuted data as the background control. At an empirically defined significance threshold of FDR = 0.05, we identified a total of 70 cell-type-specific TF regulators, including 19 potential repressors featured with negative correlations between gene expression and motif accessibility (*e.g., Olig2*, **Figure 3C and 3F**). Most cell-type-specific TFs are readily validated by previous studies. For example, *Olig2* has been reported to encode a transcriptional repressor during motor neuron differentiation and myelinogenesis (Zhang et al., 2022). Other examples include *Spi1* and *Runx1* in immune cells (Iwasaki and Akashi, 2007; Yeh and Ikezu, 2019); *Maf*, *Mef2a*, and *Tfe3* in microglia (Solé-Domènech et al., 2016; Yeh and Ikezu, 2019); and *Pax6, Nfib,* and *Arx* in neuronal progenitor cells and neuroblasts (Colombo et al., 2007; Ninkovic et al., 2013; Osumi et al., 2008). Notably, several less-characterized TFs were identified and validated by the cell-type-specific enrichment of both gene expression and motif accessibility, such as *Pou6f1*, *Hmbox1*, *Klf8,* and *Smarcc1* enriched in immature neurons and *Zfx* enriched in microglia, representing potentially regulators of progenitor cells in the adult brain (**Figure 3G; Figure S11**).

### A global view of cell-type-specific proliferation rates across the adult lifespan

We next compared the fraction of EdU+ cells across young, adult, and aged mice brains, and observed a marked reduction of cellular proliferation associated with age (**Figure 4A**). To investigate the cell-type-specific changes in proliferation rates, we then quantified the relative fractions of each newborn cell type by their fractions in the EdU+ cell population, multiplied by the ratio of EdU+ cells in the global cell population. Interestingly, we detected highly heterogeneous responses to aging across various progenitor cell types, validated by both single-cell transcriptome and chromatin accessibility profiles (**Figure 5B**). For example, dentate gyrus neuroblasts showed an 18-fold reduction in the aged brain (vs. the adult brain), while the proliferation of vascular cells were only mildly affected. In contrast, microglia and other immune cells showed a remarkable boost in the production of newborn cells (**Figure 4B-D**), possibly due to the elevated inflammatory signaling in the aged brain (Corlier et al., 2018). Compared with the aged brain, we detected overall mild changes in cellular proliferation (except the microglia) in the AD-associated mouse model (5xFAD), probably because the mutant mice were profiled at a relatively early stage (before three months).

**Figure 4.**
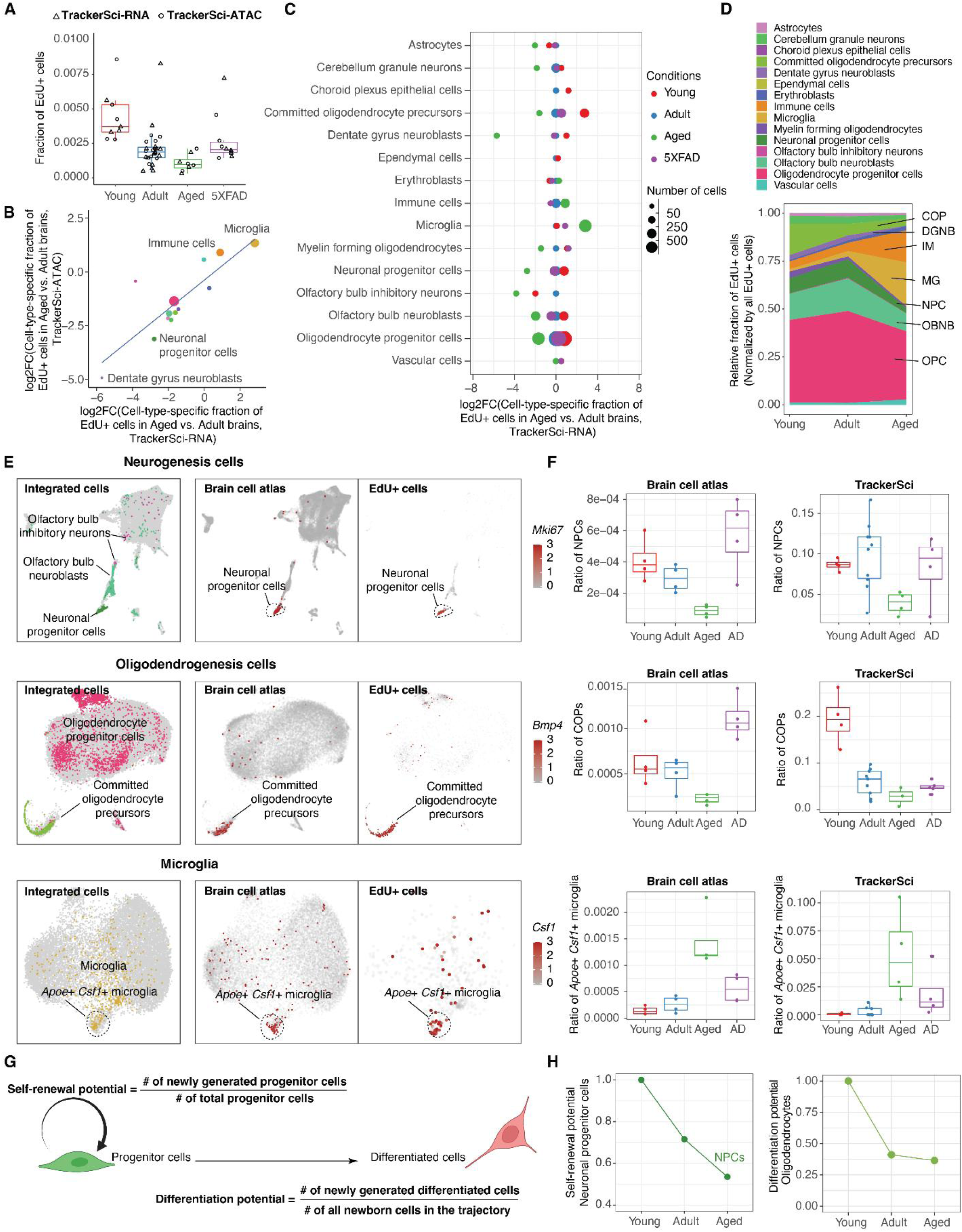
Deciphering the impact of aging on the proliferation status and differentiation dynamics of different cell types in the mammalian brain. (A) Boxplot showing the fraction of EdU+ cells in the mouse brain after five days of EdU labeling. The plot includes data from both single-cell transcriptome and chromatin accessibility experiments in *TrackerSci*. For all box plots in this figure: middle lines, medians; upper and lower box edges, first and third quartiles, respectively; whiskers, 1.5 times the interquartile range; and all individual data points are shown. (B) With the single-cell RNA-seq or ATAC-seq data of *TrackerSci*, we first calculated the cell-type-specific fractions in each condition (*i.e.,* young, adult, aged, and 5xFAD), multiplied by the fraction of EdU+ cells in the entire brain. We then quantified the fold changes of normalized cell-type-specific fractions between the aged and adult brains. The scatter plot shows the log-transformed fold changes (aged vs. adult) correlation between single-cell transcriptome and chromatin accessibility analysis in *TrackerSci*. (C) Similar to the analysis in (B), the dot plot shows the log-transformed cell-type-specific fold changes between each condition and the adult brain. For the comparison between 5xFAD and wild-type, we used mice of the same age (11-week-old) from both groups. (D) Area plot showing the cell-type-specific proportions in EdU+ cells over time. (E) We integrated cells corresponding to OB neurogenesis (top), oligodendrogenesis (middle), and microglia (bottom) in *TrackerSci* and brain cell atlas (Sziraki et al., 2022); the left UMAP plot shows the integrated cells, colored by cell type annotations in *TrackerSci* or grey (brain cell atlas). The two UMAP plots on the right show cells from the brain cell atlas or the EdU+ cells recovered by *TrackerSci*, colored by the expression of the neuronal progenitor marker *Mki67* (top), the committed oligodendrocyte precursor cells marker *Bmp4* (middle) and the aging/AD-associated microglia marker *Csf1* (bottom). (F) Box plots showing the cell-type-specific fractions of neuronal progenitor cells (top), committed oligodendrocyte precursors (middle) and aging/AD-associated microglia (bottom) across different conditions in the brain cell atlas (left) or newborn cells from *TrackerSci* (right). (G) Schematic showing the calculation of the self-renewal and differentiation potential of progenitor cells. (H) Left: Line plot showing the estimated self-renewal potential of neuronal progenitor cells over time. Right: Line plot showing the estimated differentiation potential of the newly generated oligodendrocyte progenitor cells across three age groups.

**Figure 5.**
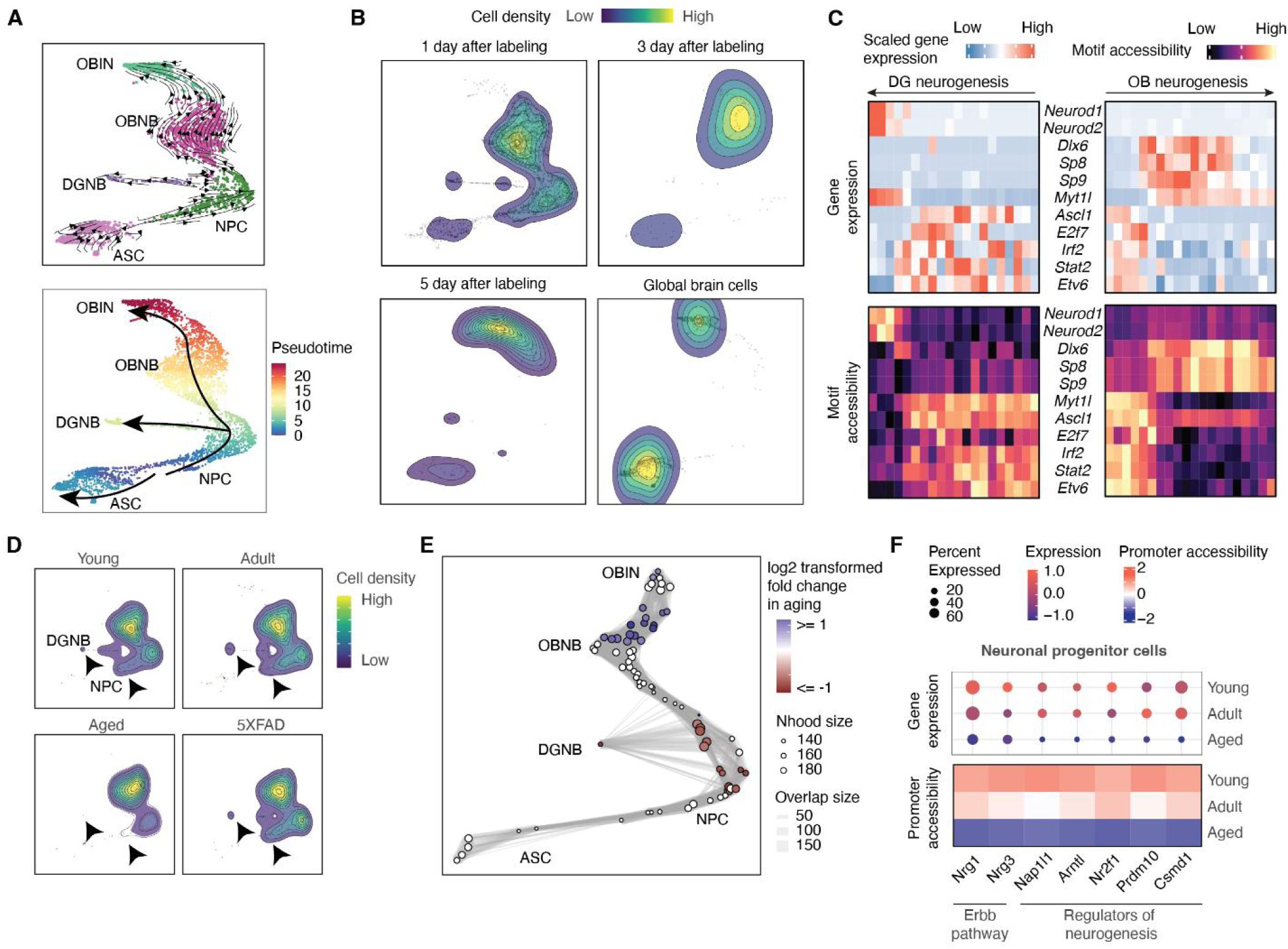
Characterizing the impact of aging on neurogenesis. (A) UMAP plots showing the differentiation trajectory of neurogenesis, colored by main cell types (top) or pseudotime (bottom). The differentiation trajectories are inferred by RNA velocity analysis (top) and annotated on the bottom plot. (B) Mice brains were harvested one day, three days and nine days after EdU labeling (EdU was administered daily through i.p. injection during the first five days), followed by *TrackerSci* profiling. The contour plots show the distribution of EdU+ cells in the neurogenesis trajectory across different harvest time points and the distribution of all brain cells without enrichment of EdU+ cells. (C) Heatmap showing the dynamics of gene expression and motif accessibility of cell-type-specific TFs across the pseudotime of neurogenesis trajectories. (D) Contour plots showing the distribution of EdU+ cells from *TrackerSci*-RNA in the neurogenesis trajectory across conditions. The arrows point to the significantly reduced cell states in each trajectory. (E) A neighborhood graph from Milo differential abundance analysis on the neurogenesis trajectory. The layout of the graph is determined by the position of the neighborhood index cell in (A). Nodes represent cellular neighborhoods from the KNN graph. Differential abundance neighborhoods are colored by the log-transformed fold change across ages. Graph edges depict the number of cells shared between neighborhoods. (F) The dot plots and heatmaps show the scaled gene expression and promoter accessibility of top differentially expressed genes in the neuronal progenitor cells.

To further validate the cell-type-specific dynamics in brain aging, we integrated the newborn cells recovered from *TrackerSci* and a global mice brain cell atlas (Sziraki et al., 2022) for sub-clustering analysis. Indeed, the integration analysis at the sub-cluster level facilitated the identification of rare progenitor cells in the global brain cell atlas, such as neuronal progenitor cells (marked by *Mki67*, *Top2a, and Egfr*) and committed oligodendrocyte precursors (marked by high expression of *Bmp4* and *Enpp6*) (**Figure 4E**). both of these cell types are remarkably reduced upon aging, validated in both datasets (**Figure 4F**). In addition, the integration analysis revealed a reactive microglia subtype, marked by high expression of *Apoe* and *Csf1* in both datasets. This microglia subtype has been previously reported to be enriched in aged and AD mammalian brains (Keren-Shaul et al., 2017). Consistent with prior studies, we found the proliferation rate of the *Apoe+, Csf1+* microglia increased significantly in both aged (p-value = 0.0045, Wilcoxon rank-sum test) and 5xFAD brains (p-value = 0.028, Wilcoxon rank-sum test), which readily explained its rapid expansion in both aged and disease conditions (**Figure 4F**).

We next sought to investigate the impact of aging on the self-renewal and differential potential of progenitor cells *in vivo*. We first defined the self-renewal potential by the number of newly generated progenitor cells divided by the number of total progenitor cells in the brain (*i.e.,* the number of new cells generated per progenitor cell in a fixed time, **Figure 4G**). For instance, the neuronal progenitor cells exhibited down-regulated self-renewal potential over ages (**Figure 4H**), which readily explained the depleted neural stem cell pool in the aged brain. Meanwhile, the differentiation potential of a cell type can be defined by the fraction of newly generated differentiated cells divided by all newborn cells in the same lineage (**Figure 4G**). For example, we observed a substantially reduced differentiation potential in oligodendrocyte progenitor cells across the adult lifespan, especially during the early growth stage (**Figure 4H**). This analysis represents a unique application of *TrackerSci* for quantitative measurement of cell-type-specific self-renewal and differentiation capacities *in vivo*.

### The impact of aging on adult neurogenesis

Adult neurogenesis and oligodendrogenesis have been reported to decline upon aging (Galvan and Jin, 2007; Pollina and Brunet, 2011); however, the detailed gene regulatory mechanism is still unclear due to technical limitations. We next sought to interrogate the impact of aging on adult neurogenesis and oligodendrogenesis, and delineate underlying transcriptional and epigenetic controls.

For adult neurogenesis, we identified three main trajectories that differentiated into DG neuroblasts, OB neuroblasts, and astrocytes, consistent with the cell state transition directions inferred by the RNA velocity analysis (Bergen et al., 2020) and prior report (Ratz et al., 2022) (**Figure 5A**). The trajectory was further validated through a pulse-chase experiment, where we harvested cells for *TrackerSci* profiling at different time points (*i.e.,* one day, three days, and nine days post-labeling). Indeed, we observed a gradual accumulation of more differentiated cell states with longer chasing time (**Figure 5B**). Through DE gene analysis, we identified 2,072 and 6,473 DE genes along the DG neurogenesis and OB neurogenesis trajectories, respectively **(Table S7 and S8)**. Of all DE genes, 1,799 genes were shared between the two trajectories, including up-regulated genes (*e.g., Dcx*) enriched in neuron development (q-value = 2.7e-8) (Chen et al., 2013) and down-regulated genes (*e.g., Notum*) enriched in negative Wnt signaling regulation (q-value = 0.0004) (Chen et al., 2013) **(Figure S12A)**. In addition, putative trajectory- and region-specific neurogenesis programs were identified, such as *Neurod1*, *Neurod2,* and *Emx1 enriched* in the DG trajectory **(Figure S12B)**. This is consistent with previous reports about their important roles in hippocampal neurogenesis (Brulet et al., 2017; Hong et al., 2007; Micheli et al., 2017).

With the chromatin accessibility profiling, we identified 3,095 and 13,790 sites showing dynamics patterns along the DG neurogenesis and OB neurogenesis trajectories, respectively **(Table S9 and S10)**, from which we further identified 20 TFs exhibiting significantly changed motif accessibility in the DG neurogenesis trajectory (FDR of 0.05, **Table S11**) and 318 TFs in OB neurogenesis (FDR of 0.05, **Table S12**). Key TFs were further validated by strong correlations between their expression and motif accessibility dynamics (**Figure 5C**). For example, the expression of the above-mentioned neurogenesis regulators, *Neurod1* and *Neurod2*, are positively correlated with their motif accessibility. In contrast, *Myt1l*, a known repressor of neural differentiation (Mall et al., 2017), shows a negatively correlated gene expression and motif accessibility. Leveraging this approach, we identified TFs shared between two neurogenesis trajectories (*e.g., Myt1l*, *Ascl1,* and *E2f7)*; as well as TFs that regulate the specification of different neuron types (*e.g., Dlx6, Sp8, Sp9* uniquely enriched in OB neurogenesis (Díaz-Guerra et al., 2013; Li et al., 2018a)). Meanwhile, we identified several TFs (*e.g., Irf2*, *Stat2,* and *Etv6*) showing strong enrichment of gene expression and motif accessibility in neuronal progenitor cells. While their functions in neurogenesis were less-characterized, some of them have been reported as essential regulators of other stem cell types, such as colonic stem cells (*Irf2*) (Minamide et al., 2020), mesenchymal stem cells (*Stat2*) (Yi et al., 2012), and hematopoietic stem cells (*Etv6*) (Hock et al., 2004; Yi et al., 2012).

To investigate the impact of aging on adult neurogenesis, we next compared the cellular density recovered from *TrackerSci* transcriptome profiling across different conditions along the neurogenesis trajectory. Consistent with the cell type level analysis (**Figure 4C**), we observed a dramatic age-dependent reduction in the cellular density of neural progenitor cells (NPC) and DG neuroblasts (DGNB), but not in OB neuroblasts (**Figure 5D**). The finding was consistent with the chromatin accessibility profiles, where we applied a recently published differential abundance testing algorithm, *Milo (Dann et al., 2021)*, to identify the cellular neighborhoods that are significantly altered upon aging. Thirty-one differentially decreased cellular neighborhoods were identified (**Figure 5E**, 5% FDR), mostly from the neural progenitor cells (NPC) and DG neuroblasts (DGNB). This analysis further validated that aging affects neurogenesis by down-regulating the proliferation rate of its progenitor cells.

To further decipher the molecular mechanisms underlying the age-dependent changes in neuronal progenitor cells, we then performed differential gene expression analysis across young, adult, and aged conditions, yielding thirty genes showing concordant changes over time, supported by both gene expression and the accessibility of promoters or linked distal sites (**Figure 5F; Table S13; Methods**). For example, two neurotrophic factors involved in the Erbb pathway, *Nrg1* and *Nrg3*, exhibited strongly reduced expression and promoter accessibility upon aging. Indeed, they have been reported to maintain neurogenesis upon *in vivo* administration (Mahar et al., 2016). In addition, we identified several other known regulators of neurogenesis, such as *Nr2f1* and *Nap1l1* (Bertacchi et al., 2020; Qiao et al., 2018), that were significantly down-regulated upon aging, which serve as potential targets for restoring adult neurogenesis in aged brains.

### The impact of aging on adult oligodendrogenesis

We next *in silico* isolated cell types that span multiple stages of oligodendrogenesis for pseudotime analysis, yielding a simple trajectory defined by integrated transcriptome and chromatin accessibility profiles (**Figure 6A**). The oligodendrogenesis trajectory was further validated by the RNA velocity analysis and the time-dependent labeling experiment mentioned above (**Figure 6B**). Through differential expression (DE) and differential accessibility (DA) analysis, we identified 8,443 DE genes and 15,164 DA sites that were significantly changed along the trajectory (5% FDR, **Table S14**). This analysis identified known oligodendrogenesis regulators (*e.g., Zfp276 (Aberle et al., 2022)* and *Myrf (Aberle et al., 2022; Fletcher et al., 2021)*) and associated pathways (*e.g.*, cholesterol biosynthesis (Mathews and Appel, 2016)), as well as novel gene markers (e.g., *Snx10*, *Rfbox2,* and *Tenm2*, **(Figure S12C)** with highly correlated changes of both molecular layers (i.e., RNA and promoter accessibility) along the trajectory of oligodendrogenesis.

**Figure 6.**
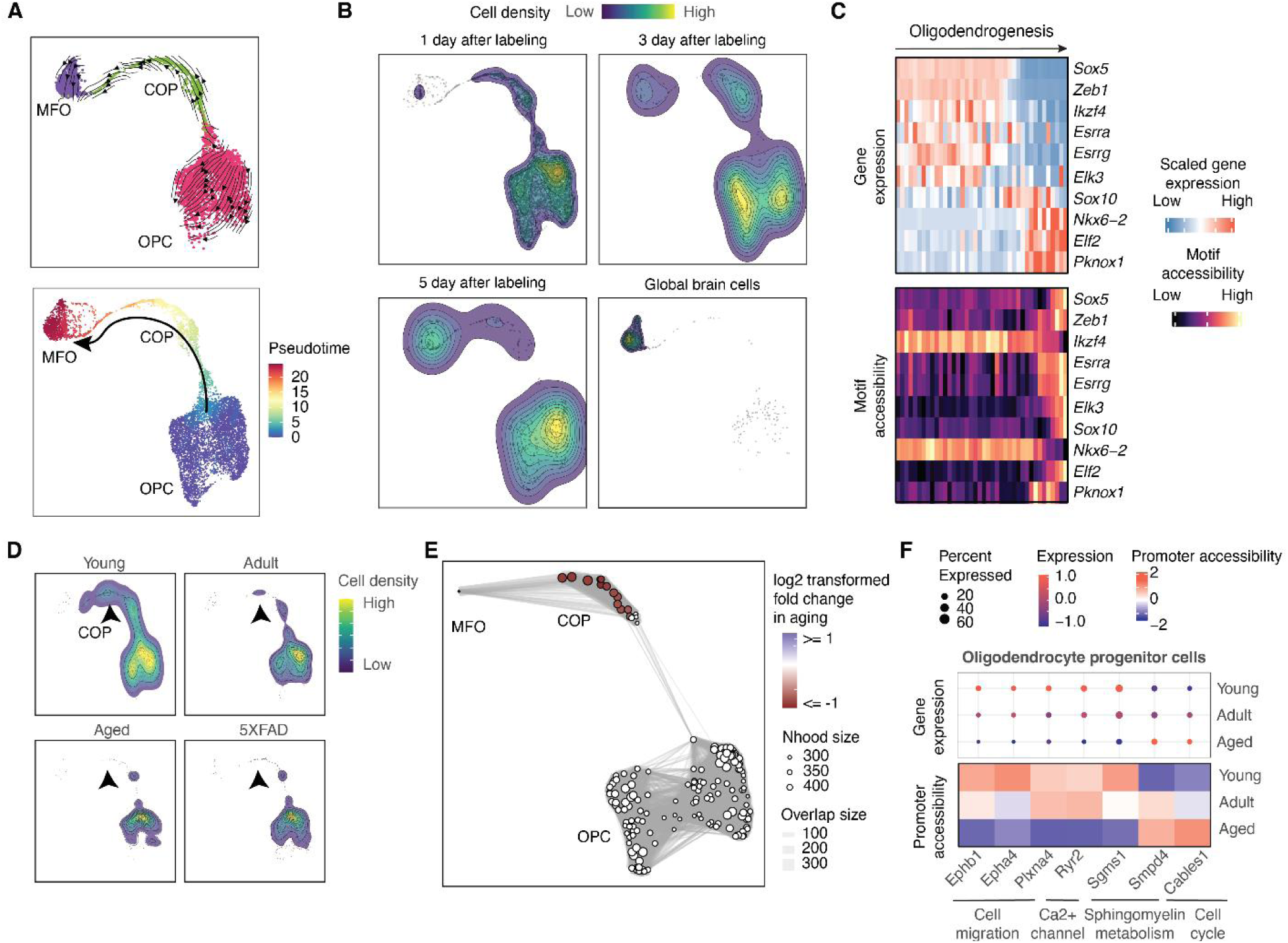
Characterizing the impact of aging on oligodendrogenesis. (A) UMAP plots showing the differentiation trajectory of oligodendrogenesis, colored by main cell types (top) or pseudotime (bottom). The differentiation trajectories are inferred by RNA velocity analysis (top) and annotated on the bottom plot. (B) Mice brains were harvested one day, three days and nine days after EdU labeling (EdU was administered daily through i.p. injection during the first five days), followed by *TrackerSci* profiling. The contour plots show the distribution of EdU+ cells in the oligodendrogenesis trajectory across different harvest time points and the distribution of all brain cells without enrichment of EdU+ cells. (C) Heatmap showing the dynamics of gene expression and motif accessibility of cell-type-specific TFs across the pseudotime of the oligodendrogenesis trajectory. (D) Contour plots showing the distribution of EdU+ cells from *TrackerSci*-RNA in the oligodendrogenesis trajectory across conditions. The arrows point to the significantly reduced cell states in each trajectory. (E) A neighborhood graph from Milo differential abundance analysis on the oligodendrogenesis trajectory. The layout of the graph is determined by the position of the neighborhood index cell in (A). Nodes represent cellular neighborhoods from the KNN graph. Differential abundance neighborhoods are colored by the log-transformed fold change across ages. Graph edges depict the number of cells shared between neighborhoods. (F) The dot plots and heatmaps show the scaled gene expression and promoter accessibility of top differentially expressed genes in the oligodendrocyte progenitor cells.

Moreover, we identified 97 TFs that exhibited highly correlated gene expression and motif accessibility in oligodendrogenesis (FDR of 5%, **Table S15 and S16**), including known regulators of oligodendrocyte differentiation, such as *Sox5*, *Sox10, Pknox1,* and *Nkx6-2 (Emery and Lu, 2015; Kato et al., 2015)*. In addition, several less-characterized TF markers were recovered, including *Ikzf4*, a known regulator of Müller glia differentiation in the retina (Javed et al., 2021), and several potential transcriptional ’repressors’ (*e.g., Esrra*, *Esrrg*, *Elk3*, *Zeb1*) characterized by the negative correlation between their expression and motif accessibility along the trajectory of oligodendrogenesis (**Figure 6C**).

We further investigated the impact of aging on adult oligodendrogenesis by examining cellular density along the cellular differentiation trajectory across different conditions. Unlike adult neurogenesis, we observed a remarkable reduction in committed oligodendrocyte precursors (COPs) rather than the early progenitor cells in single-cell transcriptome analysis (**Figure 6D**). The result is further validated through the *Milo (Dann et al., 2021)* analysis of chromatin accessibility profiles, where significantly decreased cellular neighborhoods exclusively overlapped with the committed oligodendrocyte precursors (COPs) (**Figure 6E**, 5% FDR). This observation is in accordance with the aging-associated depletion of newly formed oligodendrocytes in our companion study (Sziraki et al., 2022) and previous reports (Givre, 2003).

Finally, to delineate the molecular programs contributing to down-regulated oligodendrogenesis upon aging, we examined the significantly dysregulated genes in OPCs and identified 242 DE genes (FDR of 10%, **Table S17**). Many of the top DE genes are cross-validated by two independent molecular layers (*i.e.,* both gene expression and promoter accessibility) (**Figure 6F**). A lot of these genes are involved in molecular processes critical for oligodendrocyte differentiations, such as cell cycle (*e.g., Cables1 (He et al., 2019)*) or cell migration pathway (*e.g., Ephb1, Epha4, Plxna4*) (Linneberg et al., 2015; Smith et al., 1997) (**Figure 6F**). For example, we detected age-dependent down-regulation of *Ryr2*, a ryanodine receptor that mediates endoplasmic reticulum Ca^2+^ release, a process essential for initiating OPC differentiation (Li et al., 2018b). Intriguingly, two sphingomyelin metabolism-related genes exhibited opposite dynamics between young and aged OPCs (**Figure 6F**): *Sgms1*, a gene encoding a sphingomyelin synthase critical for converting phosphatidylcholine and ceramide to ceramide phosphocholine (sphingomyelin) and diacylglycerol at the Golgi apparatus (Huitema et al., 2004; Tafesse et al., 2007), was substantially down-regulated in the aged OPCs. By contrast, *Smpd4*, encoding a sphingomyelin phosphodiesterase that catalyzes the reverse reaction (Krut et al., 2006)(**Figure S13**), was significantly up-regulated in OPCs upon aging (**Figure 6F**). As a result, the age-dependent changes of both *Sgms1* and *Smpd4* could lead to the accumulation of ceramide and depletion of sphingomyelin in OPCs, which has been reported to increase cellular susceptibility to senescence and cell death (Hannun and Obeid, 2008; Jana et al., 2009). In fact, a recent report showed inhibiting another sphingomyelin hydrolase nSMase2 enhances the myelination and differentiation of OPCs (Yoo et al., 2020), suggesting a critical role of the dysregulated sphingomyelin metabolism in blocking oligodendrocyte differentiation in the aged brain. Furthermore, the down-regulated differentiation of oligodendrocytes is associated with dysregulated immune responses during aging, such as the accelerated proliferation of the reaction microglia subtype (**Figure 4F**) and an increased *C4b* expression in OPCs from both the EdU+ population and the global pool (**Figure S14**). Further investigation could be critical for deciphering the regulatory links between the elevated inflammation signaling and the dysregulation of oligodendrocyte differentiation in the aged brain.

### *TrackerSci* facilitates the identification of rare progenitor cells in the aged human brain

We next sought to investigate whether the *TrackerSci* dataset can be applied to facilitate the identification of rare progenitor cell types in the aged human brain. We first applied an extensively optimized single-cell RNA-seq by combinatorial indexing to profiling twenty-nine human brain samples derived from six individuals ranging from 70 to 94 in age at death (**Table S18**). Up to five regions (cerebellum, hippocampus, inferior parietal, motor cortex, and superior and middle temporal lobe (SMTG)) for each individual were included to characterize the region-specific effect of cellular dynamics. After removing low-signal cells and potential doublets, we recovered gene expression profiles in 798,434 single nuclei for downstream analysis (a median of 23,504 nuclei per brain sample, with a median of 1,013 UMIs per nucleus, **Figure S15A and S15B**)

Although this is one of the largest single-cell datasets of the aged human brain up to date, it was challenging to recover cycling or differentiating cells in the initial unsupervised clustering analysis (**Figure S15C**), potentially due to the extreme rarity of those cells in the aged human brain. We next integrated the *TrackerSci* dataset (including 5,715 EdU+ mouse brain cells and 8,380 mouse brain cells without EdU enrichment) with the human brain dataset followed by UMAP visualization (**Figure 7A**). Despite the species differences, the integration analysis facilitates the identification of extremely rare proliferating and differentiating cell populations in the aged human brain. For example, we identified a rare human cycling cell population that overlapped with cycling progenitor cells from mice (**Figure 7A**). Further sub-clustering analysis separated the population into three distinct subtypes (**Figure 7B**), corresponding to cycling microglia (569 cells, 0.07% of the total cell population, marked by *P2RY12* and *LY86*), cycling oligodendrocyte progenitor cells (56 cells, 0.007% of the total cell population, marked by *VCAN* and *PDGFRA*) and cycling erythroblasts (51 cells, 0.006% of the total cell population, marked by *CD36* and *KEL*). All of these clusters were marked by conventional proliferating markers such as *MKI67* and *TOP2A* (**Figure 7C**) and novel noncoding RNA markers such as *RP11-736I24.5, RP5-1086D14.6 and LINC01572* **(Figure S16A)**, demonstrating the application of *TrackerSci* as an anchor to capture extremely rare proliferating cells missed in the conventional single cell analysis. Interestingly, while the cycling microglia population expressed a common set of cell cycle-related genes (e.g., *MKI67*, *TOP2A*, *BUB1, SMC4*) and exhibited a similar ratio to the non-cycling microglia across brain regions (**Figure S16B**), we identified gene expression signatures unique to each region, suggesting a local control of microglia proliferation (**Figure S16C**). Of note, we detected very few neurogenesis cells in the aged human brains.

**Figure 7.**
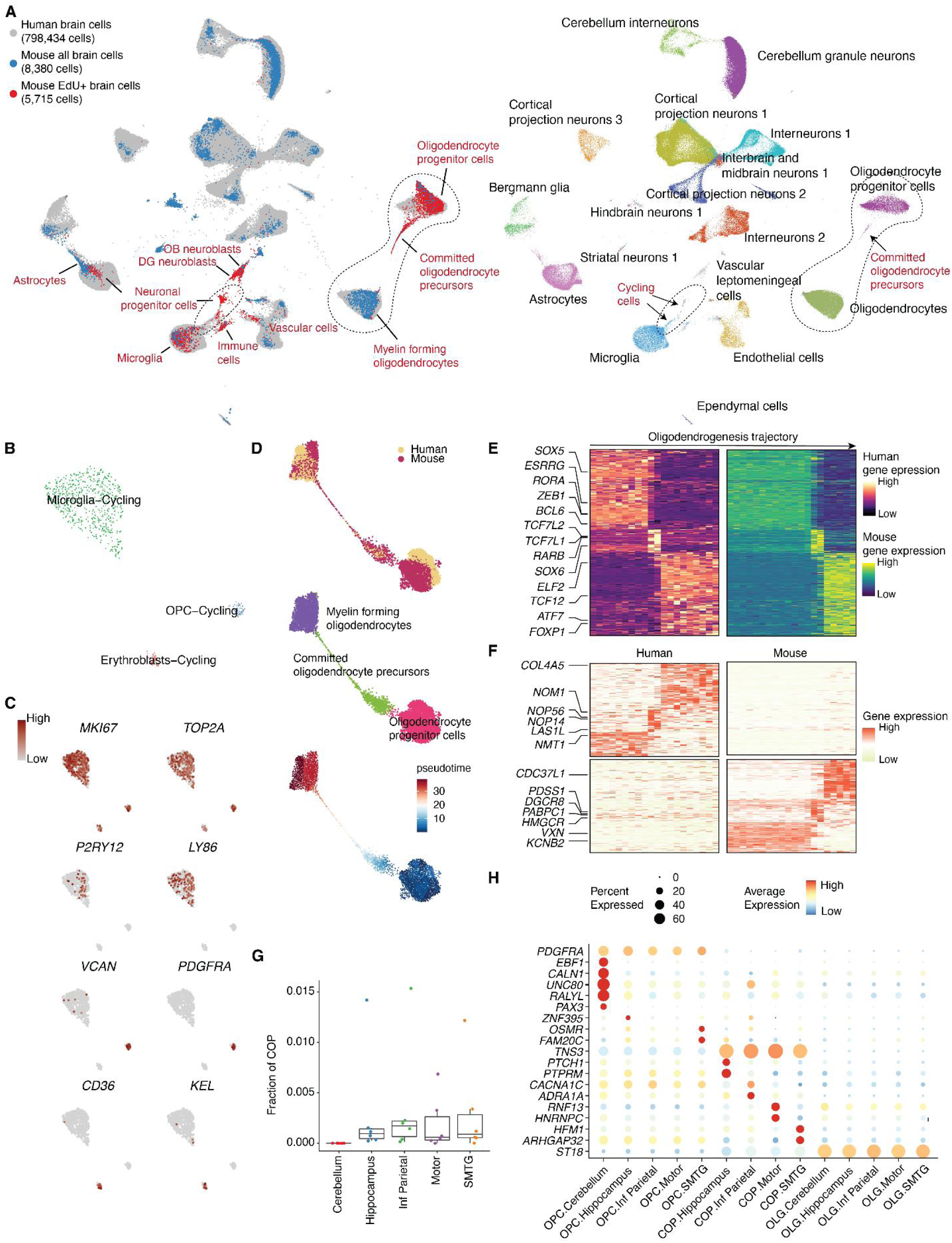
*TrackerSci* facilitates the identification of proliferating and differentiating cells in the human brain. (A) We integrated the *TrackerSci* dataset, including both EdU+ cells and DAPI singlets, with a large-scale human brain dataset comprising 798,434 cells. The UMAP plots show the integrated cells, colored by assay types (left, cell types from *TrackerSci* are annotated) or cell annotations from the human brain dataset (right, cells from *TrackerSci* are colored in grey). (B) UMAP plots showing the sub-clustering analysis of cycling cells from the human dataset, colored by cell annotation (B) and the expression level of markers for proliferation (*MKI67* and *TOP2A*; C), microglia (*P2RY12* and *LY86*; C), oligodendrocyte progenitor cells (*VCAN* and *PDGFRA*; C) and erythroblasts (*CD36* and *KEL*; C). (C) We integrated the oligodendrogenesis-related cells from *TrackerSci* and the human dataset. For the human brain dataset, we included all cells from committed oligodendrocyte precursors and randomly sampled 1,000 cells from oligodendrocyte progenitor cells and mature oligodendrocytes for the integration analysis. The UMAP plots show the resulting differentiation trajectory, colored by species (top), cell type annotations (middle) and pseudotime (bottom). (D) Heatmaps showing conserved gene expression dynamics along the oligodendrogenesis trajectory for human (left) and mouse (right), with key TF regulators annotated on the left. (E) Heatmaps showing divergent gene expression dynamics along the oligodendrogenesis trajectory enriched only in human (top) and mouse (bottom), with key genes annotated on the left. (F) Boxplot showing the fraction of committed oligodendrocyte precursors (COP) among oligodendrogenesis-related cells across different brain regions in each sample. For all box plots: middle lines, medians; upper and lower box edges, first and third quartiles, respectively; whiskers, 1.5 times the interquartile range; and all data points are shown. (G) Dotplot showing examples of commonly-changed and region-specific gene expression signatures across three differentiation stages along oligodendrogenesis trajectories.

Furthermore, integration analysis with the *TrackerSci* dataset facilitates the recovery of a stereotypical cell differentiation trajectory. For example, 188 committed oligodendrocyte precursors were identified in the aged human brain (0.02% of the total cell population), corresponding to the intermediate cells connecting the oligodendrocyte progenitor cells to mature oligodendrocytes (**Figure 7A**). To decipher the conserved gene dynamics underlying oligodendrogenesis between human and mouse, we extracted oligodendrogenesis-related cells from both species for integration analysis, yielding a smooth cell transition trajectory from progenitors to differentiated cell state (**Figure 7D**). We identified 5,680 genes that significantly changed along the human oligodendrogenesis trajectory (FDR of 5%), out of which 1,162 genes (48 TFs) were shared between human and mouse (**Figure 7E, Table S19**). While most of the conserved TFs have been previously reported as key regulators of oligodendrocyte differentiation (*e.g., TCF7L1* and *TCF7L2* (Weng et al., 2017)), several TFs have not been well characterized in the relevant context, such as *ZEB1*, *ESRRG, BCL6*, *RARB*. Notably, some less-characterized TFs were also nominated in our previous motif analysis (**Figure 6C**). In addition, we identified gene signatures that contribute to interspecies differences in oligodendrogenesis (**Figure 7F**). For example, the human-specific genes are enriched in ribosome biogenesis (e.g., *NOM1*, *NOP56*, *NOP14,* and *LAS1L*), while genes specifically linked to mouse oligodendrogenesis are involved in multiple pathways such as primary miRNA processing (e.g., *DGCR8* and *SRRT*), mRNA 3’-end processing (e.g., *PABPN1*, *SSU72,* and *PABPC1*) and isoprenoid biosynthetic processes (e.g., *PDSS1* and *HMGCR*).

Leveraging the dataset, we next investigated the differences in oligodendrogenesis across brain regions. Interestingly, we observed a depletion of the committed oligodendrocyte precursors in all cerebellum samples compared with other brain regions (**Figure 7G and Figure S17B**; p-value = 0.001, Fisher’s exact test), suggesting a reduced rate of oligodendrogenesis in the cerebellum. To gain more insight into the detailed molecular programs underlying the region-specific change of oligodendrogenesis, we performed DE analysis across regions and identified 45, 32, and 25 region-specific DE genes in OPC, COP, and OLG, respectively (**Table S20**). For example, region-specific gene signatures of COP were identified, such as *PTCH1* and *PTPRM* (hippocampus), *CACNA1C* and *ADRA1A* (inferior parietal), *RNF3* and *HNRNPC* (motor cortex), and *HFM1* and *ARHGAP32* (SMTG) (**Figure 7H**). Strikingly, 40 out of the 45 region-associated genes of OPC (*e.g., EBF1, PAX3, CALN1, and UNC30*) were highly enriched in the cerebellum (**Figure 7H**), indicating a unique molecular state of OPC in the cerebellum compared with other regions. Furthermore, one of the cerebellum-specific markers, *PAX3*, encodes a paired box transcription factor and has been reported to maintain the non-differentiating state of Schwann cells in the peripheral nervous system (Kioussi et al., 1995). This is consistent with our observation that the COP is depleted in the cerebellum. As a further illustration of this point, the cerebellum exhibited a higher fraction of OPCs accompanied by a decreased ratio of mature oligodendrocytes compared to other regions (**Figure S17A**). These analyses indicate a region-specific down-regulation of oligodendrogenesis in the cerebellum of the aged human brain.

## Discussion

The field of single-cell biology is progressing at a rapid rate to catalog and characterize each specific cell type across diverse biological systems. Although the adult or aged brains have been intensively profiled with single-cell methods (Li et al., 2021; Saunders et al., 2018; Zeisel et al., 2018), it has been challenging to capture rare progenitor cells and characterize their proliferation and differentiation potentials. Compared with prior studies (*e.g.,* Div-seq (Habib et al., 2016)), *TrackerSci* represents a unique approach to track both epigenetic and transcriptional dynamics of proliferating cells based on the strategy of combinatorial indexing. Like other sci-seq techniques (Cao et al., 2020; Domcke et al., 2020), *TrackerSci* is compatible with fresh or fixed nuclei, and can process multiple samples concurrently per experiment to reduce the batch effect. In this study, we applied *TrackerSci* to profile the single-cell transcriptome or chromatin accessibility dynamics for a total of 14,689 newborn cells from entire mouse brains spanning three age stages and two genotypes. Considering the rarity of the progenitor cells, especially in aged brains, it required deep sequencing of up to 15 million brain cells to recover the same amount of progenitor cells by conventional single-cell techniques.

Our analyses demonstrated unique advantages of *TrackerSci* over solely profiling global cell populations. For example, *TrackerSci* enabled us to reconstruct continuous cellular differentiation trajectories in adult or even aged organs by detecting intermediate progenitor cell states that are often missed in traditional single-cell analysis. Moreover, we were able to calculate the proliferation and differentiation potential of rare progenitor cells, facilitating the quantitative investigation of the impact of aging on adult neurogenesis and oligodendrogenesis. In addition, we further investigated age-dependent changes in cell-type-specific proliferation and differentiation dynamics and provided novel insights into the underlying transcriptional and epigenetic mechanisms.

There is a consensus that the self-renewal and regeneration capacity of progenitor cells reduces as we age. Through a comprehensive and quantitative view of the cell-type-specific proliferation and differentiation dynamics, however, we observed heterogeneous cellular responses to aging across progenitor cell types. While aging was associated with a depleted pool of neuronal progenitors as we expected, we found newborn oligodendrocyte progenitors were only mildly affected. Instead, the intermediate differentiation precursors were remarkably reduced especially at a relatively early stage (before six months), suggesting that aging affects oligodendrocytes mainly by blocking their differentiation process, consistent with the age-dependent downregulation of myelination in previous studies(Wang et al., 2020; Zhang et al., 2021). Intriguingly, we detected an age-dependent increase of *Smpd4* (sphingomyelin phosphodiesterase) and a decrease of *Sgms1* (sphingomyelin synthase) expression in the oligodendrocyte progenitor cells, suggesting that a high cellular ceramide level was associated with the aging-induced inhibition of oligodendrocyte differentiation.

To further investigate rare progenitor cell types in human brains, we generated a single-cell transcriptome atlas of human brains comprising almost 800,000 cells. While conventional clustering analysis failed to identify the rare progenitor cells in the dataset, integrative analysis with the *TrackerSci* dataset facilitated the identification of extremely rare cycling cells of microglia (0.07% of the total cell population) and OPCs (0.007% of the total cell population) in the aged human brain. The integration analysis enabled us to identify committed oligodendrocyte precursors (0.02% of the total cell population) across different brain regions, which confirmed the existence of oligodendrogenesis in the aged human brain. Further analysis of the data also nominated oligodendrogenesis-associated gene signatures that are shared or divergent between species. For example, we observed an increased expression of ribosome biogenesis factors in human oligodendrogenesis, while several genes involved in microRNA processing and mRNA polyadenylation are uniquely upregulated in mouse brains, suggesting a species-specific preference of regulation in global translation or transcription during oligodendrocyte differentiation. In addition, we recovered the differences of human oligodendrogenesis across anatomical locations, and identified molecular programs contributing to the down-regulated oligodendrogenesis in the aged human cerebellum.

In summary, the study represents a key step toward understanding the impact of aging on the proliferation and differentiation potential of progenitor cells in the mammalian brain. We anticipate that *TrackerSci* will be broadly used to identify and quantify cell-genesis processes across diverse systems, including other mammalian organs and humanized organoids. In addition, we envision similar strategies (*i.e.,* coupling the sci-seq platform with *in vivo* cellular labeling) can be expanded to study other critical molecular aspects, such as the cell-type specific survival, apoptosis, and senescent states. This will facilitate a comprehensive view of the global molecular programs regulating cell-type-specific dynamics during aging, thereby informing potential pathways to restore tissue homeostasis for patients with aging-related diseases.

### Endnotes

## Acknowledgments

We thank members of the Cao lab, especially Z. Zhang, Z. Xu, for helpful discussions and feedback. We are grateful to R. Satija (New York Genome Center) and J. Shendure (University of Washington) for insightful feedback related to this work. We also thank members from the Rockefeller University Flow Cytometry Resource Center and Comparative Bioscience Center for their help on FACS sorting experiments and animal maintenance.

## Author contributions

J.C. and W.Z.. conceptualized and supervised the project. Z.L. and M.Z. performed the EdU injection, mouse brain dissection, nuclei extraction, and fixation. M.Z. and Z.L developed and performed the *TrackerSci-RNA* experiments. Z.L developed and performed *TrackerSci-ATAC* experiments. S.A. and P.T.N. processed the human brain samples for single-cell profiling experiment. J.L. performed the *EasySci-RNA* experiments for the human dataset. Z.L. performed computational analyses with input from J.L. and A.S.. J.C., W.Z., and Z.L. wrote the manuscript with input and biological insight from M.Z., S.G., P.T.N. and other co-authors.

## Supplementary Figures

**Figure S1.**
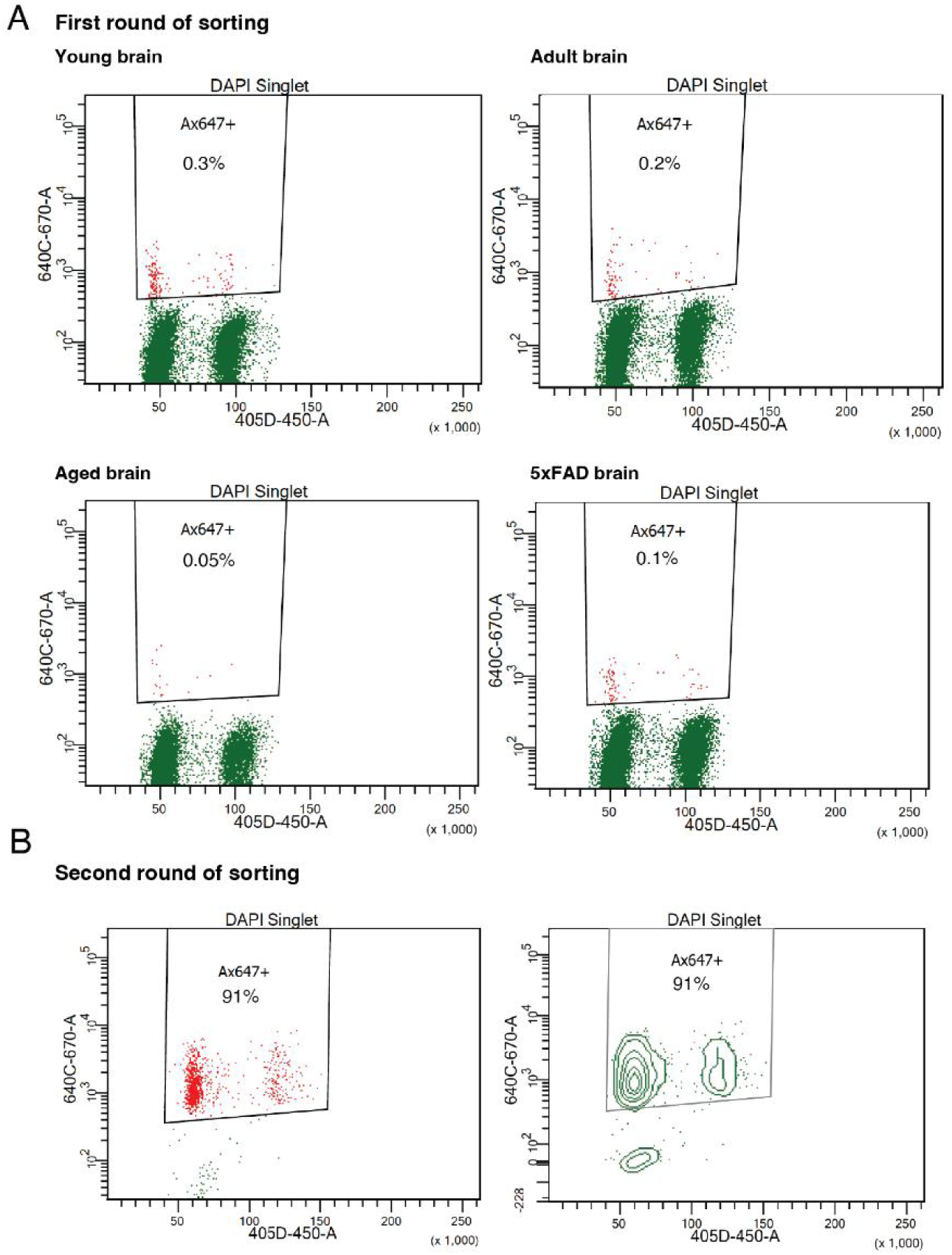
*TrackerSci* relies on two rounds of sorting to enrich and purify rare EdU+ proliferating cells in mammalian brains. (A) Representative Fluorescent-activated cell sorting (FACS) scatter plots showing the percentage of EdU+ cells in mouse brains across different conditions during the first round of sorting. (B) FACS scatter plot (left) and contour plot (right) showing the percentage of EdU+ cells during the second round of sorting in *TrackerSci*.

**Figure S2.**
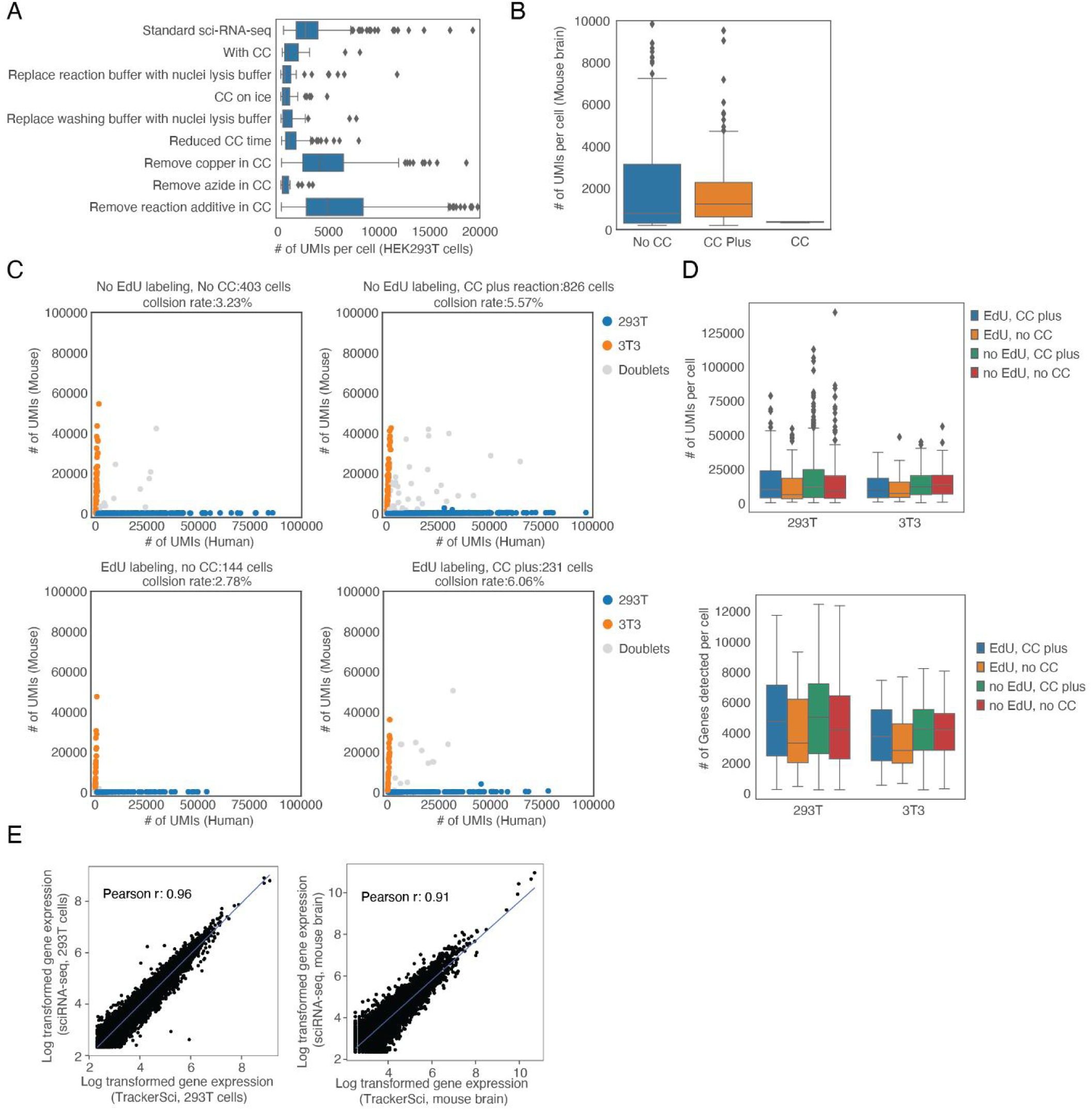
Quality control of *TrackerSci* for single-cell transcriptome profiling. (A) Boxplot showing the number of unique transcripts detected per cell (HEK293T nuclei) after different treatment conditions of click-chemistry (CC). The result indicated copper and reaction addictive in the conventional click-chemistry reaction decreased the scRNA-seq efficiency. For all box plots: middle lines, medians; upper and lower box edges, first and third quartiles, respectively; whiskers, 1.5 times the interquartile range; and diamonds are outliers. (B) Boxplot showing the number of unique transcripts detected per cell (mouse brain nuclei) across three conditions: no click-chemistry (No CC), conventional click-chemistry (CC), and click-chemistry plus condition (with picolyl azide dye and copper protectant, CC Plus). (C) Scatter plots showing the number of unique human and mouse transcripts detected per cell across different conditions (with/without EdU labeling, with/without click chemistry plus reaction). (D) Boxplot showing the number of unique transcripts (top) and genes (bottom) detected per cell in HEK293T and NIH/3T3 nuclei across the four conditions described in (C). (E) Scatter plot showing the correlation between log-transformed aggregated gene expression profiled by *TrackerSci* and sci-RNA-seq in HEK293T cells (left) and mouse brain cells (right), together with the linear regression line (blue).

**Figure S3.**
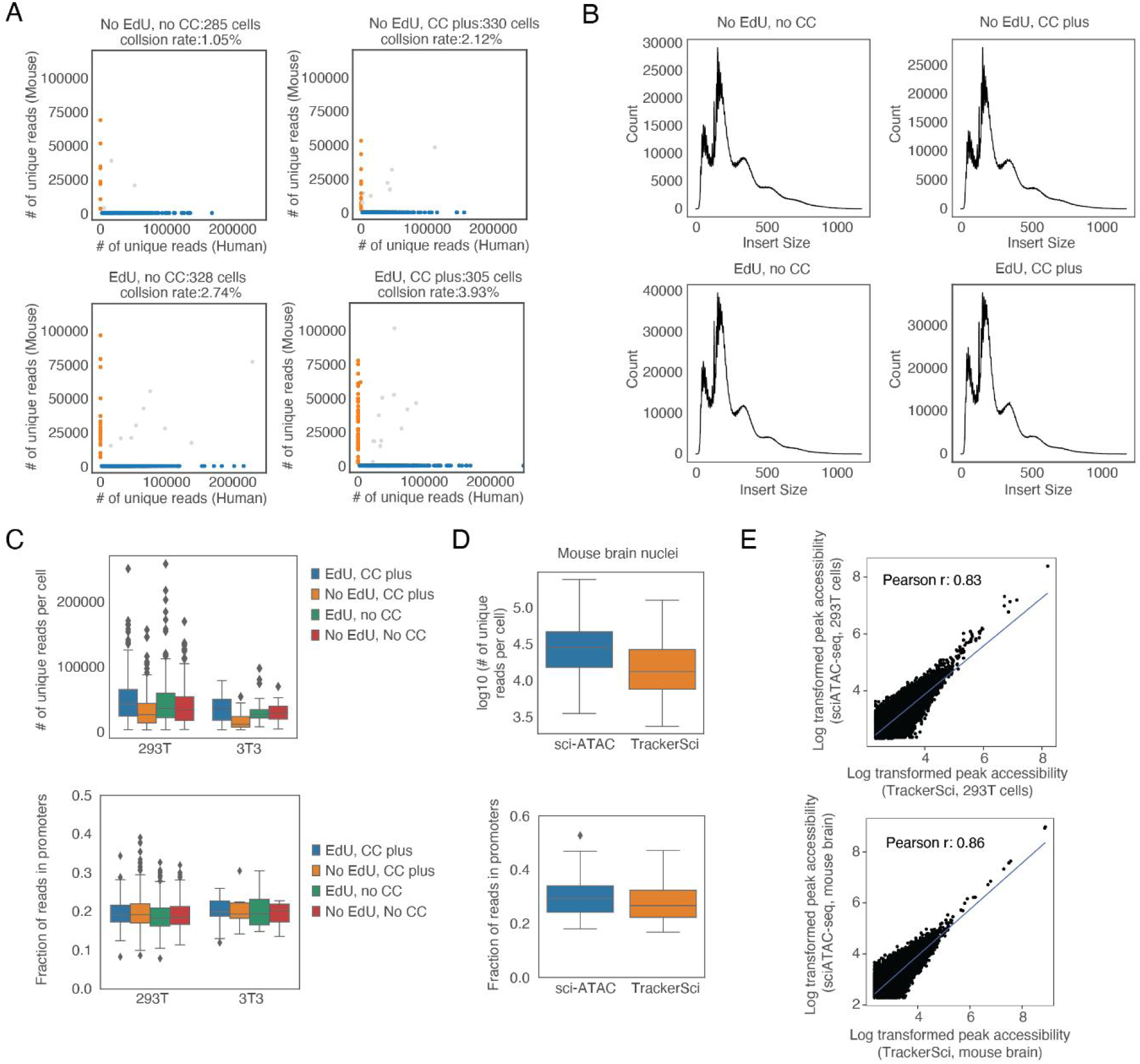
Quality control of *TrackerSci* for single-cell chromatin accessibility profiling. (A) Scatter plots showing the number of unique human and mouse ATAC-seq fragments detected per cell across different conditions (with/without EdU labeling, with/without click chemistry plus reaction). (B) The aggregated fragment length distribution in ATAC-seq from *TrackerSci* of all cells across the four conditions described in (A). No CC: no click-chemistry. CC plus: click-chemistry plus condition (with picolyl azide dye and copper protectant). (C-D) Boxplots showing the number of unique ATAC-seq reads (top) and the fraction of reads in promoters (bottom) in HEK293T and NIH/3T3 nuclei (C) and mouse brain nuclei (D). For all box plots: middle lines, medians; upper and lower box edges, first and third quartiles, respectively; whiskers, 1.5 times the interquartile range; and diamonds are outliers. (E) Scatter plot showing the correlation between log-transformed aggregated ATAC-seq peak accessibility (reads per million) profiled by *TrackerSci* and sci-ATAC-seq in HEK293T cells (top) and mouse brain cells (bottom), together with the linear regression line (blue).

**Figure S4.**
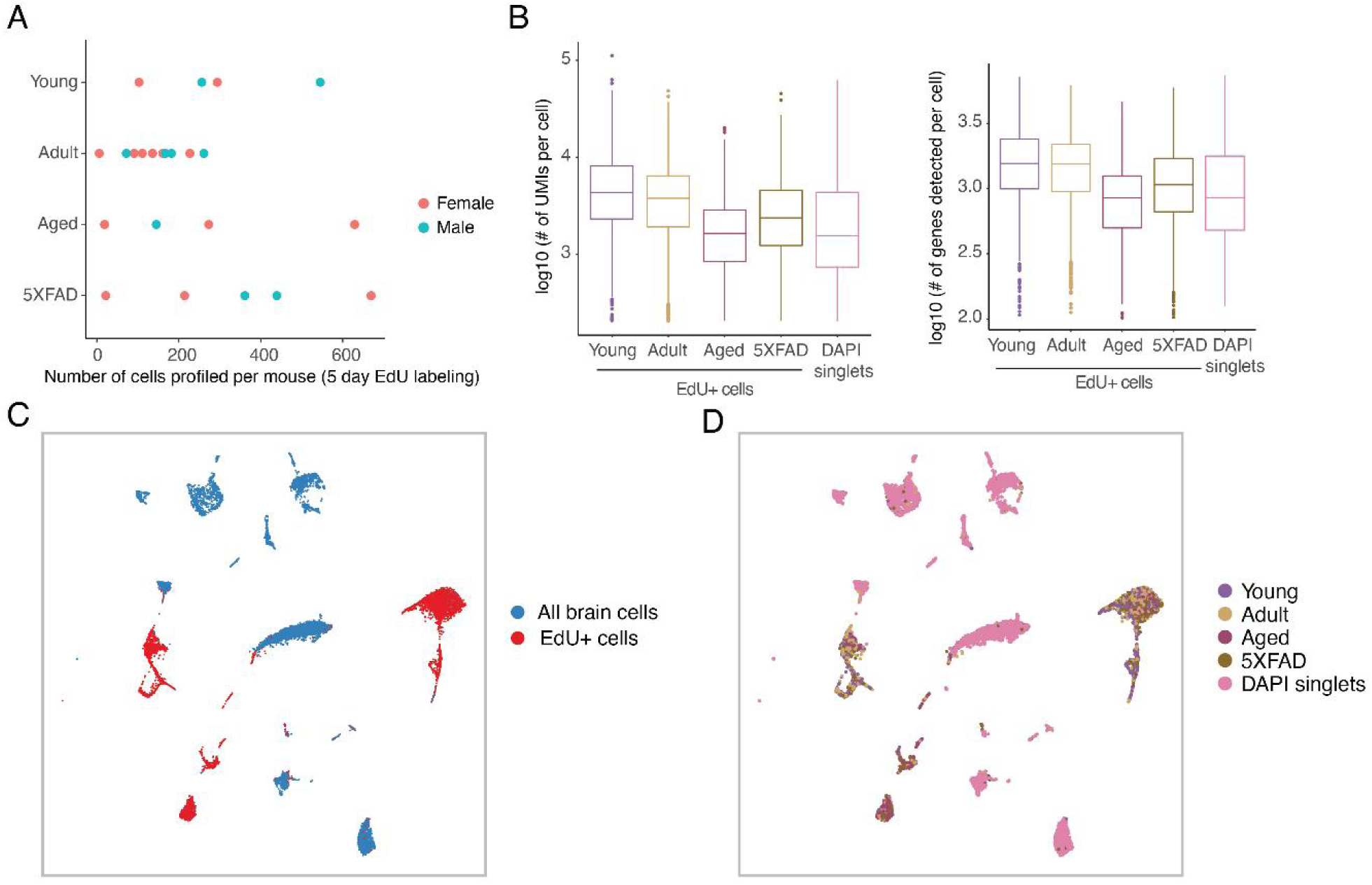
*TrackerSci* recovered single-cell transcriptomes of rare newborn cells in the mammalian brain. (A) Scatter plots showing the number of single-cell transcriptomes profiled in each mouse individual across four conditions, colored by sexes. Only mice from the main experiment group (EdU labeling for 5 days) are shown. (B) Boxplot showing the log-transformed number of unique transcripts (left) and genes (right) detected per cell profiled by *TrackerSci* and the DAPI singlet (without enrichment of EdU+ cells, adult mouse brain). For all box plots: middle lines, medians; upper and lower box edges, first and third quartiles, respectively; whiskers, 1.5 times the interquartile range; and circles are outliers. (C-D) UMAP visualization of single-cell transcriptomes, including EdU+ cells (profiled by *TrackerSci*) and all brain cells (without enrichment of EdU+ cells), colored by experiments (C) and conditions (D).

**Figure S5.**
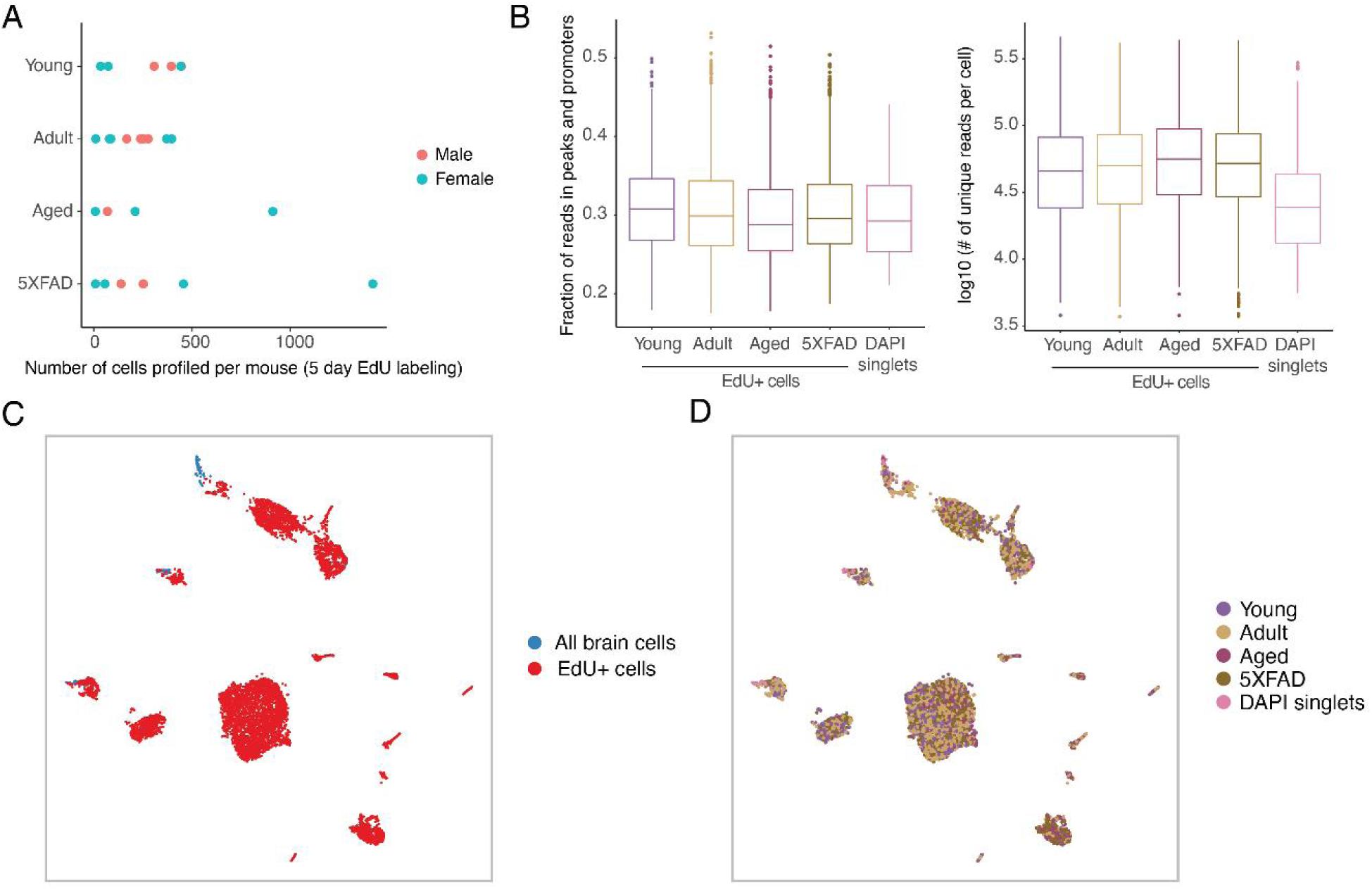
*TrackerSci* recovered single-cell chromatin accessibility of rare newborn cells in the mammalian brain. (A) Scatter plot showing the number of single-cell chromatin accessibility profiles in mouse individuals across four conditions, colored by sexes. Only mice from the main experiment group (EdU labeling for 5 days) are shown. (B) Boxplot showing the fraction of reads in promoters and peaks (left) and the log-transformed number of unique ATAC-seq reads (right) detected per cell across different conditions in *TrackerSci* and the DAPI singlet (adult mouse brain, without enrichment of EdU+ cells). For all box plots: middle lines, medians; upper and lower box edges, first and third quartiles, respectively; whiskers, 1.5 times the interquartile range; and circles are outliers. (C-D) UMAP visualization of single-cell chromatin accessibility profiles, including EdU+ cells (profiled by *TrackerSci)* and all brain cells (without enrichment of EdU+ cells), colored by experiments (C) and conditions (D)

**Figure S6.**
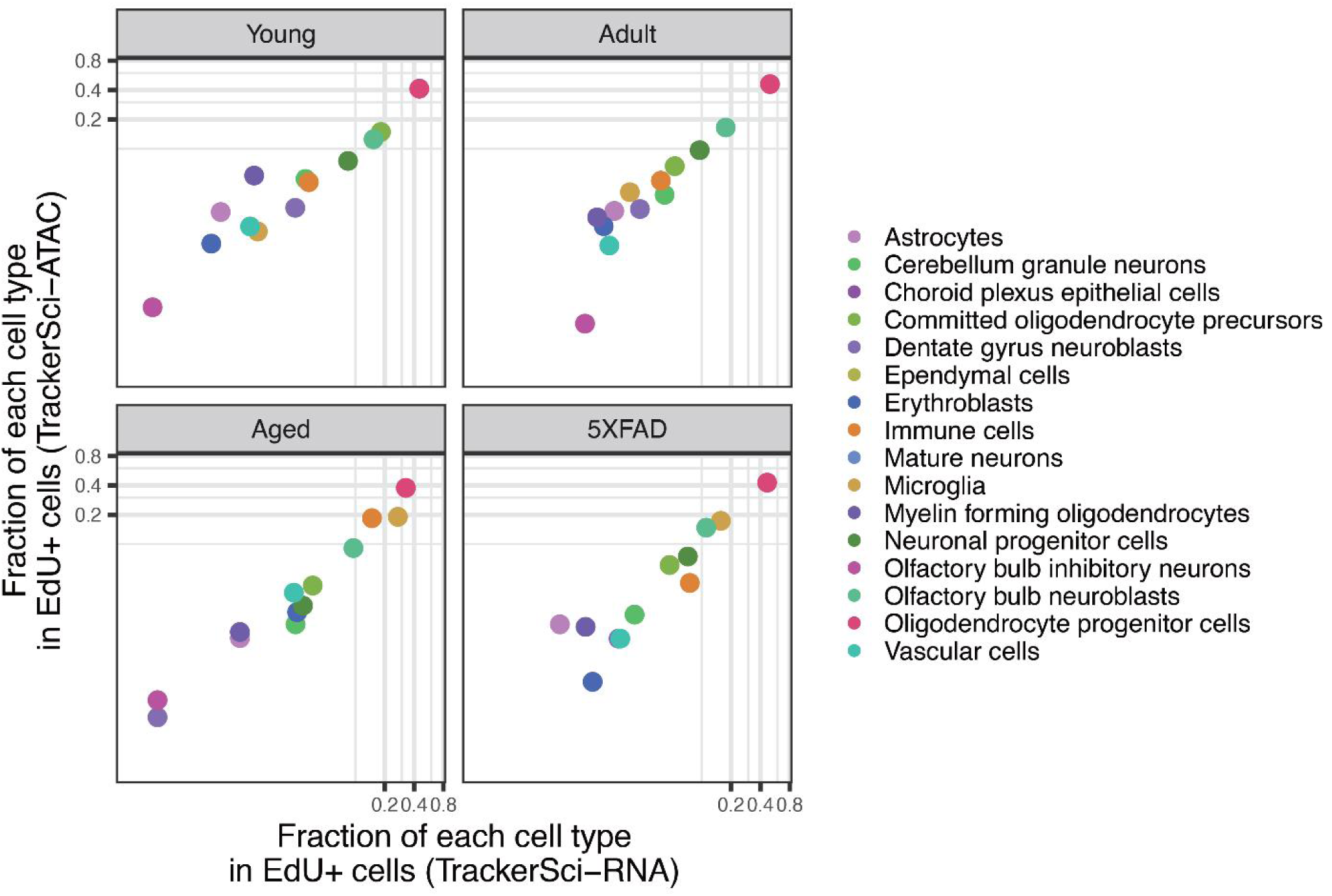
The cell population distributions are correlated between single-cell transcriptome and chromatin accessibility profiling of newborn cells in the mouse brain. Scatter plot showing the fraction of each cell type in the enriched EdU+ cell population by single-cell transcriptome (x-axis) or chromatin accessibility analysis (y-axis) in *TrackerSci* across different conditions.

**Figure S7.**
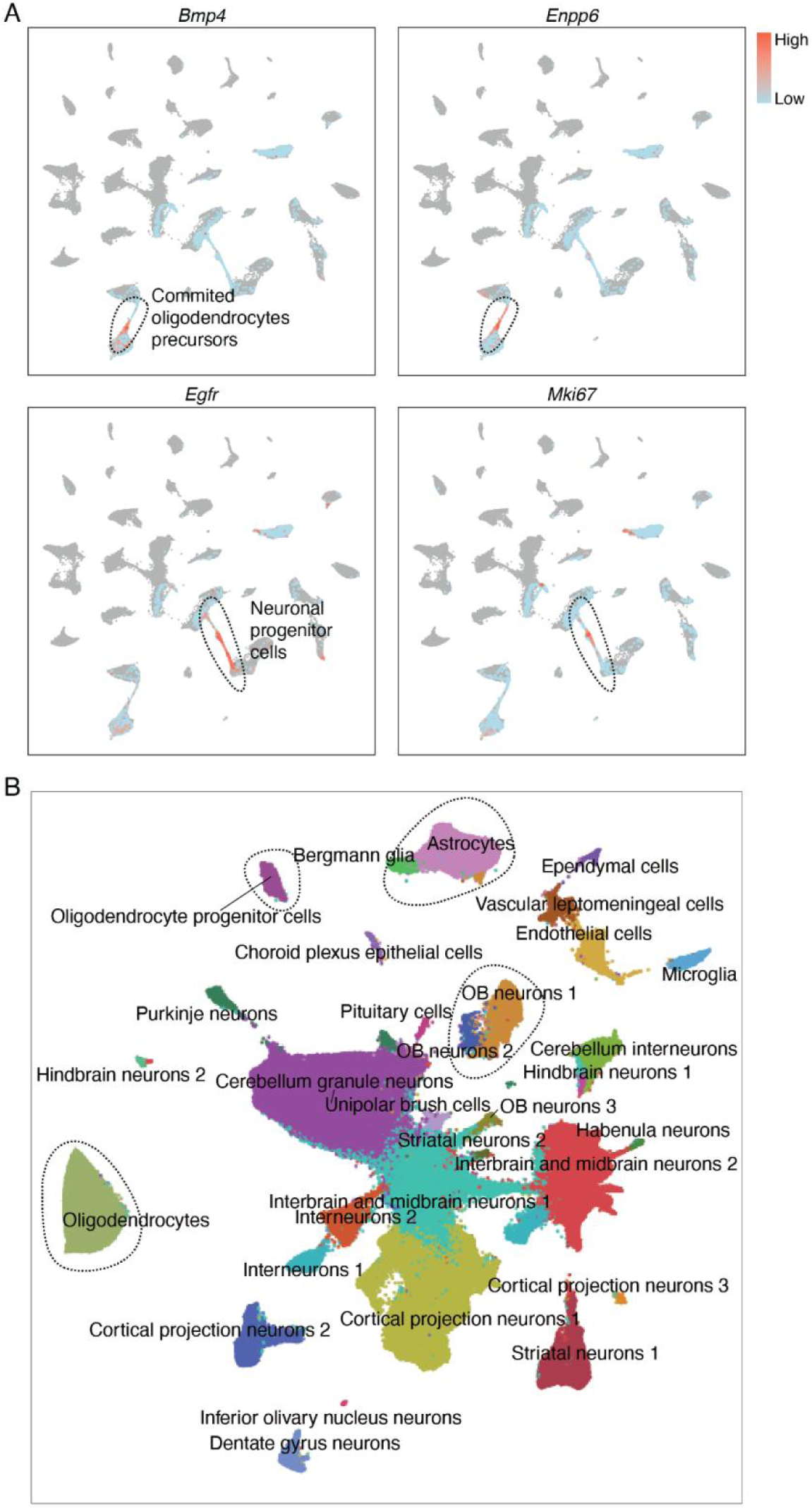
*TrackerSci* facilitates identifying continuous cell transition trajectory missed in global profiling. (A) UMAP visualization integrating *TrackerSci* dataset and *EasySci* brain cell atlas, same as Figure 3C. EdU+ cells profiled by *TrackerSci* are colored by markers for committed oligodendrocyte precursors (top) and neuronal progenitor cells (bottom); and the rest of cells are colored in grey. (B) UMAP visualization of the full brain atlas dataset (∼1.5 million cells) with the same parameter settings as in Figure 3C. Neurogenesis and oligodendrogenesis-related cell types are separated into distinct clusters, while the “bridge” cells in the intermediate stages are missing.

**Figure S8.**
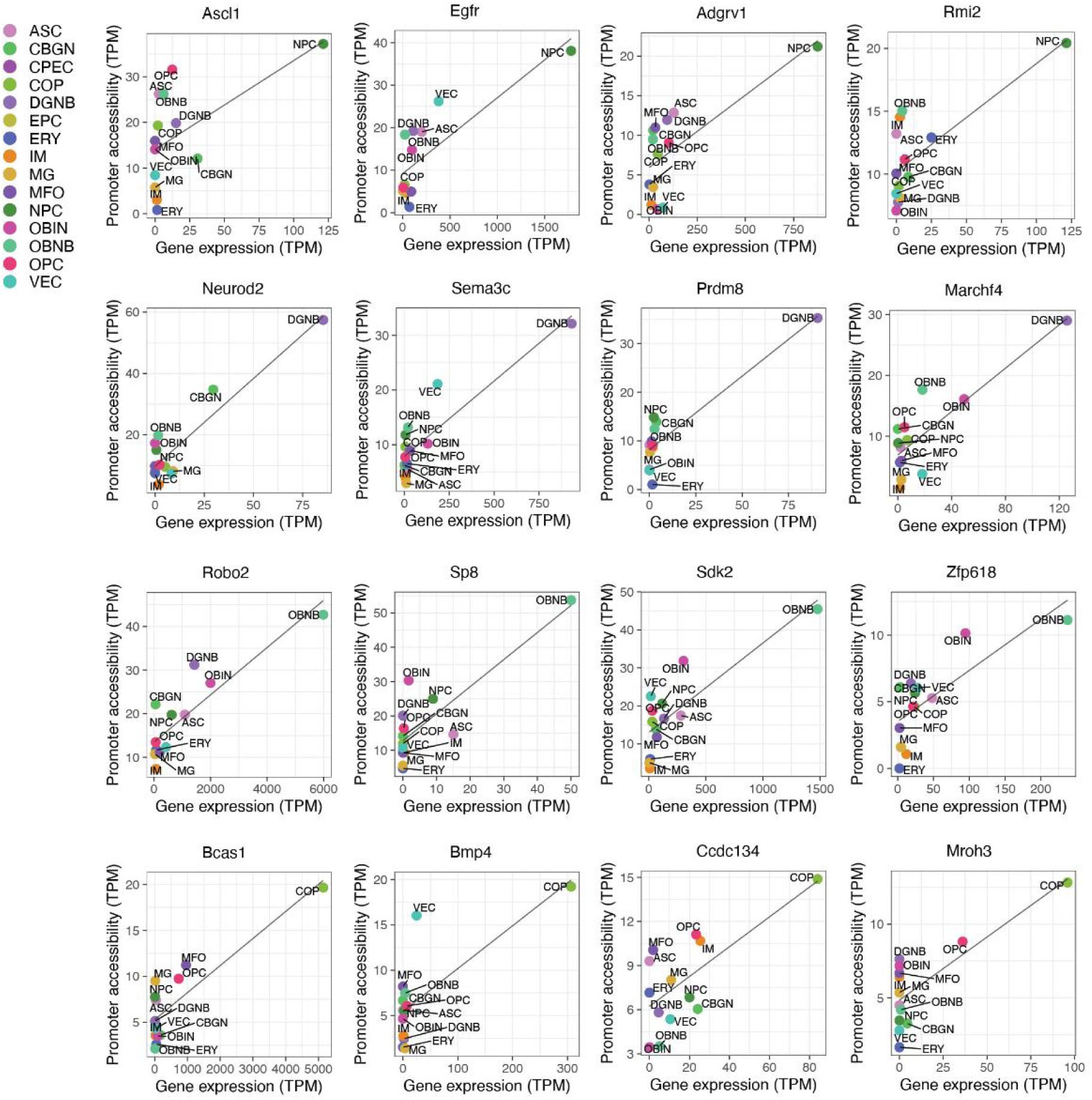
Identifying canonical and novel gene markers of neuronal progenitors and oligodendrocyte precursors. Each scatter plot shows the correlation between expression and promoter accessibility of known (left two columns) or novel (right two columns) cell-type-specific gene markers, together with a linear regression line.

**Figure S9.**
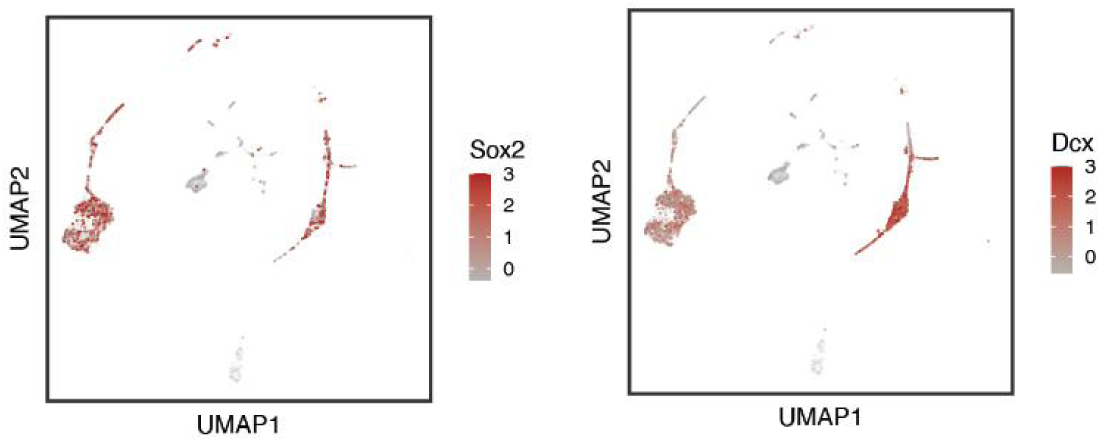
Low cell-type-specificity of certain canonical neurogenesis markers. UMAP plots showing the expression of canonical neurogenesis markers (*Sox2 and Dcx*) across different cell types. The single-cell expression data (UMI count) were normalized first by the total number of reads for each cell and then log-transformed, column centered, and scaled.

**Figure S10.**
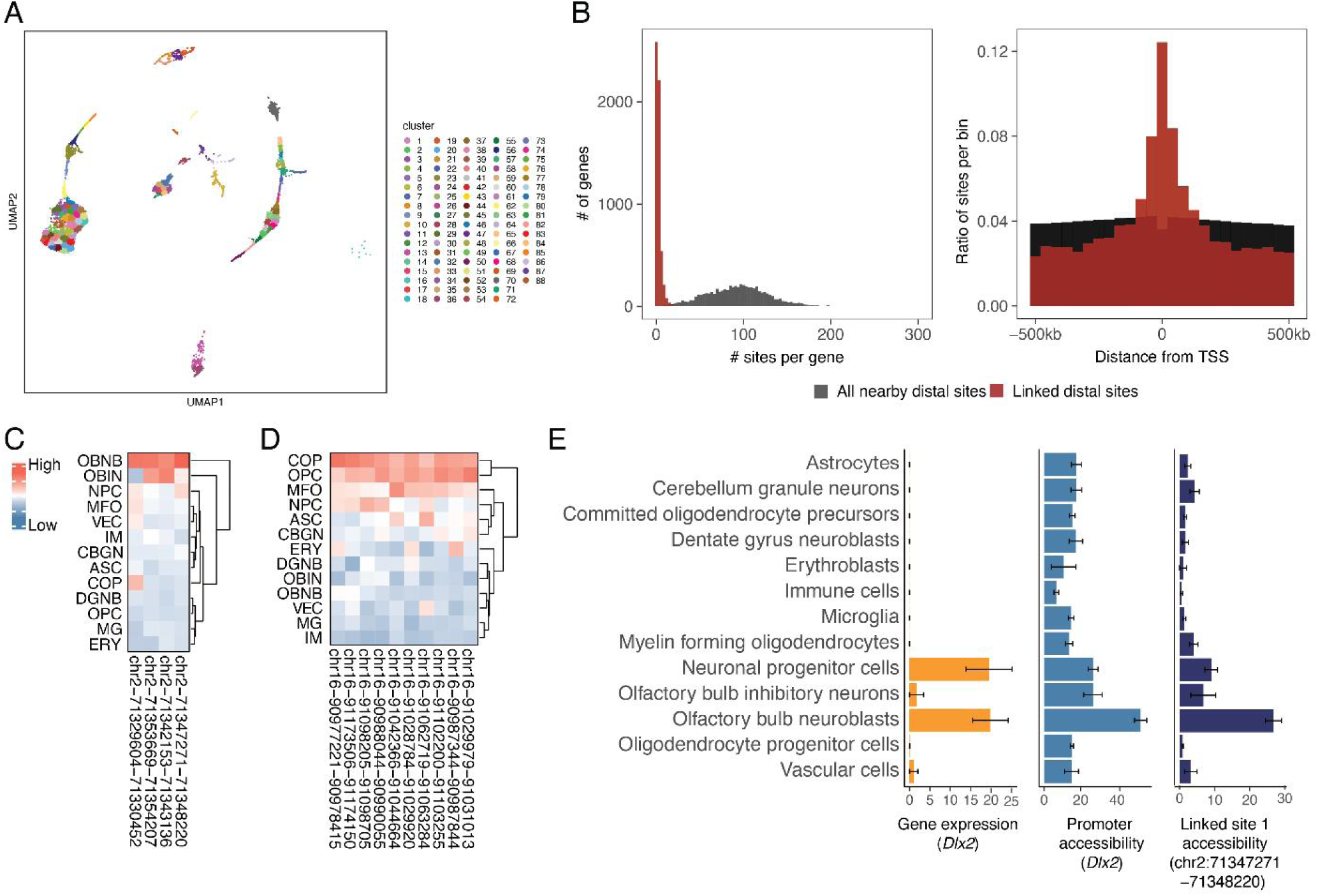
Linking cis-regulatory elements and their regulated genes. (A) UMAP visualization of EdU+ cells as in Figure 1D and 1E, colored by k-means clustering ID. (B) The left histogram shows the number of accessible sites per gene. The right histogram shows the distance distribution of accessible sites within 500 kb of genes. Both plots include all nearby accessible sites (colored in black) and the linked accessible sites (colored in red). (C) Heatmap showing the cell-type-specific peak accessibility of four *Dlx2* linked sites. Cell types are ordered by hierarchical clustering. (D) Heatmap showing the cell-type-specific peak accessibility of ten *Olig2* linked sites. Cell types are ordered by hierarchical clustering. (E) Barplots showing the average expression, the accessibility of promoter and linked distal sites for neurogenesis marker *Dlx2* across different cell types. Gene expression values for each cell type were quantified by transcripts per million (TPM). Site accessibilities for each cell were quantified by the number of reads per million. Error bars represent standard errors of the means.

**Figure S11.**
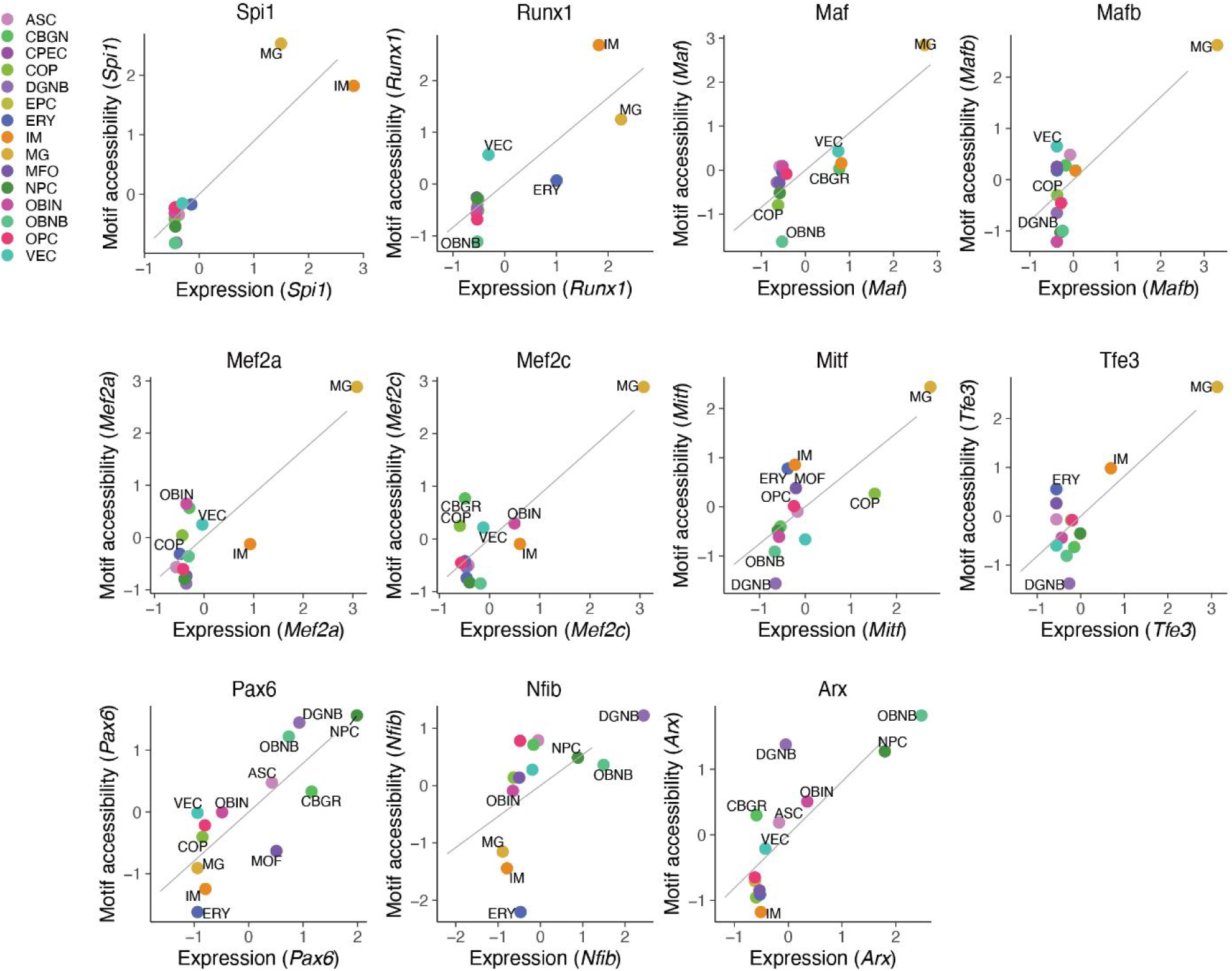
Identifying key transcription factor regulators of the newborn cells. Each scatter plot shows the correlation between cell-type-specific gene expression and motif accessibility for known TF regulators, together with a linear regression line.

**Figure S12.**
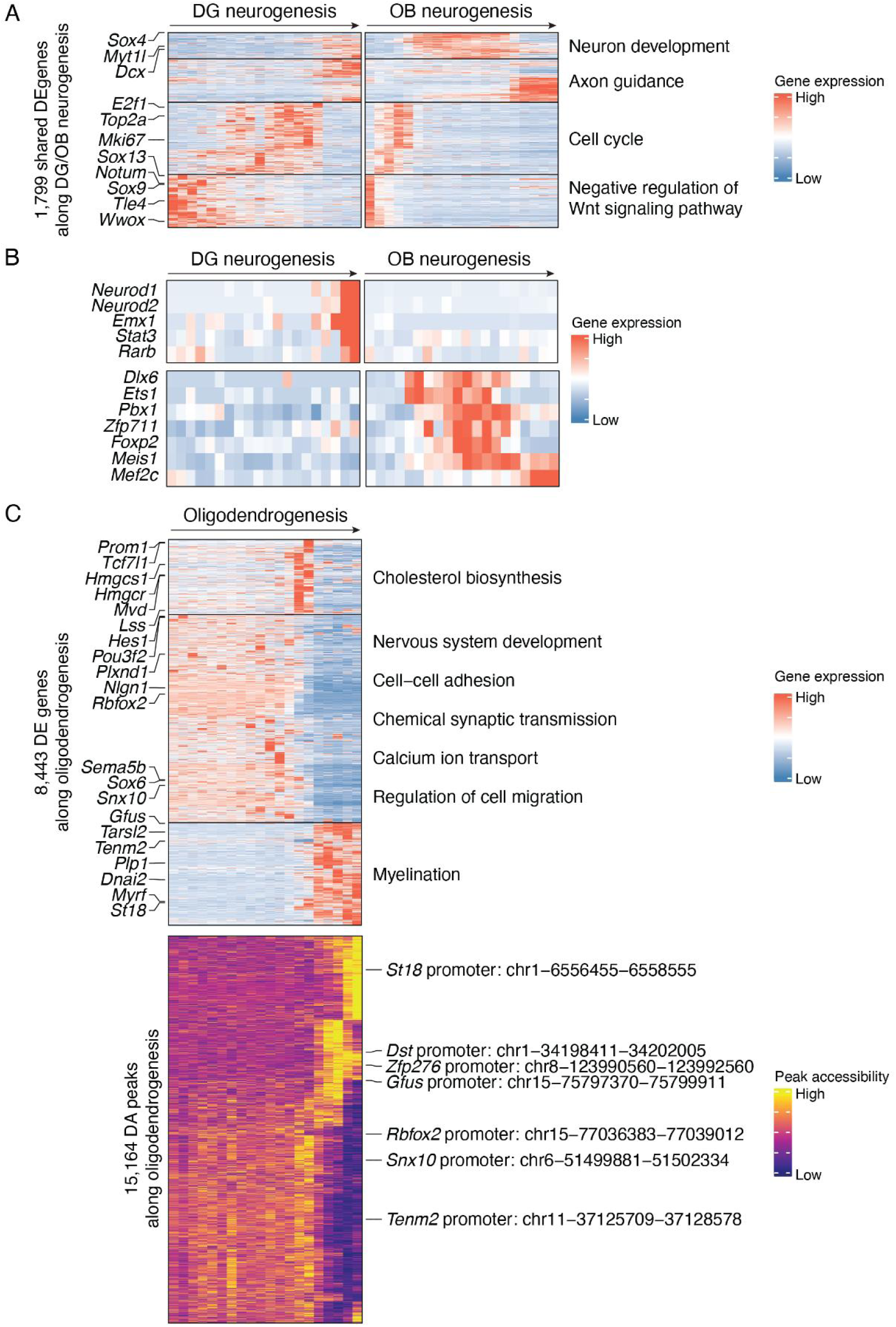
Characterizing gene expression and chromatin accessibility dynamics along adult neurogenesis and oligodendrogenesis. (A) Heatmap showing the dynamics of gene expression of 1,799 shared DE genes along DG neurogenesis (left) and OB neurogenesis (right). Genes are ordered and clustered by hierarchical clustering. Representative gene names (left) and enriched pathways (right) for each gene group are labeled. (B) Heatmap showing examples TFs exhibiting trajectory-specific gene expression dynamics: *Neurod1*, *Neurod2*, *Emx1*, *Stat3* and *Rarb* are uniquely upregulated in DG neurogenesis, while *Dlx6*, *Ets1*, *Pbx1*, *Zfp711*, *Foxp2*, *Meis1* and *Mef2c* are uniquely upregulated in OB neurogenesis. (C) Heatmap showing the dynamics of 8,443 DE genes (top) and 15,164 DA sites (bottom) along the oligodendrogenesis trajectory. Genes are ordered and clustered based on hierarchical clustering. Representative gene names (left) and enriched pathways (right) for each gene group are labeled. Peaks are ordered based on hierarchical clustering, and peaks corresponding to promoters of known and novel oligodendrogenesis markers are labeled.

**Figure S13.**
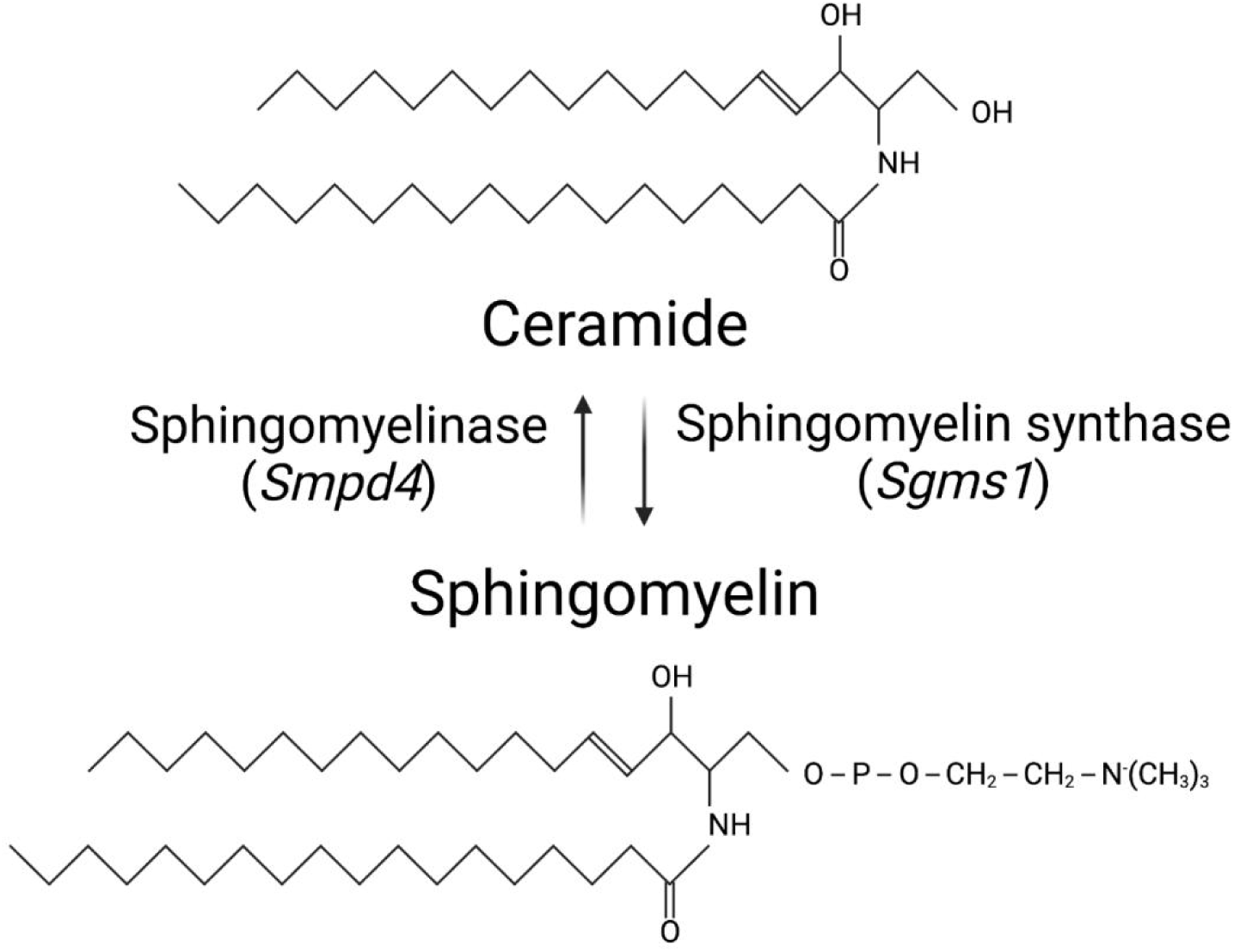
Overview of ceramide/sphingomyelin metabolism. Sphingomyelin production from ceramide is catalyzed by sphingomyelin synthase and is hydrolyzed to ceramide by sphingomyelinase.

**Figure S14.**
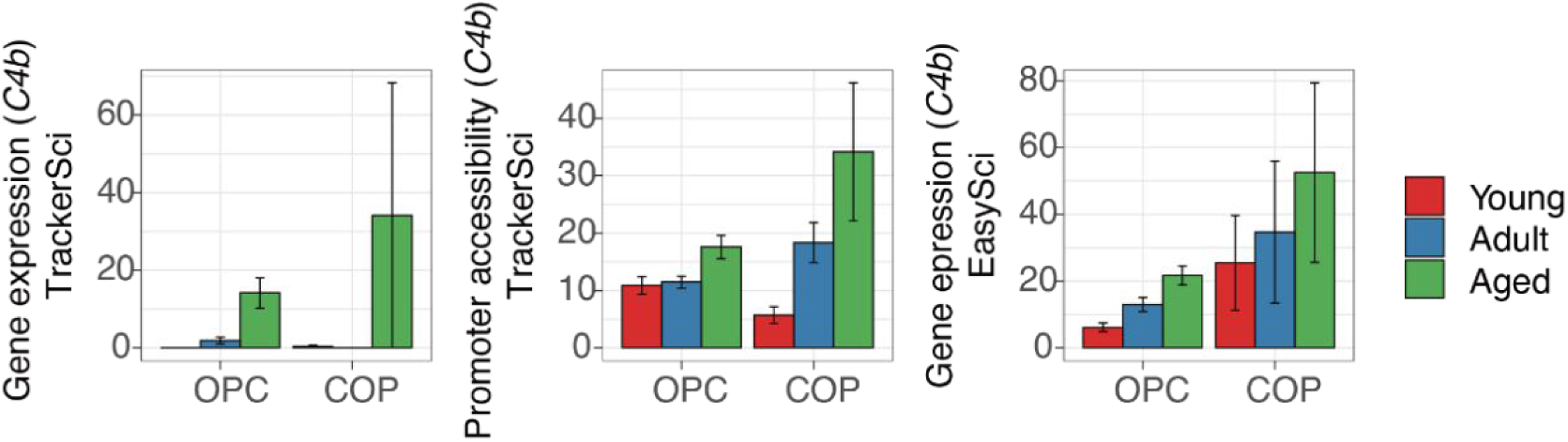
Increased expression of *C4b* in oligodendrocyte progenitor cells. Barplots showing the gene expression (left) and promoter accessibility (middle) of *C4b* from the *TrackerSci* dataset, and the gene expression of *C4b* from the *EasySci* dataset (right) in oligodendrocyte progenitor cells(OPC) and committed oligodendrocyte precursors(COP), quantified by transcripts per million(TPM) for gene expression and reads per million for promoter accessibility. Error bars represent standard errors of the means.

**Figure S15.**
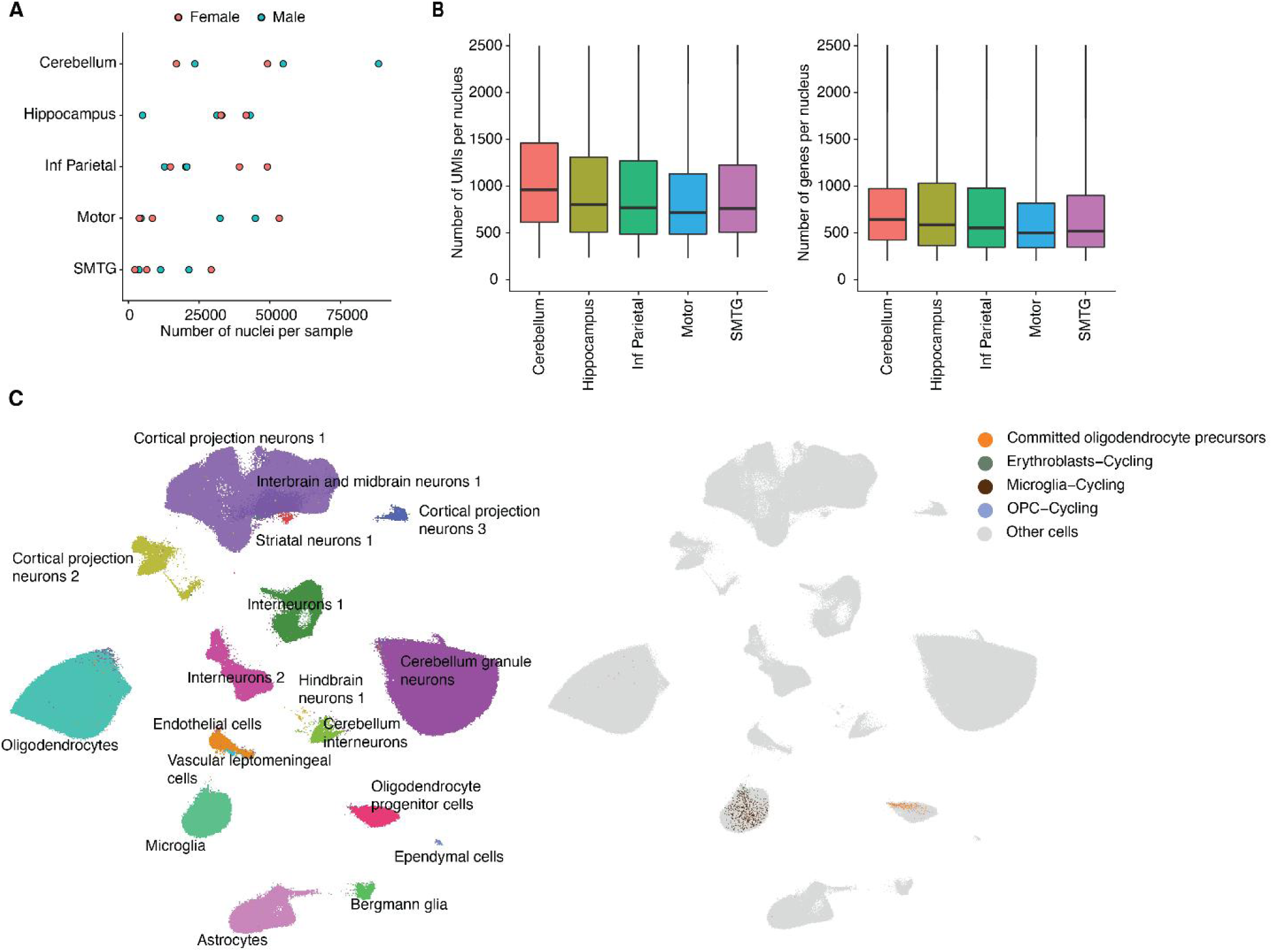
Performance, quality control and characterization of proliferating and differentiating cells in the human brain dataset. (A) Scatter plot showing the number of single-cell transcriptomes profiled in each human sample across five regions, colored by sexes. (B) Boxplots showing the number of unique transcripts (left) and genes (right) detected per nucleus profiled by *EasySci* in the human dataset. For all box plots: middle lines, medians; upper and lower box edges, first and third quartiles, respectively; whiskers, 1.5 times the interquartile range; and circles are outliers. (C) UMAP visualization of the full human brain dataset (∼800,000 cells) with the same parameter settings as in Figure 7A, colored by main cell types (left) and cycling and differentiating cells (right). Note that rare cycling and differentiating cells are masked in the main clustering analysis.

**Figure S16.**
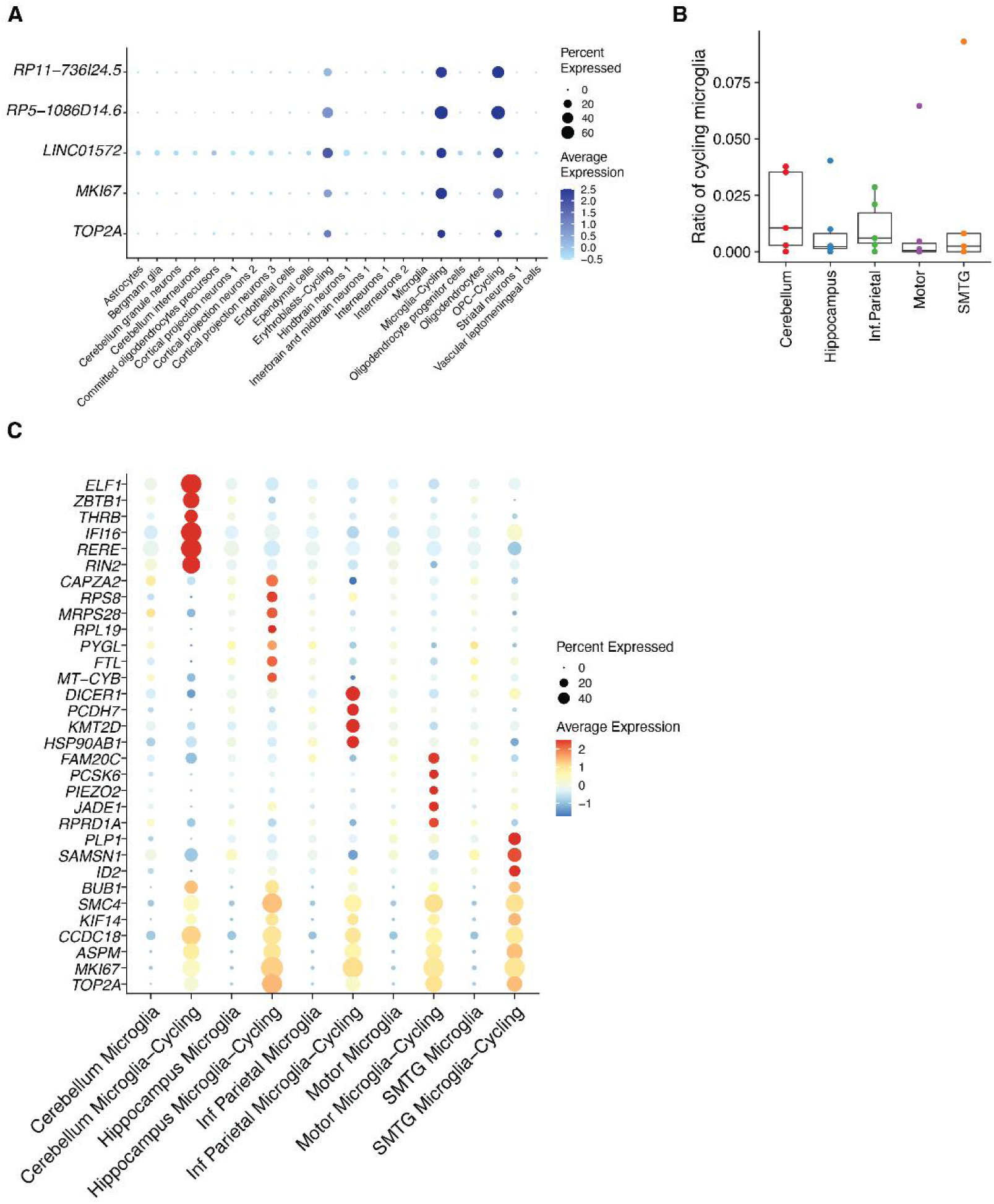
Identifications of cycling cells and region-specific gene expression signatures of cycling microglia in the human brain. (A) Dotplot showing the markers for cycling cells, including novel noncoding RNA (*RP11-736I24.5*, *RP5-1086D14.6* and *LINC01572*) and canonical cycling markers (*MKI67* and *TOP2A*). (B) Boxplot showing the fraction of cycling microglia to the rest of microglia cells across different brain regions in each sample. For all box plots: middle lines, medians; upper and lower box edges, first and third quartiles, respectively; whiskers, 1.5 times the interquartile range; and all data points are shown. (C) Dotplot showing examples of region-specific and shared gene expression signatures for cycling microglia across brain regions.

**Figure S17.**
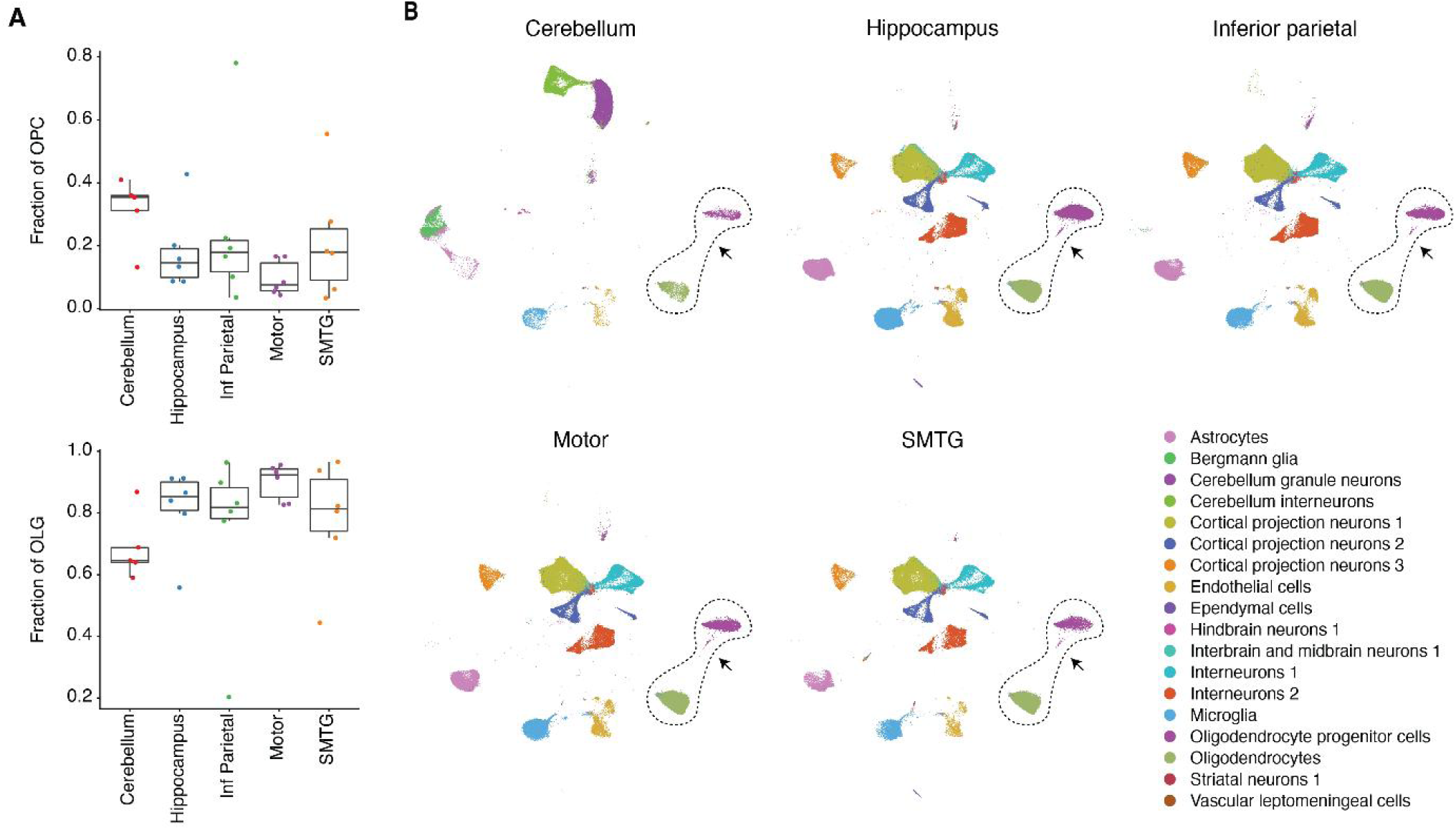
Reduction of oligodendrogenesis in the human cerebellum. (A) Boxplot showing the fraction of oligodendrocyte progenitor cells (OPC, left) and mature oligodendrocytes (OLG) among oligodendrogenesis-related cells across different brain regions in each sample. For all box plots: middle lines, medians; upper and lower box edges, first and third quartiles, respectively; whiskers, 1.5 times the interquartile range; and all data points are shown. (B) UMAP plot same as in Figure 7A splitted by five brain regions colored by main cell types, indicating the loss of intermediate oligodendrogenesis cells in the cerebrum.

## Materials and Methods

### Animals

The C57BL/6 mice were obtained from The Jackson Laboratory. All animal procedures were in accordance with institutional, state, and government regulations and approved under the IACUC protocol 21049.

### EdU Labeling of Mammalian Cell Culture

HEK293T and NIH/3T3 cells (gift from B. Martin, University of Washington) were cultured in 10 cm dishes at 37°C with 5% CO_2_ in high glucose DMEM (Gibco, 11965-118) supplemented with 10% Fetal Bovine Serum (Sigma-Aldrich, F4135) and 1X penicillin-streptomycin (Gibco, 15140-122).

EdU (5-ethynyl-2’-deoxyuridine) (Thermo Fisher Scientific, A10044) was added to culture media at 10 µM final concentration for 1 hour. After labeling, cells were harvested with 0.25% trypsin-EDTA. HEK293T and NIH/3T3 cells were combined at a 1:1 ratio, washed with ice-cold PBS, and lysed in 1 mL ice-cold EZ lysis buffer (Millipore Sigma, NUC101). The nuclei were then fixed on ice with 1% formaldehyde (Thermo Fisher Scientific, 28906) for 10 minutes and washed with EZ lysis buffer, filtered with 40 µm cell strainers (Ward’s Science, 470236-276), and resuspended in Nuclei Suspension Buffer (NSB) (10 mM Tris-HCl pH 7.5 (VWR, 97062-936), 10 mM NaCl (VWR, 97062-858), 3 mM MgCl_2_ (VWR, 97062-848) supplemented with 0.1% SUPERase•In™ RNase Inhibitor (Thermo Fisher Scientific, AM2696) and 1% BSA for *TrackerSci-RNA* or supplemented with 0.1% Tween-20 (Sigma, P9416-100ML), 1x cOmplete™, EDTA-free Protease Inhibitor Cocktail (Sigma, 11873580001) and 0.1% IGEPAL® CA-630 (VWR, IC0219859650) for *TrackerSci-ATAC* experiments).

### EdU Labeling of Mouse Tissues

C57BL/6J mice of different age groups and 5xFAD transgenic mice (MMRRC Strain #034840-JAX) were obtained from The Jackson Laboratory. Mice were injected intraperitoneally with 50 mg/kg of EdU in PBS at 24-hour intervals for five days, and mouse brains were harvested 24 hours after the final injection.

C57BL/6J mice obtained from The Jackson Laboratory were labeled and harvested for pulse-chase labeling at various time points. Specifically, four mice (two male and two female) were injected intraperitoneally with 50 mg/kg of EdU in PBS for 3 days at 24-hour intervals, and brains were harvested 24 hours after the final injection. 12 mice were injected intraperitoneally with 50 mg/kg of EdU in PBS for five days at 24-hour intervals. In addition, for five-day injections, four mice (two male and two female) were harvested 1 day, 3 days, and 5 days after the final injection.

### Tissue collection and nuclei isolation

Whole brains were extracted from mice, immediately snap-frozen in liquid nitrogen, and stored at -80°C upon further usage. For nuclei isolations, thawed brains were cut into small pieces with fine scissors (Fine Science Tools, 14060-09) in 1 mL ice-cold PBS with 1% SUPERase•In™ RNase Inhibitor and 1% BSA, pelleted, resuspended in 1.5 mL Nuclei Isolation Buffer (EZ Lysis Buffer supplemented with 1% SUPERase•In™ RNase Inhibitor, 1% BSA and 1X cOmplete™ EDTA-free Protease Inhibitor Cocktail) for 5 minutes on ice, and homogenized through 40 µm cell strainers (VWR, 470236-276) with the rubber tips of syringes. Then, extracted nuclei were pelleted, fixed in 1% formaldehyde on ice for 10 minutes, washed twice with NSB, and divided into two aliquots for both sci-RNA-seq and sci-ATAC-seq profiling. Nuclei subjected to sci-RNA-seq were briefly sonicated (Diagenode, low power mode for 12 seconds) to reduce clumping. Finally, nuclei were filtered through pluriStrainer Mini 20 µm filters (Pluriselect, 43-10020-70), resuspended in 100 µL NSB, snap frozen in liquid nitrogen, and stored at -80°C until further usage.

### Human brain sample

Twenty-nine post-mortem human brain samples across five regions and six individuals (who were cognitively normal proximal to death) ranging from 70-94 years of age at death, were collected from the University of Kentucky AD Center Tissue Bank (Nelson et al., 2018; Schmitt et al., 2012). Each surveyed sample underwent rigorous quality control including short PMI (<4 hrs). Established strategies were used to extract high-quality nuclei from frozen postmortem brain samples. Extracted nuclei were then fixed with formaldehyde, diluted, and flash-frozen for storage. For *EasySci* transcriptome profiling, nuclei from all samples were thawed and deposited into different wells for barcoded reverse transcription (RT), such that the first index identifies the source of each cell. The library was sequenced across two Illumina NovaSeq™ 6000 sequencer runs, altogether yielding 12 billion reads for ∼900,000 cells (∼13,000 sequencing reads per cell).

### TrackerSci-RNA

Detailed step-by-step *TrackerSci-RNA* protocol is included as a supplementary file (**Supplementary file 1**). Briefly, EdU staining was performed on thawed nuclei using Click-iT Plus EdU Alexa Fluor™ 647 Flow Cytometry assay Kit (Thermo Fisher Scientific, 10634). A 500 µL reaction buffer (prepared following the manufacturer’s protocol) supplemented with 1% SUPERase•In™ RNase Inhibitor was added directly to the nuclei suspension, mixed well and left in RT for 30 minutes. Then, nuclei were spun down for 5 minutes at 500g (4°C), washed once with 500 µL of 1X Click-iT saponin-based permeabilization and wash reagent, resuspended in 1 mL NSB with 1:20 dilution of 0.25 mg/ml 4’,6-diamidino-2-phenylindole (DAPI, Invitrogen D1306) and FACS sorted. Alexa647 and DAPI positive nuclei were sorted into 96-well plates with each well (250∼500 nuclei/well) containing 4 µL of NSB. Sorted plates were briefly centrifuged, mixed with 1 µL of 50 µM oligo-dT primer (5ʹ-ACGACGCTCTTCCGATCTNNNNNNNN[10bp-index]TTTTTTTTTTTTTTTTTTTTTTTTTTTTTTVN-3ʹ, where “N” is any base and “V” is either “A”, “C” or “G”, IDT) and 0.5 µL 10 mM dNTP mix (Thermo Fisher Scientific, R0194) and denatured at 55°C for 5 minutes and immediately placed on ice. 3.5 µL of first-strand reaction mix, containing 2 µL 5X SuperScript™ IV Reverse Transcriptase Buffer (Invitrogen, 18090200), 0.5 µL 100 mM DTT (Invitrogen, P2325), 0.5 µL SuperScript™ IV Reverse Transcriptase (Invitrogen, 18090200), 0.5 μL RNaseOUT™ Recombinant Ribonuclease Inhibitor (Invitrogen, 10777019) was then added to each well. Reverse transcription was carried out by incubating plates at the following temperature gradient: 4°C 2 minutes, 10°C 2 minutes, 20°C 2 minutes, 30°C 2 minutes, 40°C 2 minutes, 50°C 2 minutes and 55°C 10 minutes, and was stopped by adding 1 μL of 18 mM EDTA (VWR, 97062-656) to each well. All nuclei were then pooled, stained with DAPI at a final concentration of 3 μM, and sorted at 25 nuclei per well into 5 μL EB buffer. Cells were gated based on DAPI and Alexa647 such that singlets were discriminated from doublets and EdU+ cells were purified. 0.66 μL mRNA Second Strand Synthesis buffer and 0.34 µL mRNA Second Strand Synthesis enzyme (NEB, E6111L) were then added to each well. Second strand synthesis was carried out at 16°C for 1 hour. 6 μL tagmentation reaction mix (made by mixing 0.5 μL self-loaded Tn5 with 200 μL Tagmentation buffer containing 20 mM Tris-HCl pH 7.5, 20 mM MgCl_2_, 20% Dimethylformamide (Fisher, AC327175000)) was added to each well and tagmentation was performed at 55°C for 5 minutes. After tagmentation, each well was mixed with 0.4 μL 1% SDS, 0.4 μL BSA (NEB, B90000S), and 2 μL of 10 μM P5 primer (5’-AATGATACGGCGACCACCGAGATCTACA[i5]CCCTACACGACGCTCTTCCGATCT-3’, IDT), and incubated at 55°C for 15 minutes. Then, 2 μL 10% Tween-20, 1.2 μL nuclease-free water and 2 μL of 10 μM indexed P7 primer (5’-CAAGCAGAAGACGGCATACGAGAT[i7]GTCTCGTGGGCTCGG-3’, IDT), and 20 μL NEBNext High-Fidelity 2X PCR Master Mix (NEB, M0541L) were added to each well. Amplification was carried out using the following program: 72°C for 5 minutes, 98°C for 30 seconds, 18-22 cycles of (98°C for 10 seconds, 66°C for 30 seconds, 72°C for 1 minute), and a final 72°C for 5 minutes. After PCR, samples were pooled and purified using 0.8 volumes of AMPure XP beads (Beckman Coulter, A63882) twice. Library concentrations were determined by Qubit (Invitrogen, Q33231), and the libraries were visualized by electrophoresis on a 2% E-Gel™ EX Agarose Gels (Invitrogen, G402022). All RNA-seq libraries were sequenced on the NextSeq 1000 platform (Illumina) using a 100 cycle kit (Read 1: 58 cycles, Read 2: 60 cycles, Index 1: 10 cycles, Index 2: 10 cycles). The *TrackerSci-RNA* libraries were sequenced to ∼70,000 reads per cell.

### TrackerSci-ATAC

Detailed step-by-step *TrackerSci-ATAC* protocol is included as a supplementary file (**Supplementary file 1**). EdU staining was performed on thawed nuclei using Click-iT Plus EdU Alexa Fluor™ 647 Flow Cytometry assay Kit (Thermo Fisher Scientific, 10634). A 500 μL reaction buffer (prepared following the manufacturer’s protocol) supplemented with 1X cOmplete™ EDTA-free Protease Inhibitor Cocktail was added directly to the nuclei suspension, mixed well, and left in RT for 30 minutes. Then, nuclei were spun down for 5 minutes at 500g (4°C), washed once with 500 µL of 1X Click-iT saponin-based permeabilization and wash reagent, resuspended in 1 mL NSB with 1:20 dilution of 0.25 mg/ml 4’,6-diamidino-2-phenylindole (DAPI) and FACS sorted. Alexa647 and DAPI positive nuclei were sorted into 96-well plates with each well (250∼500 nuclei/well) containing 4 μL of NSB. Sorted plates were briefly centrifuged, mixed with 5 μL 2x TD buffer (20 mM Tris-HCl pH 7.5, 20 mM MgCl_2_, 20% Dimethylformamide) and 1 μL barcoded Tn5. Tagmentation reaction was performed at 55°C for 30 minutes and stopped by adding 11 μL 2X Stop buffer (40 mM EDTA, 1 mM Spermidine (Sigma, S0266)) to each well. All nuclei were then pooled, stained with DAPI at a final concentration of 3 μM, and sorted at 25 nuclei per well into 5 μL EB buffer. Cells were gated based on DAPI and Alexa647 such that singlets were discriminated from doublets and EdU+ cells were purified. After sorting, each well was mixed with 0.25 μL 18.9 mg / mL proteinase K (Sigma, 3115828001), 0.25 µL 1% SDS and 0.5 µL nuclease-free water, and reverse crosslinking was performed at 65°C for 16 hours. Then, 2 μL 10% Tween-20 was added to each well to quench the SDS. Following on, 1 μL of 10 μM indexed P5 primer (5′-AATGATACGGCGACCACCGAGATCTACA[i5]CCCTACACGACGCTCTTCCGATCT-3′, IDT), 1 μL of 10 μM indexed P7 primer (5’-CAAGCAGAAGACGGCATACGAGAT[i7]GTGACTGGAGTTCAGACGTGTGCTCTTCCGATCT-3’, IDT) and 10 μL NEBNext High-Fidelity 2X PCR Master Mix were added into each well. Amplification was carried out using the following program: 72°C for 5 minutes, 98°C for 30 seconds, 15-16 cycles of (98°C for 10 seconds, 66°C for 30 seconds, 72°C for 1 minute), and a final 72°C for 5 minutes. Final PCR products were pooled and purified by a Zymoclean DNA clean and concentration kit (Zymoresearch, D4014). Library concentrations were determined by Qubit, and the libraries were visualized by electrophoresis on a 2% E-Gel™ EX Agarose Gels. All ATAC-seq libraries were sequenced on the NextSeq 1000 platform (Illumina) using a 100 cycle kit (Read 1: 58 cycles, Read 2: 60 cycles, Index 1: 10 cycles, Index 2: 10 cycles). The *TrackerSci-ATAC* libraries were sequenced to ∼120,000 reads per cell.

### *TrackerSci-RNA* data processing

Read alignment and gene count matrix generation for the scRNA-seq were performed using the pipeline we developed before (Cao et al., 2017). Briefly, base calls were converted to fastq format and demultiplexed using Illumina’s bcl2fastq/v2.19.0.316 tolerating one mismatched base in barcodes (edit distance (ED) < 2). The RT barcode for each read was corrected to its nearest barcode (edit distance (ED) < 2), and reads with uncorrected barcodes (ED >= 2) were removed. Demultiplexed reads were then adaptor clipped using trim_galore/v0.4.1 (https://github.com/FelixKrueger/TrimGalore) with default settings. Trimmed reads were mapped to a chimeric reference genome of human and mouse (hg19/mm10) for the species-mixing experiment and to the mouse only (mm39) for mouse brain experiments, using STAR/v2.5.2b (Dobin et al., 2013) with default settings. Uniquely mapping reads were extracted, and duplicates were removed using the unique molecular identifier (UMI) sequence, reverse transcription (RT) index, and read 2 end-coordinate (i.e. reads with identical UMI, RT index, and tagmentation site were considered duplicates). Finally, mapped reads were split into constituent cellular indices by further demultiplexing reads using the RT index.

To generate digital expression matrices, we calculated the number of strand-specific UMIs for each cell mapping to the exonic and intronic regions of each gene with python/v2.7.18 HTseq package (Anders et al., 2015). For multi-mapped reads, reads were assigned to the closest gene, except in cases where another intersected gene fell within 100 bp to the end of the closest gene, in which case the read was discarded. For most analyses, we included both expected-strand intronic and exonic UMIs in per-gene single-cell expression matrices. Exonic and intronic gene count matrices were used in RNA velocity analysis.

For the species-mixing experiment, RNA barcodes with more than 200 UMIs and 100 unique genes were identified as real cells, and those with fewer than that were discarded. The percentage of uniquely mapping reads for genomes of each species was calculated. Cells with over 90% of UMIs assigned to one species were regarded as species-specific cells, with the remaining cells classified as mixed cells or “collisions”. The collision rate was calculated as the ratio of mixed cells.

### TrackerSci-ATAC data processing

Single-cell ATAC-seq data was performed using a published pipeline (Cao et al., 2018; Cusanovich et al., 2015) with mild modifications. Base calls were converted to fastq format and demultiplexed using Illumina’s bcl2fastq/v2.19.0.316 tolerating one mismatched base in barcodes (edit distance (ED) < 2). The indexed Tn5 barcode for each read was corrected to its nearest barcode (edit distance (ED) < 2), and reads with uncorrected barcodes (ED >= 2) were removed. Demultiplexed reads were then adaptor-clipped using trim_galore/0.4.1 with default settings. Trimmed reads were mapped to a chimeric reference genome of human and mouse (hg19/mm10) for the species-mixing experiment and to the mouse only (mm39) for mouse brain experiments, using STAR/v2.5.2b (Dobin et al., 2013) with default settings. Duplicates were removed by picard MarkDuplicates/v2.25.2 (Broad Institute, 2019) per PCR sample. Deduplicated reads were split into constituent cellular indices by further demultiplexing reads using the Tn5 index.

A snap-format (Single-Nucleus Accessibility Profiles) file was generated from deduplicated bam files using SnapTools/v1.4.8 with default settings (https://github.com/r3fang/SnapTools) (Fang et al., 2021). A cell-by-bin count matrix with 5kb bin size was created from the resulting snapfile. The promoter ratio for each cell was calculated as the number of fragments mapping to genomic bins overlapping with promoter regions(defined as 2kb upstream of the gene body).

For the species-mixing experiment, ATAC barcodes with more than 1000 fragments and more than 0.2 promoter ratio were identified as real cells, and those with fewer than that were discarded. The percentage of uniquely mapping reads for genomes of each species was calculated. Cells with over 90% of reads assigned to one species were considered species-specific cells, with the remaining cells classified as mixed cells or “collisions”. The collision rate was calculated as the ratio of mixed cells.

### Cell filtering, clustering, and annotation for *TrackerSci-RNA*

A digital gene expression matrix was constructed from the raw sequencing data as described above. EdU+ cells and global cells were combined and analyzed together. Cells with less than 200 UMIs and 100 unique genes were discarded. Potential doublet cells and doublet-derived subclusters were detected using an iterative clustering strategy similar to before (Cao et al., 2020). Cells labeled as doublets(by scanpy/v1.6.0 and scrublet/v0.2.3) (Wolf et al., 2018; Wolock et al., 2019) or from doublet-derived sub-clusters were filtered. The downstream dimension reduction and clustering analysis were done by Seurat/v4.0.2 (Hao et al., 2021). Briefly, the dimensionality of the data was reduced by PCA (30 components) first and then with UMAP, followed by Louvain clustering. Clusters were assigned to known cell types based on cell type-specific markers (**Table S2**).

Differentially expressed genes across different cell types were identified using monocle/v2.22.0 (Qiu et al., 2017) with the differentialGeneTest() function. Genes detected in less than 10 cells were filtered out before the analysis. To identify cell type-specific gene markers, we selected genes that were differentially expressed across different cell types (5% FDR, likelihood ratio test), with FC > 2 between the target cell type and the second highest expressed cell type, and with maximum transcripts per million (TPM) > 10 in the target cell types.

### Cell filtering, clustering, and annotation for *TrackerSci-ATAC*

Single-cell ATAC-seq profiles were generated as described above. EdU+ cells and global cells are combined and analyzed together. Cells with less than 1000 fragments and less than 0.2 promoter ratio were discarded. Dimensionality reduction for ATAC-seq data was performed using the snapATAC/v1.0.0 (Fang et al., 2021). A cell-by-bin matrix at 5-kb resolution was used. We focused on bins on chromosomes 1–19, X and Y. High-coverage bins (top 5% bins that overlap with invariant features) or low-coverage bins (bottom 5% bins that represent general inaccessible regions) were filtered out before the analysis. Diffusion maps dimensionality reduction was performed on the filtered cell-by-bin matrix after binarization. UMAP analyses were performed on the top 20 eigenvectors, followed by unsupervised clustering via the densityPeak algorithm implemented in R package densityClust/v0.3 (Rodriguez and Laio, 2014)

We performed integration analysis between the *TrackerSci-RNA* dataset and *TrackerSci-ATAC* dataset to annotate the ATAC dataset. The gene activity score for ATAC cells was computed using the snapATAC function createGmatFromMat() by summing up the counts of bins overlapping with the gene body. A Seurat object was generated using the gene activity matrix and previously calculated diffusion map embeddings for single cell ATAC-seq. Then, variable genes were identified from *TrackerSci-RNA* data and used for identifying anchors between these two modalities. Next, we co-embedded the RNA-seq and ATAC-seq profiles in the same low-dimensional space to visualize all the cells together. We then used overlapped RNA clusters to annotate ATAC cells in the integrated UMAP space. ATAC cells without overlapped RNA cells were removed with careful inspection since they usually represent potential doublets or low-quality cells. Finally, single-cell ATAC dimension reduction, clustering, and integration analysis were rerun on the remaining dataset following the same procedure.

### Peak calling and identifications of cell-type-specific peaks

To define peaks of accessibility across all sites, we used MACS2/v2.1.1 (Zhang et al., 2008). Nonduplicate ATAC-seq reads of cells from each main cell type were aggregated, and peaks were called on each group separately with these parameters: --nomodel --extsize 200 --shift -100 -q 0.1. Peak summits were extended by 250bp on either side and then merged with bedtools/v2.30.0 (Quinlan and Hall, 2010; Zhang et al., 2008), together with gene promoter regions (annotated transcription start site (TSS) in GENCODE VM27 minus/plus 1000 base pairs in a strand-specific manner). Each read alignment was extended by 100 bp upstream and downstream from the insertion site of tagmentation. Cells were determined to be accessible at a given peak if a read from a cell overlapped with the peak. The peak count matrix was generated by a custom python script with the HTseq package (Anders et al., 2015; Quinlan and Hall, 2010; Zhang et al., 2008). Differentially accessible peaks across cell types were identified using monocle/v2.22.0 (Qiu et al., 2017) with the differentialGeneTest() function. Peaks detected in less than 10 cells were filtered out before the analysis. To determine cell-type-specific peak markers, we selected peaks that were differentially accessible across different cell types (5% FDR, likelihood ratio test), with FC > 2 between the target cell type and the second highest expressed cell type, and with TPM > 10 in the target cell types.

### Analysis for linking cis-regulatory elements (CRE) to regulated genes

We aim to identify links between chromatin accessible sites and regulated genes based on their covariance. Only EdU+ cells were kept in this analysis. We first constructed pseudo-cells by aggregating the RNA-seq and ATAC-seq profiles of highly similar cells through k-means clustering the integrative UMAP coordinates using the kmeans function from R package stats/v4.1.2. The k was selected so that the average cell number per subcluster is 150. Subclusters overrepresented by one molecular layer(the percentage of cells from either RNA-seq or ATAC-seq profile greater than ninety percent) were merged with a nearby subcluster. After aggregating cells within each sub-cluster, we obtained a total of 88 pseudo-cells, with a median of 54 cells from RNA-seq profile and 93 cells from ATAC-seq profile. Aggregated count matrices for RNA-seq and ATAC-seq were normalized to transcripts per million(TPM) and log1p transformed. We only retained genes and peaks with TPM value greater than 10 in the maximum expressed pseudo-cells. Then, for each gene, we calculated the Pearson Correlation Coefficient (PCC) between its gene expression and the chromatin accessibility of its nearby accessible sites(minus/plus 500 kb from the TSS) across pseudo-cells. Sites overlapping with minus/plus 1kb from the TSS were considered promoters, while the rest were considered distal regions. To define a threshold at PCC score, we also generated a set of background pairs by permuting the pseudo cell id of the ATAC-seq matrix and with an empirically defined significance threshold of FDR < 0.05, to select significant positively correlated cCRE-gene pairs. We further filtered the linkage by requiring that either the maximum expressed cell types in the RNA profile and the ATAC profile were the same or the top two or top three highest expressed cell types were in the same cell trajectory (Oligodendrogenesis trajectory: OPC, COP, OLG; Astrocytes trajectory: ASC, NPC; DG neurogenesis trajectory: NPC, DGNB; OB neurogenesis trajectory: NPC, OBNB, OBIN). Finally, we only keep the one top linked gene with the highest PCC for each peak.

### Transcription factor analysis

To identify key TF regulators of each main cell type, we searched for TF that can be validated in two molecular layers by correlating gene expression and motif accessibility. First, using the *TrackerSci-ATAC* dataset, we selected the top 300 sites per main cell type (from the differential peak analysis described above, filtered by q-value < 0.05, maximum expressed TPM > 10 and ranked by FC between the highest and the second expressed cell type) to a combined peak set. We then resized the peaks to a fixed length of 500 bp (± 250 bp around the center) and generated a binarized peak-by-motif matrix using the R package motifmatchr/v1.16.0 (Schep, 2017) with the matchMotifs() function to identify the occurrences of motifs in each peak from a filtered collection of the cisBP motif database curated by chromVARmotifs/v0.2.0 (Weirauch et al., 2014; Schep et al., 2017). A matrix of motif-by-cell counts was obtained by multiplying the peak-by-cell matrix with the peak-by-motif matrix, and was aggregated into pseudo-cells based on the k-means clustering described before. We then computed the PCC between the scaled TF motif accessibility and the scaled TF gene expression across pseudo-cells. To select significantly positive and negative correlations of TF gene expression and motif accessibility pairs, we permuted the pseudo cell id of the motif-by-cell matrix to compute a background PCC distribution and selected the TF pairs with an empirically defined significance threshold of FDR < 0.05. In addition, we only keep TF with TPM > 10 in the maximum expressed cell type.

### Trajectory analysis

Cells corresponding to the neurogenesis trajectory (ASC, NPC, DGNB, OBNB and OBIN) or the oligodendrogenesis trajectory (OPC, COP and OLG) from both RNA-seq data and ATAC-seq data were selected for detailed investigation. We next performed UMAP dimension reduction at the trajectory level with the integration function from Seurat (Hao et al., 2021), using the top 3,000 highly variable genes and top 50 PCs. Each cell was assigned a pseudotime value based on its position along the trajectory using monocle3/v1.0.0 function order_cells() (Trapnell et al., 2014). RNA velocity analyses were performed using scVelo/v0.2.3 (Bergen et al., 2020) using the exonic and intronic gene count matrix generated from sci-RNA-seq pipeline to validate the cell differentiation direction and estimate the position of the progenitor cell state. For the two neurogenesis trajectories (DG neurogenesis and OB neurogenesis), pseudotime assignment was calculated separately and scaled so that the cells shared between two trajectories received the same pseudotime value. Specifically, we first used the pseudotime value calculated from the OB trajectory for common progenitor cells in both DG and OB trajectories. We then fitted a linear regression line using R function lm() to predict the OB-pseudotime based on the DG-pseudotime. Then, for cells unique to the DG neurogenesis, we adjusted their pseudotime using the predict() function using DG-pseudotime as input. Gene expression and peak accessibility dynamics along pseudotime were identified using monocle/v2.22.0 (Qiu et al., 2017) with the differentialGeneTest() function with pseudotime values and their main cluster identity as variables. Genes or peaks that passed a significant test (FDR of 5%) were considered as dynamically regulated genes or sites. Furthermore, differential accessible sites along pseudotime were used to infer TF motif accessibility dynamics. We computed a motif deviation score for each single cell using chromVar/v1.4.1 (Schep et al., 2017) with the dynamic peak set (resized to 500 bp) as input. Then, the motif deviation scores of each single cell were rescaled to (0, 10) using R function rescale() and differential accessible motifs were identified using monocle/v2.22.0 with the differentialGeneTest() function. TF motifs that passed a significant test (FDR of 5%) were considered as dynamically regulated motifs. For gene enrichment analysis we used the enrichR (Chen et al., 2013) and the following pathways collections were considered: Panther_2016, Reactome_2016, KEGG_2019_Mouse, GO_Biological_Process_2018, GO_Molecular_Function_2018. For visualizing the dynamics of gene expression, peak accessibility and motif accessibility, we used R package ComplexHeatmap/v2.10.0 (Gu et al., 2016).

### Cell proportion analysis

To quantify the cell-type-specific changes in the proliferation dynamics across conditions, we calculated the fraction of each cell type within EdU+ population from each condition for RNA-seq data and ATAC-seq data separately, which was further multiplied by the median of EdU+ ratio for each group obtained from FACS sorting. For adult WT mice, we only included those that were harvested 24h after five-day labeling to avoid artifacts introduced by the labeling time.

To quantify the effects of aging on cell differentiation dynamics along neurogenesis and oligodendrogenesis trajectories, we applied miloR/v1.3.1 (Dann et al., 2021), a single-cell differential abundance testing framework using k-nearest neighbor (KNN) graphs. We first constructed the KNN graph on the UMAP space for each trajectory using the buildGraph() function with k = 120 for the neurogenesis trajectory and k = 250 for the oligodendrogenesis trajectory. Cell neighborhoods were then defined using the makeNhoods() function and the number of cells from each experiment sample were counted for each neighborhood using the countCells() function. Testing for differential abundance in neighborhoods was performed using the testNhoods() function and significance levels for Spatial FDR of 0.05 were used. Visualization of differential abundance neighborhoods was done using the plotNhoodGraphDA() function.

### Differential analysis of NPC and OPC across aged groups

Differential gene expression analysis across young, adult, and aged groups of NPC and OPC was performed using monocle/v2.22.0 (Qiu et al., 2017) function differentialGeneTest() with the number of genes detected per cell included as a covariant. For adult WT mice, only cells from the animals harvested at 24h after 5-day labeling were included to avoid artifacts introduced by the labeling time. In addition, only differentially expressed genes (> expressed in more than 10 cells) along the neurogenesis or the oligodendrogenesis trajectory were included in the differential gene test. Differentially expressed genes were selected by a q-value cutoff of 0.1, a TPM cutoff of 50 in the maximum expressed group, and with at least 1.5 FC between the maximum expressed group and the minimum expressed group. Next, differentially expressed genes were grouped to aged-depleted genes and aged-enriched genes by the following criteria: for aging-depleted genes, we first selected the genes with minimum expression in aged mice, and only kept those with either maximum expression in young mice or within less than 2 FC between the young group and the adult group. For aging-enriched genes, we first selected the genes with maximum expression in aged mice, and only kept those with either minimum expression in young mice or with less than 2 FC between the young group and the adult group. We then further filtered the DE genes based on the consistency on their promoters or linked sites. For aging-depleted genes, we required that the mean of promoter accessibility or linked site accessibility was at the minimum level in the aged group compared to young and adults. For aging-enriched genes, we required that the mean of promoter accessibility or the linked site accessibility was at the maximum level in the aged group compared to young and adults. Genes that were lowly detected in both promoter accessibility and linked sites (represented by the mean of TPM < 10 in all conditions) were also discarded.

### Integration analysis between *TrackerSci-RNA* and *EasySci-RNA*

Integration analysis of scRNA-seq dataset profiled using *TrackerSci* and *EasySci* was performed using Seurat/v4.0.2 (Hao et al., 2021). We first integrated 14,095 *TrackerSci-RNA* cells (including 5,715 EdU+ cells and 8,380 all brain cells without EdU enrichment) with 126,285 *EasySci-RNA* cells (up to 5,000 cells randomly sampled from each of 31 cell types) in our companion study (Sziraki et al., 2022). Shared variable genes, selected by SelectIntegrationFeatures() function, were used for identifying anchors using FindIntegrationAnchors(). The two datasets were then integrated together with the IntegrateData() function. To visualize all the cells together, we co-embedded all the cells in the same low-dimensional space. We further applied the same integrative analysis strategy to cells matching the same cellular state from both datasets. Specifically, for the neurogenesis trajectory, we integrated 1,214 EdU+ cells from *TrackerSci-RNA* (NPC, OBNB, and OBIN) with 37,258 OB neurons 1 cells from *EasySci-RNA*. For the oligodendrogenesis trajectory, we integrated 3,044 EdU+ cells from *TrackerSci-RNA* (OPC and COP) to 22,718 oligodendrocyte progenitor cells from *EasySci-RNA*. For the microglia, we integrated 600 EdU+ microglia from *TrackerSci-RNA* to 15,754 microglia from *EasySci-RNA*. Microglia subclusters corresponding to peripheral immune cells were excluded before the analysis.

### Quantifications of the self-renewal potential and the differentiation potential

The self-renewal potential was defined as the ratio of newly generated progenitor cells within 5 days of EdU labeling divided by the ratio of total progenitor cells detected from the global population. To account for potential variations due to slight differences of animal ages between *TrackerSci* and the brain cell atlas, we first fitted a linear model between the ages and the ratio of progenitor cells using the *EasySci* data for the following cell type: neuronal progenitor cells, oligodendrocyte progenitor cells, and microglia. We used that to predict the ratio of progenitor cells for each individual mice profiled by *TrackerSci*. We then divided the ratio of newly generated progenitor cells from each 5-day labeled mice by the predicted cellular fraction of the global progenitor pool for the same cell type. A line plot was generated using the median values of proliferation potential for each aged group normalized to the young mice. RNA and ATAC cells were both included, and samples with less than 50 cells were excluded from the calculation. The differentiation potential was quantified by the ratio of differentiated cells divided by all EdU+ cells in the same trajectory. We calculated such a ratio only for oligodendrogenesis trajectory since it’s a unidirectional route. For this analysis, we divided the ratio of committed oligodendrocytes and myelin-forming oligodendrocytes to the ratio of oligodendrocyte progenitor cells for each sample and median values of each age group were used to generate the line plot. RNA and ATAC cells were included, and samples with less than 50 cells were excluded from the calculation.

### Cell filtering, clustering, and annotation for the human dataset

A digital gene expression matrix was constructed from the raw sequencing data as described in our companion study (Sziraki et al., 2022). Potential doublet cells and doublet-derived subclusters were detected using an iterative clustering strategy similar to before (Cao et al., 2020). Cells labeled as doublets(by scanpy/v1.6.0 and scrublet/v0.2.3) (Wolf et al., 2018; Wolock et al., 2019) or from doublet-derived sub-clusters were filtered. To identify distinct clusters of cells corresponding to different cell types in the human data, we performed the downstream dimension reduction and clustering analysis using Seurat/v4.0.2 (Hao et al., 2021). Briefly, the dimensionality of the data was reduced by PCA (50 components) first and then with UMAP, followed by Louvain clustering. We then co-embedded the human data with the mouse brain atlas from profiled in our companion study (Sziraki et al., 2022) through Seurat (Stuart et al., 2019), and clusters were annotated based on overlapped cell types. The annotations were manually verified and refined based on marker genes.

### Integration analysis between human and mouse

Integration analysis of scRNA-seq dataset of human and mouse was performed using Seurat/v4.0.2 (Hao et al., 2021). Similar to the integration of mouse dataset profiled between *TrackerSci-RNA* and *EasySci-RNA*, we first integrated 14,095 mouse cells (including 5,715 EdU+ cells and 8,380 all brain cells without EdU enrichment) with 71,743 human cells (up to 5,000 cells randomly sampled from each of 18 cell types) to construct a coembedding UMAP space. We then project the rest of human cells into this UMAP structure using MapQuery() and TransferData() function. Cycling cells and committed oligodendrocytes from the human dataset were extracted based on the UMAP coordinates overlapping with mouse cells. Cycling cells were subjected to sub-clustering analysis for identifying their cell types. Markers for cycling cells were identified by comparing them to the rest of all cells using the Seurat function FindMarkers().

### Identifications of shared and unique features between human and mouse oligodendrogenesis

To construct a continuous oligodendrogenesis trajectory shared between human and mouse, we subjected all 4,194 oligodendrogenesis-related cells (OPC, COP and OLG) from mouse data and took 2,188 oligodendrogenesis-related cells from human data (including all of 188 cells from COP and randomly sampled 1,000 cells from OPC and OLG) to integration analysis using Seurat/v4.0.2. Each cell was assigned a pseudotime value based on its position along the trajectory using monocle3 function order_cells(). For human cells, gene expression dynamics along pseudotime were identified using monocle/v2.22.0 (Qiu et al., 2017) with the differentialGeneTest() function with pseudotime values and their main cluster identity (i.e, OPC, COP and OLG) as variables. For mouse cells, we used the results from DE gene analysis along pseudotime calculated before. Conserved gene expression dynamics were selected by a q-value cutoff of 0.05, a TPM (transcript per million) cutoff of 50 in the same maximum expressed stage in both species. This reveals 1,162 DE genes along oligodendrogenesis shared between human and mouse. To select genes with species-unique expression dynamics, we filtered the DE genes with the following criteria: significantly changed along pseudotime (q-value <0.05) and TPM of the maximum expressed stage larger than 50 in one species, while no significantly changed (q-value >0.05) and TPM of the maximum expressed stage less than 50 in the other species. This reveals 458 and 361 DE genes along oligodendrogenesis unique to human and mouse respectively. For visualizations of gene expression dynamics, we use R package ComplexHeatmap/v2.10.0 and the genes were ordered by the hierarchical clustering implemented in the function Heatmap().

### Analysis of region-specific oligodendrogenesis

To study region-specific effects of oligodendrogenesis, we quantified the ratio of each stage (OPC, COP and OLG) within all the cells along the oligodendrogenesis trajectory for each region. Cycling Oligodendrocyte progenitor cells were not included into the calculation. Statistical analysis was performed by comparing the ratio of COP to OPC in cerebellum vs. non-cerebellum cells using Fisher exact test. To study the region-specific transcriptional controls of each stage along oligodendrogenesis, we performed differential expression analysis across regions using monocle/v2.22.0 with the differentialGeneTest() function. Region-specific gene expression signatures were selected by the following cutoffs: q-value < 0.05, with FC > 2 between the maximum expressed region and the second highest expressed region, and with maximum transcripts per million (TPM) > 50 in the highest expressed region.

### Code Availability

The detailed experimental protocols and computation scripts of *TrackerSci* were included as supplementary files.

## Supplementary Tables (provided as Microsoft Excel files)

**Supplementary Table 1:** Metadata for animal individuals included in the *TrackerSci* profiling, including 38 animals injected with EdU and 2 animals injected with PBS. For each mouse, the metadata includes the mouse genotype (WT, 5xFAD), age group (young, adult, aged), gender, the exact day of age, DOB (date of birth), DOD (date of death), the time of EdU labeling, and number of cells recovered from *TrackerSci-RNA* and *TrackerSci-ATAC*.

**Supplementary Table 2:** Annotated cell types together with reference gene markers for annotation, number of cells per cell type identified in *TrackerSci-RNA* and *TrackerSci-ATAC* dataset, as well as the medium and mean values for the number of UMIs/genes/unique reads for each cell type.

**Supplementary Table 3:** Differentially expressed genes across newborn cell types. For each gene, the “max.cluster” is the cell type with the highest expression (“max.expr”). The “second.cluster” is the cell type with the second highest expression (“second.expr”). The “fold.change” is the fold change between the max expression and second max expression. The “qval” is the false detection rate (one-sided likelihood ratio test with adjustment for multiple comparisons) for the differential expression test across different cell clusters.

**Supplementary Table 4:** Differentially accessible sites for all newborn cell types. For each gene, the “max.cluster” is the cell type with the highest accessibility (“max.expr”). The “second.cluster” is the cell type with the second highest accessibility (“second.expr”). The “fold.change” is the fold change between the max accessibility and second max accessibility. The “qval” is the false detection rate (one-sided likelihood ratio test with adjustment for multiple comparisons) for the differential accessibility test across different cell clusters. The “is_promoter” indicates whether a site is a promoter or not, and if True, information of corresponding genes is included in “promoter_gene_id”, “promoter_gene_short_name” and “promoter_gene_type”.

**Supplementary Table 5:** Identified linkages between cis-regulatory elements and regulated genes. For each linkage, the “pearson_correlation_coefficient” is Pearson correlation between peak accessibility and gene expression across pseudo-cells. The “region” is either “promoter” or “distal”, indicating whether a site overlaps with the promoter of the linked gene. The “max.cluster.RNA” is the cell type with the highest expression, and the “max.cluster.ATAC” is the cell type with the highest accessibility.

**Supplementary Table 6:** Transcription factors significantly correlated in gene expression and motif accessibility. For each TF, the “PCC” is the Pearson correlation between motif accessibility and gene expression across pseudo-cells. The “max.RNA” is the cell type with the highest gene expression (“max.expr.RNA”). The “second.RNA” is the cell type with the second highest expression (“second.expr.RNA”).

**Supplementary Table 7:** Differentially expressed genes along DG neurogenesis. The “qval” is the false detection rate (one-sided likelihood ratio test with adjustment for multiple comparisons) for the differential test.

**Supplementary Table 8:** Differentially expressed genes along OB neurogenesis. The “qval” is the false detection rate (one-sided likelihood ratio test with adjustment for multiple comparisons) for the differential test.

**Supplementary Table 9:** Differentially accessible sites along DG neurogenesis. The “qval” is the false detection rate (one-sided likelihood ratio test with adjustment for multiple comparisons) for the differential test.

**Supplementary Table 10:** Differentially accessible sites along OB neurogenesis. The “qval” is the false detection rate (one-sided likelihood ratio test with adjustment for multiple comparisons) for the differential test.

**Supplementary Table 11:** Differentially accessible transcription factors along DG neurogenesis. The “qval” is the false detection rate (one-sided likelihood ratio test with adjustment for multiple comparisons) for the differential test.

**Supplementary Table 12:** Differentially accessible transcription factors along OB neurogenesis. The “qval” is the false detection rate (one-sided likelihood ratio test with adjustment for multiple comparisons) for the differential test.

**Supplementary Table 13:** Differentially expressed genes across different age groups for neuronal progenitor cells. For each gene, the “max.group” is the age group with the highest expression (“max.expr”). The “second.group” is the age group with the second highest expression (“second.expr”). The “third.group” is the age group with the minimum expression (“third.expr”). The “qval” is the false detection rate (one-sided likelihood ratio test with adjustment for multiple comparisons) for the differential test. The “promoter_consistent” and “distal_consistent” indicate whether a differentially expressed gene can be supported by its promoter accessibility or its linked distal sites accessibility. The “comments” refers to either “aging_depleted_genes” or “aging_enriched_genes” based on the change of direction.

**Supplementary Table 14:** Differentially expressed genes along oligodendrogenesis. The “qval” is the false detection rate (one-sided likelihood ratio test with adjustment for multiple comparisons) for the differential test.

**Supplementary Table 15:** Differentially accessible sites along oligodendrogenesis. The “qval” is the false detection rate (one-sided likelihood ratio test with adjustment for multiple comparisons) for the differential test.

**Supplementary Table 16:** Differentially accessible transcription factors along oligodendrogenesis. The “qval” is the false detection rate (one-sided likelihood ratio test with adjustment for multiple comparisons) for the differential test.

**Supplementary Table 17:** Differentially expressed genes across different age groups for oligodendrocyte progenitor cells. For each gene, the “max.group” is the age group with the highest expression (“max.expr”). The “second.group” is the age group with the second highest expression (“second.expr”). The “third.group” is the age group with the minimum expression (“third.expr”). The “qval” is the false detection rate (one-sided likelihood ratio test with adjustment for multiple comparisons) for the differential test. The “promoter_consistent” and “distal_consistent” indicate whether a differentially expressed gene can be supported by its promoter accessibility or its linked distal sites accessibility. The “comments” refers to either “aging_depleted_genes” or “aging_enriched_genes” based on the change of direction.

**Supplementary Table 18:** Metadata for human individuals included in this study.

**Supplementary Table 19:** Differentially expressed genes along oligodendrogenesis for human cells. The “qval” is the false detection rate (one-sided likelihood ratio test with adjustment for multiple comparisons) for the differential test.

**Supplementary Table 20:** Differentially expressed genes across regions for each stage along oligodendrogenesis. For each gene, the “max.region” is the region with the highest expression (“max.expr”). The “second.region” is the region with the second highest expression (“second.expr”). The “qval” is the false detection rate (one-sided likelihood ratio test with adjustment for multiple comparisons) for the differential test. The “fold.change” is the fold change between the max expression and second max expression. The “stage” indicates which differentiation stage (i.e, OPC, COP or OLG) the test was performed on.

## Supplementary files

**Supplementary file 1:** Detailed experiment protocols for *TrackerSci-RNA* and *TrackerSci-ATAC*, including all materials and equipment needed, step-by-step descriptions, and representative gel images.

**Supplementary file 2:** Computational pipeline scripts for processing *TrackerSci* data, from sequencer-generated files to single-cell gene count matrix for *TrackerSci-RNA* and single-cell read files for *TrackerSci-ATAC*.

