## Supplementary File 2 for "A comprehensive view of cell-type-specific temporal dynamics in human and mouse brains": Demultiplex_sciRNA.sh

```
run_folder=""
sample_sheet=""
output_folder=""

echo "---------------start demultiplex-----------------------"
echo $(date)
echo "run folder is $run_folder"
echo "sample sheet is $sample_sheet"
echo "output_folder is $output_folder"

# Empty the output folder 
rm -rf $output_folder

#create the output folder:
echo 
echo start making the output_folder
mkdir -p $output_folder/report

# do the demultiplex
/rugpfs/fs0/cao_lab/scratch/jcao/software/anaconda3/bin/bcl2fastq --runfolder-dir $run_folder -o $output_folder --sample-sheet $sample_sheet --reports-dir $output_folder/report --barcode-mismatches 1 --create-fastq-for-index-reads --no-lane-splitting --use-bases-mask Y*,I*,I*,Y* --minimum-trimmed-read-length 0 --mask-short-adapter-reads 0

#remove un-demultiplexed reads
echo remove undetermined reads
rm $output_folder/Undetermined*.fastq.gz

echo "------------------demultiplex done -----------------"
```
