## Supplementary File 2 for "A comprehensive view of cell-type-specific temporal dynamics in human and mouse brains": README.docx

***TrackerSci* Data Processing**

**Step1: Demultiplexing**

We convert sequencer-generated .bcl files into .fastq files using Illumina’s bcl2fastq tolerating one mismatched base in barcodes. Demultiplexing is performed based on the P5 and P7 barcodes.

1. Example master script: Demultiplex_sciRNA.sh
2. Input parameters need to be specified:
   2.1. “run_folder”: the folder containing the raw sequencer-generated data.
   2.2. “sample_sheet”: a .csv file in the sample sheet format as requested for running bcl2fastq. An example is included in the related_files folder.
   2.3. “output_folder”: the complete directory where the demultiplexed .fastq files will be stored.

**Step2: sci-RNA-pipeline/sci-ATAC pipeline**

***TrackerSci-RNA***

From the demultiplexed .fastq files, we followed up with barcode extraction, adaptor trimming, genome alignment, duplicates removal, and we generated the single-cell gene count matrix.

1. Example master script: sciRNA_pipeline.sh
   1.1. All scripts called in the master script can be found in the “Scripts/sciRNA” folder.
2. Input parameters need to be specified:
   2.1. “fastq_folder”: the directory where the demultiplexed .fastq files is stored, same as “output_folder” from Step1.
   2.2. “sample_ID”: a one-column .txt file specifying which PCR samples to be processed line by line, must be a subset of the “Sample_ID” column from the “sample_sheet” file in Step1. An example is included in the “related_files” folder.
   2.3. “all_output_folder”: the complete directory where to output all the intermediate and final files.
   2.4. “Index”: genome index folder used for read alignment with STAR.
   2.5. “gtf_file” : genome annotation file in .gtf format, we used gencode mouse V27 downloaded from <https://ftp.ebi.ac.uk/pub/databases/gencode/Gencode_mouse/release_M27/>.
3. Key output files:
   3.1. Single-cell gene count matrix, including both the exonic, the intronic and the combined matrix, as well as the cell annotation and gene annotation table are in a .RData object in the following directory under “all_output_folder”: “/report/Sci2_Summary.RData”.

***TrackerSci-ATAC***

From the demultiplexed .fastq files, we followed up with barcode extraction, adaptor trimming, genome alignment, duplicates removal, and we generated multiple output formats suitable for downstream analysis, including the single-cell read file, the promoter count matrix, and the snapfile file for SnapATAC processing.

1. Example master script: sciATAC_pipeline.sh
   1.1. All scripts called in the master script are included in the “Scripts/sciATAC” folder.
2. Input parameters need to be specified:
   2.1. “fastq_folder”: the directory where the demultiplexed .fastq files is stored, same as “output_folder” from Step1.
   2.2. “sample_ID”: a one-column .txt file specifying which PCR samples to be processed line by line, must be a subset of the “Sample_ID” column from the “sample_sheet” file in Step1.
   2.3. “all_output_folder”: the complete directory where to output all the intermediate and final files.
   2.4. “Index”: genome index folder used for read alignment with STAR.
   2.5. “gtf_file” : genome annotation file in .gtf format, we used gencode mouse V27 downloaded from <https://ftp.ebi.ac.uk/pub/databases/gencode/Gencode_mouse/release_M27/>.
3. Key output files:
   3.1. Single-cell read files: individual sam files for each single cell are stored in “sam_splitted” under “all_output_folder”.
   3.2. Promoter read-count matrix (±1 kb around TSS): .RData object containing the promoter matrix, cell annotation and promoter annotation in the directory: “/peak_count/summary_count_onlypromoter/sciATAC_summary.RData”.
   3.3. A snapfile format containing the cell by bin matrix counting insertion counts across genome-wide(5000-bp bins) for downstream processing in SnapATAC are in the following directory under “all_output_folder”: “snapfile/SAMPLE.snap”.
