## Supplementary File 2 for "A comprehensive view of cell-type-specific temporal dynamics in human and mouse brains": sciATAC_pipeline.sh

```
# This scATAC-seq pipeline accepts a input folder, a .txt file containing the name of PCR samples to be proceesed and a output directory.

########################################################## MODIFY THIS
fastq_folder=""
sample_ID=""
all_output_folder=""
########################################################## MODIFY THIS

bashrc_location=/ru-auth/local/home/asziraki/projects/AS_20200820_single_cell_pipeline/Scripts/original_pipeline/sci-RNA-seq3_pipeline/.bashrc
source $bashrc_location
conda activate ATAC_pipeline

core=16 # Define number of cores used 
cutoff=200 # the number of unique reads cutoff for splitting single cell
gene_size="mm" # for mouse, use "mm", need to be specified when calling peaks using MACS2
##gene_size="hs" # for human, use "hs", need to be specified when calling peaks using MACS2

# define the location of index files for reads alignment with STAR
# Mouse
index="/ru-auth/local/home/asziraki/projects/AS_20200820_single_cell_pipeline/Raw_data/AS_20200820_genome_files/original_pipeline/Reference_ALL/index/index/STAR/mouse-latest-gencode-release-m27/STAR_mouse_M27"
# Human
# index="/ru-auth/local/home/asziraki/projects/AS_20200820_single_cell_pipeline/Raw_data/AS_20200820_genome_files/original_pipeline/Reference_ALL/index/index/STAR/STAR_hg19_RNAseq/"
# Human + Mouse
# index="/ru-auth/local/home/asziraki/projects/AS_20200820_single_cell_pipeline/Raw_data/AS_20200820_genome_files/original_pipeline/Reference_ALL/index/index/STAR/STAR_hg19_mm10_RNAseq/"

# define the gtf file for gene counting
# Mouse
# gtf_file="/ru-auth/local/home/asziraki/projects/AS_20200820_single_cell_pipeline/Raw_data/AS_20200820_genome_files/original_pipeline/Reference_ALL/index/GTF/gencode.vM12.chr_patch_hapl_scaff.annotation.gtf.gz"
gtf_file="/ru-auth/local/home/asziraki/projects/AS_20200820_single_cell_pipeline/Raw_data/AS_20200820_genome_files/original_pipeline/Reference_ALL/index/GTF/mouse-latest-gencode-release-m27/gencode.vM27.primary_assembly.annotation.gtf.gz"
# Human
# gtf_file="/ru-auth/local/home/asziraki/projects/AS_20200820_single_cell_pipeline/Raw_data/AS_20200820_genome_files/original_pipeline/Reference_ALL/index/GTF/rmchr.gencode.v19.chr_patch_hapl_scaff.annotation.gencode.gtf.gz"
# Human + Mouse
# gtf_file="/ru-auth/local/home/asziraki/projects/AS_20200820_single_cell_pipeline/Raw_data/AS_20200820_genome_files/original_pipeline/Reference_ALL/index/GTF/rmchr.gencode.v19.chr_patch_hapl_scaff.annotation.gencode.vM12.chr_patch_hapl_scaff.annotation.gtf.gz"


########################################################### STEP 1: Trimming, alignment, filter low quality alignments and remove duplicates

################ Barcode extraction
script_folder="/Scripts/sciATAC//"
input_folder=$fastq_folder
output_folder=$all_output_folder/UMI_attach
script=$script_folder/UMI_barcode_attach_gzipped_with_dic_sciATAC_2level.py
echo "Changing the name of the fastq files..."
for sample in $(cat $sample_ID); do echo changing name $sample; mv $input_folder/*$sample*R1*fastq.gz $input_folder/$sample.R1.fastq.gz; mv $input_folder/*$sample*R2*fastq.gz $input_folder/$sample.R2.fastq.gz; done

echo "Attaching barcode...."
mkdir -p $output_folder
################ Define the location of a whitelist of all N5/N7 Tn5 barcodes
RT_barcode_N5=$script_folder/RT_barcodes_N5_RC.txt
RT_barcode_N7=$script_folder/RT_barcodes_N7_RC.txt
python $script $input_folder $sample_ID $output_folder $RT_barcode_N5 $RT_barcode_N7 $core
echo "Barcode transformed"

################# Trimming the reads
echo
echo "Start trimming the read file..."
mkdir -p $all_output_folder/trimmed_fastq
trimmed_fastq=$all_output_folder/trimmed_fastq
for sample in $(cat $sample_ID); do echo trimming $sample; trim_galore $all_output_folder/UMI_attach/$sample.R1.fastq.gz $all_output_folder/UMI_attach/$sample.R2.fastq.gz --paired -a CTGTCTCTTATACACATCT -o $trimmed_fastq; done
echo "All trimmed file generated."

############align the reads with STAR, filter the reads based on q > 30, and remove duplicates based on exactly UMI sequence and tagmentation site
#define the output folder
input_folder=$trimmed_fastq
STAR_output_folder=$all_output_folder/STAR_alignment
filtered_sam_folder=$all_output_folder/filtered_sam
rmdup_sam_folder=$all_output_folder/rmdup_sam
#align read1, read2 to the index file using STAR with default setting
echo "Start alignment using STAR..."
mkdir -p $STAR_output_folder
STAR --genomeDir $index --genomeLoad Remove
#start the alignment
for sample in $(cat $sample_ID); do echo aligning $sample; STAR --runThreadN $core --outSAMstrandField intronMotif --genomeDir $index --readFilesCommand zcat --readFilesIn $input_folder/$sample*R1*.gz $input_folder/$sample*R2*.gz --outFileNamePrefix $STAR_output_folder/$sample --genomeLoad LoadAndKeep; done
#remove the index from the memory
STAR --genomeDir $index --genomeLoad Remove
echo "All alignment done."

################################ #make the filter sam folder, and filter and sort the sam file 
#make the filtered sam folder
echo
echo "Start filter and sort the sam files..."
mkdir -p $filtered_sam_folder
for sample in $(cat $sample_ID); do echo Filtering $sample; samtools view -bh -q 30 -F 4 $STAR_output_folder/$sample*.sam|samtools sort -@ 10 -|samtools view -h -|awk '$3 != "MT" && $3 != "chrM"' - >$filtered_sam_folder/$sample.sam; done

################################## remove duplicates using Picard
echo Remove duplicates
mkdir -p $rmdup_sam_folder
for sample in $(cat $sample_ID); do echo removing duplicates $sample; picard MarkDuplicates INPUT=$filtered_sam_folder/$sample.sam OUTPUT=$rmdup_sam_folder/$sample.bam REMOVE_DUPLICATES=true ASSUME_SORTED=True METRICS_FILE=$rmdup_sam_folder/Picard_metrics_file.txt VALIDATION_STRINGENCY=LENIENT; done

################################# calculate the reads number in each file
report_folder=$all_output_folder/report
mkdir -p $report_folder
fastq_folder=$fastq_folder
trimmed_folder=$trimmed_fastq
alignment=$STAR_output_folder
filtered_sam=$filtered_sam_folder
rm_dup_sam=$rmdup_sam_folder
echo sample,Total reads,Barcode matching,After trimming,Uniquely aligned reads,Filtered reads,After remove duplicates>$report_folder/read_number.csv
for sample in $(cat $sample_ID); do echo calculating $sample; echo $sample,$(expr $(zcat $fastq_folder/$sample.R[0-9].fastq.gz|wc -l) / 4),$(expr $(zcat $all_output_folder/UMI_attach/$sample*.gz|wc -l) / 4),$(expr $(zcat $trimmed_folder/$sample*.gz|wc -l) / 4),$(samtools view $alignment/$sample*.sam|wc -l),$(samtools view $filtered_sam/$sample.sam|wc -l),$(samtools view $rm_dup_sam/$sample.bam|wc -l)>>$report_folder/read_number.csv; done
echo "Read number calculation is done."

################################# generate snap file for downstream processing in SnapATAC
export PATH=/ru-auth/local/home/zlu/software/bin:$PATH
sam_folder=$all_output_folder/rmdup_sam/
snap_file_folder=$all_output_folder/snapfile/
mkdir $snap_file_folder

for sample in $(cat $sample_ID); do echo bam to sam: $sample; samtools view $sam_folder/$sample.bam > $snap_file_folder/$sample.sam; done
tmpsam=$(cat $sample_ID | head -n 1)
grep "@" $all_output_folder/filtered_sam/$tmpsam.sam > $snap_file_folder/merged.sam
for sample in $(cat $sample_ID); do sed -e "s/^/$sample./" $snap_file_folder/$sample.sam >> $snap_file_folder/merged.sam ; done
samtools view -S -b $snap_file_folder/merged.sam > $snap_file_folder/merged.bam
samtools sort -o $snap_file_folder/merged_sorted.bam -n $snap_file_folder/merged.bam
chromosome_length_file=$index/chrNameLength.txt
source /rugpfs/fs0/cao_lab/scratch/zlu/anaconda3/etc/profile.d/conda.sh
conda activate py2
snaptools snap-pre --input-file=$snap_file_folder/merged_sorted.bam --output-snap=$snap_file_folder/SAMPLE.snap --genome-name=mm10 --genome-size=$chromosome_length_file --min-mapq=30 --min-flen=0 --max-flen=1000 --keep-chrm=TRUE --keep-single=FALSE --keep-secondary=FALSE --overwrite=True --min-cov=500 --verbose=True
snaptools snap-add-bmat --snap-file=$snap_file_folder/SAMPLE.snap --bin-size-list 5000 --verbose=True
conda deactivate
#########################

######################### STEP 2: Transform alignment bam files to bed files and combine every file to call peaks with macs2
bashrc_location=/ru-auth/local/home/asziraki/projects/AS_20200820_single_cell_pipeline/Scripts/original_pipeline/sci-RNA-seq3_pipeline/.bashrc
source $bashrc_location
conda activate ATAC_pipeline

input_folder=$all_output_folder/rmdup_sam/
output_folder=$all_output_folder/bed_file/
# transform the bam file to bed files
echo 
echo "Start bam to bed file transformation.."
mkdir $output_folder
for sample in $(cat $sample_ID); do echo transforming $sample; samtools view -bh $input_folder/$sample.bam|bedtools bamtobed -split -i -|sort -k1,1 -k2,2n ->$output_folder/$sample.bed; done
echo "all bed file transformed"
# combine the bed files
combined_bed=$output_folder/bed.combined
cat $output_folder/*.bed |sort -k1,1 -k2,2n >$combined_bed
mv $combined_bed $output_folder/combined.bed
# Call peaks for the bed files
input_file=$output_folder/combined.bed
output_folder=$output_folder/macs2_peaks/
mkdir -p $output_folder
macs2 callpeak -t $input_file --nomodel --keep-dup all --extsize 200 --shift -100 -q 0.1 -B --SPMR -f BED -g $gene_size --outdir $output_folder --call-summits
conda deactivate

############################################## STEP 3: extract promoters from gtf file, combine this with called peaks from macs2
macs_folder=$all_output_folder/bed_file/macs2_peaks
reference_folder=$all_output_folder/reference_folder/
promoter_length_upstream=1000
promoter_length_downstream=1000
python_use="/rugpfs/fs0/cao_lab/scratch/asziraki/anaconda3/bin/python"
merge_peak_dis=0
mkdir -p $reference_folder
promoter_file=$reference_folder/promoter_gtf.bed
script=$script_folder/gtf_to_promoter_bed.r
conda activate ATAC_pipeline_createreference
Rscript $script $gtf_file $promoter_file $promoter_length_upstream $promoter_length_downstream
conda deactivate
conda activate ATAC_pipeline
bedtools sort -i $reference_folder/promoter_gtf.bed | bedtools merge -c 4 -o distinct -i - > $reference_folder/promoter_gtf.overlappingmerged.bed 
promoter_bed=$reference_folder/promoter_gtf.overlappingmerged.bed
output_folder=$reference_folder
merge_file=$output_folder/merged.bed
promoter_file=$output_folder/promoter.bed
# generate a combined bed file
echo Start merging the promoter bed file and the MACS2 peak files
cat $promoter_bed $macs_folder/*peaks.narrowPeak | cut -f 1,2,3 | sortBed -i - |bedtools merge -d $merge_peak_dis -i - >$merge_file
bedtools intersect -a $merge_file -b $promoter_bed -wa -wb > $promoter_file
conda deactivate


################################## STEP 4: count reads in peaks and promoter regions and create final R_Object file 
########################### convert the bam files to samfiles
conda activate ATAC_pipeline
sam_folder=$all_output_folder/rmdup_sam/
for sample in $(cat $sample_ID); do echo bam to sam: $sample; samtools view $sam_folder/$sample.bam >$sam_folder/$sample.sam; done
echo "bam to sam all done."

########################### split the sam file based on the barcode, and mv the result to the report folder
output_folder=$all_output_folder/sam_splitted
echo
echo "Start splitting the sam file..."
R_script=$script_folder/sci3_bash_input_ID_output_core.R
bash_script=$script_folder/sci3_split.sh
########################### At this step, we need a txt file containing all possible N5/N7 barcode combinations
barcodes=$script_folder/N5N7_all_combinations.txt
Rscript $R_script $bash_script $sam_folder $sample_ID $output_folder $core $barcodes $cutoff
cat $output_folder/*sample_list.txt>$output_folder/All_samples.txt
cp $output_folder/All_samples.txt $output_folder/../barcode_samples.txt
# output the report the report/barcode_read_distribution folder
mkdir -p $output_folder/../report/barcode_read_distribution
mv $output_folder/*.txt $output_folder/../report/barcode_read_distribution/
mv $output_folder/*.png $output_folder/../report/barcode_read_distribution/
echo
echo "All sam file splitted."

########################### Count total reads per cell
rm $all_output_folder/total_read_per_cell.csv
for barcode in $(cat $all_output_folder/barcode_samples.txt);do echo $barcode,$(grep -v "@" $all_output_folder/sam_splitted/$barcode.sam | wc -l) >> $all_output_folder/total_read_per_cell.csv ;done


####################### Count reads in peaks
all_output=$all_output_folder/peak_count
peak_count_folder=$all_output/peak_cout/
summary_folder=$all_output/summary_count/
mkdir -p $peak_count_folder
mkdir -p $summary_folder
sam_folder=$all_output_folder/sam_splitted/
sample_list=$all_output_folder/barcode_samples.txt
ref_bed=$merge_file
dis=100 # define the length of extending the reads from the tagmentation site for peak counting

###################### Combine peak count files
script=$script_folder/peak_map_new.py
out_folder=$peak_count_folder
python $script $sample_list $sam_folder $ref_bed $out_folder $dis $core
rm $out_folder/cell_count.MM
rm $out_folder/cell_nread.txt
for sample in $(cat $sample_list); do echo combining $sample; cat $out_folder/$sample.count >>$out_folder/cell_count.MM; rm $out_folder/$sample.count; done
for sample in $(cat $sample_list); do cat $out_folder/$sample.nread.txt >> $out_folder/cell_nread.txt; rm $out_folder/$sample.nread.txt; done


echo All files are combined.
conda deactivate
conda activate ATAC_pipeline_createreference

##################### Post-processing
script=$script_folder/peak_count_summary_no_report.r
Rscript $script $peak_count_folder $reference_folder $summary_folder
echo peak count summary generated.
conda deactivate

#################################### 5th STEP : count only in promoter regions
echo Start promoter counting
conda activate ATAC_pipeline
out_folder=$all_output/peak_cout_promoters/
mkdir -p $out_folder
ref_bed=$promoter_bed

script=$script_folder/peak_map_new.py
python $script $sample_list $sam_folder $ref_bed $out_folder $dis $core
rm $out_folder/cell_count.MM
rm $out_folder/cell_nread.txt
for sample in $(cat $sample_list); do echo combining $sample; cat $out_folder/$sample.count >>$out_folder/cell_count.MM; rm $out_folder/$sample.count; done
for sample in $(cat $sample_list); do cat $out_folder/$sample.nread.txt >> $out_folder/cell_nread.txt; rm $out_folder/$sample.nread.txt; done

conda deactivate
conda activate ATAC_pipeline_createreference
peak_count_folder=$all_output/peak_cout_promoters/
summary_folder=$all_output/summary_count_onlypromoter/
mkdir -p $summary_folder
script=$script_folder/peak_count_summary_no_report_promoter.r
Rscript $script $peak_count_folder $reference_folder $summary_folder
conda deactivate
echo Promoter counting finished

#### Move Star log files
mkdir $all_output_folder/report/Log_files
mv Log.out $all_output_folder/report/Log_files/
mv Log.progress.out $all_output_folder/report/Log_files/
mv Aligned.out.sam $all_output_folder/report/Log_files/
mv _STARtmp $all_output_folder/report/Log_files/
```
