## Supplementary File 2 for "A comprehensive view of cell-type-specific temporal dynamics in human and mouse brains": sciRNA_pipeline.sh

```
# This scRNA-seq pipeline accepts a input folder, a .txt file containing the name of PCR samples to be proceesed and a output directory.

########################################################## MODIFY THIS
fastq_folder=""
sample_ID=""
all_output_folder=""
########################################################## MODIFY THIS

bashrc_location=/ru-auth/local/home/asziraki/projects/AS_20200820_single_cell_pipeline/Scripts/original_pipeline/sci-RNA-seq3_pipeline/.bashrc
source $bashrc_location
conda activate original_pipeline

core=16 # Define number of cores used 
cutoff=200 # the number of unique reads cutoff for splitting single cell
script_folder="//Scripts/sciRNA//"

barcodes=$script_folder/barcode_1836.txt

# define the location of index files for reads alignment with STAR
# Mouse
index="/ru-auth/local/home/asziraki/projects/AS_20200820_single_cell_pipeline/Raw_data/AS_20200820_genome_files/original_pipeline/Reference_ALL/index/index/STAR/mouse-latest-gencode-release-m27/STAR_mouse_M27"
# Human
# index="/ru-auth/local/home/asziraki/projects/AS_20200820_single_cell_pipeline/Raw_data/AS_20200820_genome_files/original_pipeline/Reference_ALL/index/index/STAR/STAR_hg19_RNAseq/"
# Human + Mouse
# index="/ru-auth/local/home/asziraki/projects/AS_20200820_single_cell_pipeline/Raw_data/AS_20200820_genome_files/original_pipeline/Reference_ALL/index/index/STAR/STAR_hg19_mm10_RNAseq/"

# define the gtf file for gene counting
# Mouse
# gtf_file="/ru-auth/local/home/asziraki/projects/AS_20200820_single_cell_pipeline/Raw_data/AS_20200820_genome_files/original_pipeline/Reference_ALL/index/GTF/gencode.vM12.chr_patch_hapl_scaff.annotation.gtf.gz"
gtf_file="/ru-auth/local/home/asziraki/projects/AS_20200820_single_cell_pipeline/Raw_data/AS_20200820_genome_files/original_pipeline/Reference_ALL/index/GTF/mouse-latest-gencode-release-m27/gencode.vM27.primary_assembly.annotation.gtf.gz"
# Human
# gtf_file="/ru-auth/local/home/asziraki/projects/AS_20200820_single_cell_pipeline/Raw_data/AS_20200820_genome_files/original_pipeline/Reference_ALL/index/GTF/rmchr.gencode.v19.chr_patch_hapl_scaff.annotation.gencode.gtf.gz"
# Human + Mouse
# gtf_file="/ru-auth/local/home/asziraki/projects/AS_20200820_single_cell_pipeline/Raw_data/AS_20200820_genome_files/original_pipeline/Reference_ALL/index/GTF/rmchr.gencode.v19.chr_patch_hapl_scaff.annotation.gencode.vM12.chr_patch_hapl_scaff.annotation.gtf.gz"

#define the mismatch rate for removing duplicates:
mismatch=0

#define the bin of python
python_path="/rugpfs/fs0/cao_lab/scratch/asziraki/anaconda3/envs/original_pipeline/bin/"

#define the location of script:
# script_path=/ru-auth/local/home/asziraki/projects/AS_20200820_single_cell_pipeline/Scripts/original_pipeline/sci-RNA-seq2_pipeline/scRNA_seq_pipe

# define the location of the R script for multi-core processing
script_path=$script_folder
R_script=$script_folder/sci3_bash_input_ID_output_core.R


############ UMI attach
# this script take in a input folder, a sample ID, a output folder, a oligo-dT barcode file, a corresponding N5 barcode file, and
# it pass the factors to the python script
input_folder=$fastq_folder
output_folder=$all_output_folder/UMI_attach
script=$script_folder/UMI_barcode_attach_gzipped.py
echo "changing the name of the fastq files..."
for sample in $(cat $sample_ID); do echo changing name $sample; mv $input_folder/$sample*R1*gz $input_folder/$sample.R1.fastq.gz; mv $input_folder/$sample*R2*gz $input_folder/$sample.R2.fastq.gz; done
echo "Attaching barcode and UMI...."
mkdir -p $output_folder
$python_path/python $script $input_folder $sample_ID $output_folder $barcodes $core
echo "Barcode transformed and UMI attached."

################# Trimming the read2
echo
echo "Start trimming the read2 file..."
echo $(date)

trimmed_fastq=$all_output_folder/trimmed_fastq
UMI_attached_R2=$all_output_folder/UMI_attach
bash_script=$script_path/sci3_trim.sh
Rscript $R_script $bash_script $UMI_attached_R2 $sample_ID $trimmed_fastq $core

############align the reads with STAR, filter the reads based on q > 30, and remove duplicates based on exactly UMI sequence and tagmentation site
#define the output folder
input_folder=$trimmed_fastq
STAR_output_folder=$all_output_folder/STAR_alignment
filtered_sam_folder=$all_output_folder/filtered_sam
rmdup_sam_folder=$all_output_folder/rmdup_sam

#align read2 to the index file using STAR with default setting
echo "Start alignment using STAR..."
echo input folder: $input_folder
echo sample ID file: $sample_ID
echo index file: $index
echo output_folder: $STAR_output_folder
#make the output folder
mkdir -p $STAR_output_folder
#remove the index from the memory
STAR --genomeDir $index --genomeLoad Remove
#start the alignment
for sample in $(cat $sample_ID); do echo Aligning $sample;STAR --runThreadN $core --outSAMstrandField intronMotif --genomeDir $index --readFilesCommand zcat --readFilesIn $input_folder/$sample*gz --outFileNamePrefix $STAR_output_folder/$sample --genomeLoad LoadAndKeep; done
#remove the index from the memory
STAR --genomeDir $index --genomeLoad Remove
echo "All alignment done."

#make the filter sam folder, and filter and sort the sam file 
#make the flltered sam folder
echo
echo "Start filter and sort the sam files..."
echo input folder: $STAR_output_folder
echo output folder: $filtered_sam_folder
core_num=5
bash_script=$script_path/sci3_filter.sh
Rscript $R_script $bash_script $STAR_output_folder $sample_ID $filtered_sam_folder $core_num


# make a folder for rmdup_sam_folder, 
# Then for each filtered sam file, remove the duplicates based on UMI and barcode, chromatin number and position
echo
echo "Start removing duplicates..."
echo input folder: $filtered_sam_folder
echo output folder: $rmdup_sam_folder
mkdir -p $rmdup_sam_folder
rmdup_tmp=$rmdup_sam_folder/tmp
mkdir -p $rmdup_tmp
bash_script=$script_path/sci3_rmdup_nomismatch.sh # for removing duplicates only considering exact match
Rscript $R_script $bash_script $filtered_sam_folder $sample_ID $rmdup_tmp $core $mismatch
mv $rmdup_tmp/*.sam  $rmdup_tmp/../
echo "removing duplicates completed.."
echo
echo "Alignment and sam file preprocessing are done."  

################# split the sam file based on the barcode, and mv the result to the report folder
sam_folder=$all_output_folder/rmdup_sam
sample_list=$sample_ID
output_folder=$all_output_folder/sam_splitted
barcode_file=$barcodes
cutoff=$cutoff

echo
echo "Start splitting the sam file..."
echo samfile folder: $sam_folder
echo sample list: $sample_list
echo ouput folder: $output_folder
echo barcode file: $barcode_file
echo cutoff value: $cutoff
mkdir -p $output_folder
for sample in $(cat $sample_list); do echo Now splitting $sample; $python_path/python $script_path/sam_split.py $sam_folder/$sample.sam $barcode_file $output_folder $cutoff; done
cat $output_folder/*sample_list.txt>$output_folder/All_samples.txt
cp $output_folder/All_samples.txt $output_folder/../barcode_samples.txt
# output the report the report/barcode_read_distribution folder
mkdir -p $output_folder/../report/barcode_read_distribution
mv $output_folder/*.txt $output_folder/../report/barcode_read_distribution/
mv $output_folder/*.png $output_folder/../report/barcode_read_distribution/
echo
echo "All sam file splitted."

################### calculate the reads number
fastq_folder=$fastq_folder
trimmed_folder=$trimmed_fastq
UMI_attach=$UMI_attached_R2
alignment=$STAR_output_folder
filtered_sam=$filtered_sam_folder
rm_dup_sam=$rmdup_sam_folder
#split_sam=$parental_folder/splited_sam
report_folder=$all_output_folder/report/read_num
echo
echo "Start calculating the reads number..."
#make the report folder
mkdir -p $report_folder
#calculate the read number and output the read number into the report folder
echo sample,total reads,after filtering barcode,after trimming,uniquely aligned reads,After remove duplicates>$report_folder/read_number.csv
for sample in $(cat $sample_ID); do echo calculating $sample; echo $sample,$(expr $(zcat $fastq_folder/$sample*R2*.gz|wc -l) / 4),$(expr $(zcat $UMI_attach/$sample*R2*.gz|wc -l) / 4),$(expr $(zcat $trimmed_folder/$sample*R2*.gz|wc -l) / 4),$(samtools view $filtered_sam/$sample.sam|wc -l),$(samtools view $rm_dup_sam/$sample.sam|wc -l)>>$report_folder/read_number.csv; done
echo "Read number calculation is done."

################# gene count
# count reads mapping to genes
output_folder=$all_output_folder/report/human_mouse_gene_count/
input_folder=$all_output_folder/sam_splitted
sample_ID=$all_output_folder/barcode_samples.txt
core_number=$core

script=$script_folder/sciRNAseq_count.py
echo "Start the gene count...."
$python_path/python $script $gtf_file $input_folder $sample_ID $core_number

echo "Make the output folder and transfer the files..."
mkdir -p $output_folder
cat $input_folder/*.count > $output_folder/count.MM
rm $input_folder/*.count
cat $input_folder/*.report > $output_folder/report.MM
rm $input_folder/*.report
mv $input_folder/*_annotate.txt $output_folder/
echo "All output files are transferred~"

conda deactivate

################ Post processing
conda activate original_pipeline_final_step
R_script=$script_folder/gene_count_processing_sciRNAseq_exon_intron.R
Rscript $R_script $all_output_folder/report
conda deactivate

mv $all_output_folder/report/Sci3_Summary.RData $all_output_folder/report/Sci2_Summary.RData
mkdir $all_output_folder/report/Log_files
mv Log.out $all_output_folder/report/Log_files/
mv Log.progress.out $all_output_folder/report/Log_files/
mv Aligned.out.sam $all_output_folder/report/Log_files/
mv _STARtmp $all_output_folder/report/Log_files/

############### Convert the Rdata object into mtx files for the convenience of downstream processing in python
source /rugpfs/fs0/cao_lab/scratch/zlu/anaconda3/etc/profile.d/conda.sh
conda activate sc-01
R_script=$script_folder/RData_to_mtx.R
input_folder=$all_output_folder
output_folder=$all_output_folder/report
Rscript $R_script $input_folder $output_folder
conda deactivate
```
