## Supplementary File 2 for "A comprehensive view of cell-type-specific temporal dynamics in human and mouse brains": gtf_to_promoter_bed.r

```
#!/usr/bin/Rscript

library(stringr)
library(dplyr)
library(tidyr)
library(Matrix.utils)
library(Matrix)
options(stringsAsFactors = FALSE)

args = commandArgs(trailingOnly=TRUE)

gtf_file = args[1]
output = args[2]
upstream =  as.numeric(args[3])
downstream = as.numeric(args[4])

cat("read in the table and extract transcript promoters...")
gtf = read.table(gtf_file, sep = "\t")


gtf = gtf %>% filter(V3 == "transcript")

# extract the gene id information from each string
extract_gene_id <- function(str)
    {
    str = str_split(str, ";")[[1]][1]
    str = str_split(str, " ")[[1]][2]
    return(str)
}

cat("extract the gene features...")
gtf["gene"] = sapply(gtf$V9, extract_gene_id)

gtf_pos = gtf %>% filter(V7 == "+")
gtf_neg = gtf %>% filter(V7 == "-")

cat("Generating the promter region...")
# if the stand is in plus, then the first site is the start site -500
# and then extract the promoter region as start site - 500 to start site

gtf_pos["start"] = gtf_pos$V4 - upstream
gtf_pos["end"] = gtf_pos$V4 + downstream

gtf_pos["start"][gtf_pos["start"] < 0] = 0

# if the stand is in minus strand, then the first site is the end site 
# and then add the promoter region as start site to start site + 500

gtf_neg["start"] = gtf_neg$V5 - downstream
gtf_neg["end"] = gtf_neg$V5 + upstream

gtf_neg["start"][gtf_neg["start"] < 0] = 0

# combine the start and end site
gtf_combine = rbind(gtf_pos, gtf_neg) %>% select(c(V1, start, end, gene))
gtf_combine$start <- str_trim(format(gtf_combine$start, scientific = FALSE), side="both")
gtf_combine$end <- str_trim(format(gtf_combine$end, scientific = FALSE), side="both")

cat("Writing the files...")
# output the bed file into the output file
write.table(gtf_combine, file = output, quote = F, col.names = F, row.names = F, sep = "\t")
```
