## Supplementary File 2 for "A comprehensive view of cell-type-specific temporal dynamics in human and mouse brains": peak_count_summary_no_report.r

```
args = commandArgs(trailingOnly=TRUE)
peak_count_folder = args[1]
merged_peak_folder = args[2]
output_folder = args[3]

library(stringr)
library(dplyr)
library(tidyr)
library(Matrix.utils)
library(Matrix)
library(data.table)
library(methods)
options(stringsAsFactors = FALSE)

# first, use the ATAC summary folder for generating the summary cell annotation file
cat("\nGenerate df_cell, df_peak and df_promoter data set...")
# read in the cell annotation file in the peak count data and combine it with the cell annotation file
peak_cell_file = paste(peak_count_folder, "cell_annot.txt", sep = "")
peak_cell = fread(peak_cell_file)
colnames(peak_cell) = c("sample", "id")
cell_annot = peak_cell

cell_nread_file = paste(peak_count_folder, "cell_nread.txt", sep = "")
cell_nread = fread(cell_nread_file)
colnames(cell_nread) = c("id", "total_reads",'mapped_reads','mapped_ratio')
rownames(cell_nread) = cell_nread$id
cell_annot$total_reads = cell_nread[peak_cell$id,'total_reads']

# read in the gene annote file, the gene annotation file in the merged gene bed file, and 
# the promoter file, and then annotate the peak location, and promoter/not
peak_annot_file = paste(peak_count_folder, "peak_annot.txt", sep = "")
peak_annot = fread(peak_annot_file)
colnames(peak_annot) = c("id", "peak")

peak_bed_file = paste(merged_peak_folder, "merged.bed", sep = "")
peak_bed = fread(peak_bed_file)
colnames(peak_bed) = c("chr", "start", "end")

peak_annot = cbind(peak_annot, peak_bed)

promoter_file = paste(merged_peak_folder, "promoter.bed", sep = "")
promoter_bed = fread(promoter_file)
promoter_bed = promoter_bed %>% select(c(V1, V2, V3, V7))
colnames(promoter_bed) = c("chr", "start", "end", "gene")

promoter_bed$peak = paste(promoter_bed$chr, promoter_bed$start, promoter_bed$end, sep = "-")

# read in the sparse matrix object and generate the sparse matrix for read count
df_peak = peak_annot
rownames(df_peak) = peak_annot$peak

df_cell = cell_annot
rownames(df_cell) = cell_annot$sample

df_promoter = promoter_bed

cat("\nStart generating the sparse matrix...")
peak_matrix_file = paste(peak_count_folder, "cell_count.MM", sep = "")
peak_matrix = fread(peak_matrix_file)

peak_count = sparseMatrix(i = peak_matrix$V1, j = peak_matrix$V2, x = peak_matrix$V3)

df_peak = df_peak[1:nrow(peak_count),]
df_cell = df_cell[1:ncol(peak_count),]
df_cell$peak_reads = colSums(peak_count)
df_cell$peak_ratio = df_cell$peak_reads / df_cell$total_reads

rownames(peak_count) = df_peak$peak
colnames(peak_count) = df_cell$sample

output_file = paste(output_folder, "sciATAC_summary.RData", sep ="")
if (file.exists(output_folder) == F)
    dir.create(output_folder)
    
save(df_cell, df_peak, df_promoter, peak_count, file = output_file)
```
