## Supplementary File 2 for "A comprehensive view of cell-type-specific temporal dynamics in human and mouse brains": peak_map.py

```
import itertools
import collections
import numpy as np
import pandas as pd
from multiprocessing import Pool
from multiprocessing import *
import HTSeq
import sys
from functools import partial
import logging
import os
import sys

sample_list = sys.argv[1]
sam_folder = sys.argv[2]
ref_bed = sys.argv[3]
out_folder = sys.argv[4]
dis = int(sys.argv[5])
core_number = int(sys.argv[6])


def peak_count(sample, sam_folder, DHS, sample_ID, dis):
    # define the function for read count for one sam file

    cell_ID = sample_ID.index(sample) + 1

    sam_file = sam_folder + "/" + sample + ".sam"
    count_output = sam_folder + "/" + sample + ".count"
    counts = collections.Counter()
    almnt_file = HTSeq.SAM_Reader(sam_file)
    sam_name = sample


    print("Start read the input file: " + sam_file + "....")
    for alnmt in almnt_file:
        
        if not alnmt.aligned:
            continue
        
        if alnmt.iv.chrom not in DHS.chrom_vectors:
            continue
            
        # extract the end point of the alnmt
        start_site_iv = alnmt.iv
        #print(start_site_iv, start_site_iv.start_d)
        tn_site = HTSeq.GenomicInterval(start_site_iv.chrom, int(start_site_iv.start_d),int(start_site_iv.start_d) + 1)
        tn_span = HTSeq.GenomicInterval(start_site_iv.chrom, max(int(start_site_iv.start_d) - dis, 0),int(start_site_iv.start_d) + dis)
        peak_tn = set()
        for iv, val in DHS[tn_site].steps():
            peak_tn |= val
        if len(peak_tn) != 0:
            # find the exact peak
            #print("exact match: ", peak_tn)
            for p in peak_tn:
                counts[p] += 1/float(len(peak_tn))
        else:
            # find the overlapped peak
            peak_tn = set()
            for iv, val in DHS[tn_span].steps():
                peak_tn |= val

            if (len(peak_tn) > 0):
                #print("adjacent match: ", peak_tn)
                for p in peak_tn:
                    counts[p] += 1/float(len(peak_tn))
            
    with open(count_output, 'w') as count_output:
        for peak in counts:
            line = str(peak) + "," + str(cell_ID) + "," + str(counts[peak]) + "\n"
            count_output.write(line)
    return 0


peak_id_output = out_folder + "peak_annot.txt"
cell_id_output = out_folder + "cell_annot.txt"

# generate the genome object for read count mapping
DHS = HTSeq.GenomicArrayOfSets("auto", stranded=False)
peak_id = 0

bed = open(ref_bed, "r")
if not os.path.exists(out_folder):
    os.makedirs(out_folder)
peak_file = open(peak_id_output, "w")

print("Generate geome peak matrix and annotation...")
for line in bed:
    peak_id += 1
    peak = line.strip().split("\t")
    chro = peak[0]
    start = peak[1]
    end = peak[2]
    peakname = chro + "-" + str(start) + "-" + str(end)
    output_id_line = str(peak_id) + "\t" + peakname + "\n"
    peak_file.write(output_id_line)
    
    DHS[HTSeq.GenomicInterval( chro, int(start), int(end), "." ) ] += str(peak_id)
    
bed.close()
peak_file.close()

# generate the sample list and the cell annotate list
print("Generate cell annotation...")
sample_ID = list(pd.read_csv(sample_list, header=None)[0])
cell_annotat = open(cell_id_output, "w")
# generate the cell ID annotate file
cell_count = 0
for i in sample_ID:
    cell_count += 1
    message = i + "," + str(cell_count) + "\n"
    cell_annotat.write(message)
cell_annotat.close()

# parallele for the functions
print("Data processing in parallel...")
p = Pool(processes = int(core_number))
#print("Processing core number: ", core_number)
func = partial(peak_count, sam_folder = sam_folder, DHS = DHS, sample_ID = sample_ID, dis = dis)
#sciRNAseq_count(sample, input_folder, exons, genes, gene_end)
result = p.map(func, sample_ID)
p.close()
p.join()
print("All analysis done~")
```
