## Supplementary File 2 for "A comprehensive view of cell-type-specific temporal dynamics in human and mouse brains": sci3_bash_input_ID_output_core.R

```
require("BiocParallel")

# accept a input folder, a sample ID file, and a output folder 
# and a core number and then perform trimming for each file

args = commandArgs(trailingOnly=TRUE)

bash_script=args[1]
input_folder=args[2]
sample_ID=args[3]
output_folder=args[4]
core = as.numeric(args[5])

cat("script: ", bash_script)
cat("\nintput folder: ", input_folder)
cat("\nsample ID: ", sample_ID)
cat("\noutput folder: ", output_folder)
cat("\ncore number: ", core)

others = " "

if (length(args) > 5){
    others = paste(args[6:length(args)], collapse = " ")
    cat("\nother arguments: ", others[1])
    cat("\n")
}


function_sample <- function(input_folder, sample, output_folder, others) {
    trim_c <- paste("bash", bash_script, input_folder, sample, output_folder, others, sep = " ") 
    cat("\n")
    cat(trim_c)
    cat("\n")
    result = system(trim_c, intern = T)
    cat(result)
}


sample_name = (read.csv(sample_ID, header = F))$V1

# make the output folder
if(!dir.exists(output_folder)) 
{
    dir.create(output_folder, recursive = T)
    }

result = bplapply(sample_name, function(x) {function_sample(input_folder, x, output_folder, others)}, 
                  BPPARAM = MulticoreParam(workers = core))

cat("\nAll sample are processed.")
```
