## Supplementary File 2 for "A comprehensive view of cell-type-specific temporal dynamics in human and mouse brains": sci3_split.sh

```
input_folder=$1
sample=$2
output_folder=$3
barcode_file=$4
cutoff=$5

mismatch=1
python="/rugpfs/fs0/cao_lab/scratch/asziraki/anaconda3/envs/ATAC_pipeline/bin/python2.7"
python_script="/ru-auth/local/home/asziraki/projects/AS_20200820_single_cell_pipeline/Scripts/bulkATAC/scripts/peak_count_sciATACseq/sam_split.py"

$python $python_script $input_folder/$sample.sam $barcode_file $output_folder $cutoff
echo splitting sample done: $sample
```
