## Supplementary File 2 for "A comprehensive view of cell-type-specific temporal dynamics in human and mouse brains": UMI_barcode_attach_gzipped_with_dic_sciATAC_2level.py

```
import subprocess
import sys
from Levenshtein import distance
import gzip
from multiprocessing import Pool
from functools import partial
import pickle

'''
    this script accept a read1 file, a read2 file, a read3 file, a output_file, a ligation barcode list,
    an RT barcode list,
    and mismatch rate, then it open the read1, read2 and read3, output file,
    then extract the RT barcode and UMI sequence in the read 1 file and the ligation barcode from the read 2 file, and convert the
    barcode to the real barcode in the barcode list based on the mismatch rate (this step does not happen),
    then it attach the barcodes and UMI sequence to the read name of the read3 file
'''    
    
def UMI_attach_read2_barcode_list(sample, input_folder, output_folder, RT_barcode_list_N5, RT_barcode_list_N7, mismatch_rate = 1):
    #open the read1, read2, and output file
    Read1 = input_folder + "/" + sample + ".R1.fastq.gz"
    Read2 = input_folder + "/" + sample + ".R2.fastq.gz"
    output_file1 = output_folder + "/" + sample + ".R1.fastq.gz"
    output_file2 = output_folder + "/" + sample + ".R2.fastq.gz"
    mismatch_rate = int(mismatch_rate)
    f1 = gzip.open(Read1)
    f2 = gzip.open(Read2)
    f3 = gzip.open(output_file1, 'wb')
    f4 = gzip.open(output_file2, 'wb')
  
    
    line1 = f1.readline()
    line2 = f2.readline()
    total_line = 0
    filtered_line = 0
    
    while (line1):
        total_line += 1
        line1_header = line1
        line2_header = line2
        line1 = f1.readline()
        line2 = f2.readline()
        # first check if the Tagmentaion Read1 barcode match with the barcode
        BC1 = line1[0:6]

        if BC1 in RT_barcode_list_N5:
            
            BC1_match = BC1
            # check Tagmentation Read2 barcode
            BC2 = line2[0:6]

            if BC2 in RT_barcode_list_N7:
                BC2_match = BC2
                filtered_line += 1
                first_line = '@' + BC1_match + BC2_match + ',' + line1_header[1:]
                f3.write(first_line)
                first_line = '@' + BC1_match + BC2_match + ',' + line2_header[1:]
                f4.write(first_line)
                
                second_line = line1[25:]
                f3.write(second_line)
                second_line = line2[25:]
                f4.write(second_line)
                
                third_line = f1.readline()
                f3.write(third_line)
                third_line = f2.readline()
                f4.write(third_line)
                
                four_line = f1.readline()
                four_line2 = four_line[25:]
                f3.write(four_line2)
                four_line = f2.readline()
       	       	four_line2 = four_line[25:]
                f4.write(four_line2)
                
                line1 = f1.readline()
                line2 = f2.readline()
            
            else:
                line1 = f1.readline()
                line1 = f1.readline()
                line1 = f1.readline()
                line2 = f2.readline()
                line2 = f2.readline()
                line2 = f2.readline()
                
        else:
            line1 = f1.readline()
            line1 = f1.readline()
            line1 = f1.readline()
            line2 = f2.readline()
            line2 = f2.readline()
            line2 = f2.readline()


    f1.close()
    f2.close()
    f3.close()
    f4.close()
    print("sample name: %s, total line: %f, filtered line: %f, filter rate: %f" 
          %(sample, total_line, filtered_line, float(filtered_line) / float(total_line)))

# this function accept an input folder and a output folder and then generate the output file with the index
def attach_UMI_files(input_folder, sampleID, output_folder, RT_barcode_file_N5, RT_barcode_file_N7, core):
    
    init_message = '''
    --------------------------start attaching UMI-----------------------------
    input folder: %s
    sample ID: %s
    output_folder: %s
    RT barcode file N5: %s
    RT barcode file N7: %s
    ___________________________________________________________________________
    ''' %(input_folder, sampleID, output_folder, RT_barcode_file_N5, RT_barcode_file_N7)
    
    print(init_message)
    
    
    print("Load RT barcode dictionary...")
    
    # generate the N5 RT barcode list:
    barcodes_N5 = open(RT_barcode_file_N5, "rb")
    with barcodes_N5 as f:
        RT_barcode_list_N5 = f.read().splitlines()
    barcodes_N5.close()

    # generate the N7 RT barcode list:
    barcodes_N7 = open(RT_barcode_file_N7, "rb")
    with barcodes_N7 as f:
        RT_barcode_list_N7 = f.read().splitlines()
    barcodes_N7.close()

    #for each sample in the sample list, use the read1 file, read2 file, output file
    # and barcode_list to run UMI_attach_read2_barcode_list
    sample_file = open(sampleID)
    sample_list = []
    for line in sample_file:
        sample = line.strip()
        sample_list.append(sample)
    sample_file.close()
    
    # parallele for the functions
    p = Pool(processes = int(core))
    #print("Processing core number: ", core_number)
    func = partial(UMI_attach_read2_barcode_list, input_folder = input_folder, output_folder=output_folder, RT_barcode_list_N5=RT_barcode_list_N5, RT_barcode_list_N7=RT_barcode_list_N7, mismatch_rate = 1)
    #sciRNAseq_count(sample, input_folder, exons, genes, gene_end)
    result = p.map(func, sample_list)
    p.close()
    p.join()
    
    #print the completion message
    com_message = '''~~~~~~~~~~~~~~~UMI attachment done~~~~~~~~~~~~~~~~~~'''
    print(com_message)
    
if __name__ == "__main__":
    input_folder = sys.argv[1]
    sampleID = sys.argv[2]
    output_folder = sys.argv[3]
    RT_barcode_file_N5 = sys.argv[4]
    RT_barcode_file_N7 = sys.argv[5]
    core=sys.argv[6]
    attach_UMI_files(input_folder, sampleID, output_folder,  RT_barcode_file_N5, RT_barcode_file_N7, core)
```
