## Supplementary File 2 for "A comprehensive view of cell-type-specific temporal dynamics in human and mouse brains": gene_count_processing_sciRNAseq_exon_intron.R

```
args = commandArgs(trailingOnly=TRUE)

# define the report folder from sci-RNA-seq pipeline
report_folder = args[1]

# define the output folder for output the df_cell, df_gene and gene_count matrix
output_folder = args[1]

suppressMessages(library(Matrix))
suppressMessages(library(tidyverse))
suppressMessages(library(data.table))


combine_exon_intron <- function (df_gene, gene_count) 
{
    gene_count_exon = gene_count[df_gene$exon_intron == "exon", 
        ]
    gene_count_intron = gene_count[df_gene$exon_intron == "intron", 
        ]
    if (nrow(gene_count_exon) == nrow(gene_count_intron)) {
        gene_count_combine = gene_count_exon + gene_count_intron
    }
    else {
        gene_count_combine = gene_count_exon[-nrow(gene_count_exon), 
            ] + gene_count_intron
        gene_count_combine = rbind(gene_count_combine, gene_count_exon[nrow(gene_count_exon), 
            ])
        gene_count_intron = as(rbind(gene_count_intron, 0), "sparseMatrix")
    }
    return(list(gene_count_combine, gene_count_exon, gene_count_intron))
}

sciRNAseq_gene_count_summary <- function (gene_count_folder) {
    gene_matrix = paste(gene_count_folder, "/count.MM", sep = "")
    df_gene = paste(gene_count_folder, "/gene_name_annotate.txt", 
        sep = "")
    df_cell = paste(gene_count_folder, "/cell_annotate.txt", 
        sep = "")
    df_report = paste(gene_count_folder, "/report.MM", sep = "")
    report_annotate = paste(gene_count_folder, "/report_annotate.txt", 
        sep = "")
    df_gene = read.csv(df_gene, header = F)
    df_cell = read.csv(df_cell, header = F)
    gene_matrix = read.csv(gene_matrix, header = F)
    colnames(df_gene) = c("gene_id", "gene_type", "exon_intron", 
        "gene_name", "index")
    colnames(df_cell) = c("sample", "index")
    rownames(df_gene) = df_gene$gene_id
    rownames(df_cell) = df_cell$cell_name
    gene_count = sparseMatrix(i = gene_matrix$V1, j = gene_matrix$V2, 
        x = gene_matrix$V3)
    df_gene = df_gene[1:nrow(gene_count), ]
    rownames(gene_count) = df_gene$gene_id
    colnames(gene_count) = df_cell$cell_name
    combine = combine_exon_intron(df_gene, gene_count)
    gene_count = combine[[1]]
    df_gene = df_gene %>% filter(exon_intron == "exon")
    reportMM = read.csv(df_report, header = F)
    df_report = sparseMatrix(i = reportMM$V1, j = reportMM$V2, 
        x = reportMM$V3)
    df_report = as.matrix(t(df_report))
    df_report_annotate = read.csv(report_annotate, header = F)
    colnames(df_report) = df_report_annotate$V2
    df_report = data.frame(df_report)
    df_report["index"] = as.numeric(rownames(df_report))
    df_cell_combine = inner_join(df_cell, df_report, by = "index")
    df_cell_combine["all_exon"] = df_cell_combine$X.Perfect.intersect.exon.match + 
        df_cell_combine$X.Nearest.intersect.exon.match + df_cell_combine$X.Perfect.combine.exon.match + 
        df_cell_combine$X.Nearest.combine.exon.match
    df_cell_combine["all_intron"] = df_cell_combine$X.Perfect.intersect.gene.match + 
        df_cell_combine$X.Nearest.intersect.gene.match + df_cell_combine$X.Perfect.combine.gene.match + 
        df_cell_combine$X.Nearest.combine.gene.match
    df_cell_combine["all_reads"] = df_cell_combine$all_exon + 
        df_cell_combine$all_intron + df_cell_combine$X.No.match
    df_cell_combine["unmatched_rate"] = df_cell_combine$X.No.match/df_cell_combine$all_reads
    df_cell = df_cell_combine %>% select(sample, unmatched_rate)
    df_cell$UMI_count = df_cell_combine$all_exon + df_cell_combine$all_intron - df_cell_combine$X.ambiguous.match.for.exons - df_cell_combine$X.ambiguous.match.for.genes
    df_gene = df_gene %>% select(gene_id, gene_type, gene_name)
    return(list(df_cell, df_gene, gene_count, combine[[2]], combine[[3]]))
}
result = sciRNAseq_gene_count_summary(paste0(report_folder, "/human_mouse_gene_count/"))
df_cell = result[[1]]
df_gene = result[[2]]
gene_count_all = result[[3]]
gene_count_exon = result[[4]]
gene_count_intron = result[[5]]
save(df_cell, df_gene, gene_count_all, gene_count_exon, gene_count_intron, file = paste0(output_folder, "/Sci3_Summary.RData"))
```
