## Supplementary File 2 for "A comprehensive view of cell-type-specific temporal dynamics in human and mouse brains": RData_to_mtx.R

```
library(tidyverse)
library(Matrix)
library(ggplot2)

args = commandArgs(trailingOnly=TRUE)

input_folder = args[1]
output_folder = args[2]

cat("\nintput folder: ", input_folder)
cat("\noutput folder: ", output_folder)

if(!file.exists(output_folder)) {
    dir.create(output_folder)
}

load(file.path(input_folder, "report/Sci2_Summary.RData"))


df_cell$Gene_count = colSums(gene_count_all > 0)
df_gene$Human_genes = (str_sub(df_gene$gene_id, 1, 4) == "ENSG")
df_cell$Human_UMI = colSums(gene_count_all[df_gene$Human_genes, ])
df_cell$Mouse_UMI = colSums(gene_count_all[!df_gene$Human_genes, ])
df_cell$Human_gene = colSums(gene_count_all[df_gene$Human_genes, ] > 0)
df_cell$Mouse_gene = colSums(gene_count_all[!df_gene$Human_genes, ] > 0)
df_cell$Human_cells = (df_cell$Human_UMI / df_cell$UMI_count > 0.9)

tmp = df_cell %>% separate(sample, c("sample", "bc"), sep = "\\.")
df_cell$RT_barcode = tmp$bc

writeMM(obj = gene_count_all,file = file.path(output_folder,"genecount.mtx"))
write.csv(df_cell,file.path(output_folder,"df_cell.csv"))
write.csv(df_gene,file.path(output_folder,"df_gene.csv"))
```
