## Supplementary File 2 for "A comprehensive view of cell-type-specific temporal dynamics in human and mouse brains": sam_split.py

```
'''
in this script, I will read into the sam file and the barcode file, and then count the reads number per
barcode
'''
import sys
import matplotlib as mpl
mpl.use('Agg')
import matplotlib.pyplot as plt
from matplotlib.backends.backend_pdf import PdfPages

def samfile_barcode_count(sam_file, barcode_file):
    
    #generate the barcode list and barcode dictionary
    barcodes = open(barcode_file)
    barcode_ls = []
    barcode_dic = {}
    for line in barcodes:
        barcode = line.strip()
        barcode_ls.append(barcode)
        barcode_dic[barcode] = 0
    barcodes.close()
    #read the sam file, and count the number per barcode
    sam = open(sam_file)
    for line in sam:
        if (line[0] == '@'):
            continue
        else:
            name = (((line.split('\t'))[0]).split(','))
            barcode = name[0]
            barcode_dic[barcode] += 1
    sam.close()
    return barcode_dic

def split_samfile(sam_file, barcode_file, output_folder, cutoff):
    '''
    this script accept a sam file, a barcode file, a output_file, a cutoff value,
    then it will call the samfile_barcode_count function and get the total read count per barcode,
    then it use the cutoff value to filter the barcode,
    and generate the output samfile for single cells, generate the sample_ID.txt in the output folder,
    generate the reads distribution in the output folder/read_distribution_barcode;
    '''
    
    # generate the count per barcode
    barcode_count = samfile_barcode_count(sam_file, barcode_file)
    
    # plot the barcode reads distribution and save the result to the ouput folder
    plot_name = (sam_file.split('/')[-1]).split('.')[0]
    fig = plt.figure()
    plt.hist(barcode_count.values(), bins=100)
    plt.ylabel('frequency')
    plt.xlabel('Number of unique reads')
    fig_output = output_folder + '/' + plot_name + '.png'
    
    fig.savefig(fig_output)

    #also output the barcode number and distribution to the output folder
    read_dist = open(output_folder + '/' + plot_name + '.txt', 'w')
    for barcode in barcode_count:
        line = barcode + ', %d\n' %(barcode_count[barcode])
        read_dist.write(line)
    read_dist.close()

    #Generate the read distribution in the output foler/read_distribution_barcode
    
    #filter the barcode based on the cutoff value
    barcode_filtered = []
    for barcode in barcode_count:
        if barcode_count[barcode] >= cutoff:
            barcode_filtered.append(barcode)
    #print barcode_filtered    
    #generate the output sam file and sample_list file
    sample_list_file = open(output_folder + '/' + plot_name + '.' + 'sample_list.txt', 'w')
    output_files = {}
    for barcode in barcode_filtered:
        # output_file = output_folder + '/' + plot_name + '.' + barcode + '.sam'
        # output_files[barcode] = open(output_file, 'w')
        sample_list_file.write(plot_name + '.' + barcode + '\n')
     
    # output the each read to the output sam file
    sam = open(sam_file)
    for line in sam:
        if (line[0] == '@'):
            for barcode in barcode_filtered:
                output_file = output_folder + '/' + plot_name + '.' + barcode + '.sam'
                output_files[barcode] = open(output_file, 'a')
                output_files[barcode].write(line)
                output_files[barcode].close()
        else:
            barcode = (((line.split('\t'))[0]).split(','))[0]
            if barcode in barcode_filtered:
                output_file = output_folder + '/' + plot_name + '.' + barcode + '.sam'
                output_files[barcode] = open(output_file, 'a')
                output_files[barcode].write(line)
                output_files[barcode].close()
    
    #close the files:
    sample_list_file.close()
    sam.close()
    # for barcode in barcode_filtered:
        # output_files[barcode].close()

if __name__ == '__main__':
    sam_file = sys.argv[1]
    barcode_file = sys.argv[2]
    output_folder = sys.argv[3]
    cutoff = int(sys.argv[4])
    split_samfile(sam_file, barcode_file, output_folder, cutoff)
```
