## Supplementary File 2 for "A comprehensive view of cell-type-specific temporal dynamics in human and mouse brains": sci3_filter.sh

```
input_folder=$1
sample=$2
output_folder=$3

echo Filtering sample: $sample
samtools view -bh -q 30 -F 4 $input_folder/$sample*.sam|samtools sort -@ 10 -|samtools view -h ->$output_folder/$sample.sam
echo Filtering $sample done.
```
