## Supplementary File 2 for "A comprehensive view of cell-type-specific temporal dynamics in human and mouse brains": sci3_rmdup_nomismatch.sh

```
input_folder=$1
sample=$2
output_folder=$3
mismatch=$4

python="/rugpfs/fs0/cao_lab/scratch/asziraki/anaconda3/envs/original_pipeline/bin/python2.7"
python_script="/ru-auth/local/home/asziraki/projects/AS_20200820_single_cell_pipeline/Scripts/original_pipeline/sci-RNA-seq3_pipeline/script_folder/rm_dup_barcode_UMI_no_mismatch.py"

echo Filtering sample: $sample

$python $python_script $input_folder/$sample.sam $output_folder/$sample.sam $mismatch

echo Filtering $sample done.
```
