## Supplementary File 2 for "A comprehensive view of cell-type-specific temporal dynamics in human and mouse brains": sci3_trim.sh

```
input_folder=$1
sample=$2
output_folder=$3

echo Trimming sample: $sample
trim_galore $input_folder/$sample*.gz -a AAAAAAAA --three_prime_clip_R1 1 -o $output_folder
module unload python/2.7.3
echo Trimming $sample done.
```
