## Supplementary File 2 for "A comprehensive view of cell-type-specific temporal dynamics in human and mouse brains": UMI_barcode_attach_gzipped.py

```
"""
Created on Tue Apr  5 22:16:54 2016

@author: Junyue
"""

import subprocess
import sys
from Levenshtein import distance
import gzip
from multiprocessing import Pool
from functools import partial

'''
    this script accept a read1 file, a read2 file, a output_file, a oligodT, barcode list
    and mismatch rate, then it open the read1 and read2, output file,
    then extract the barcode and UMI sequence in the read 1 file, and convert the
    barcode to the real barcode in the barcode list based on the mismatch rate,
    then it attach the barcode and UMI sequence to the read name of the read2 file
'''

def UMI_attach_read2_barcode_list(sample, input_folder, output_folder, barcode_list, mismatch_rate = 1):
    #open the read1, read2, and output file
    Read1 = input_folder + "/" + sample + ".R1.fastq.gz"
    Read2 = input_folder + "/" + sample + ".R2.fastq.gz"
    output_file = output_folder + "/" + sample + ".R2.fastq.gz"
    mismatch_rate = int(mismatch_rate)
    f1 = gzip.open(Read1)
    f2 = gzip.open(Read2)
    f3 = gzip.open(output_file, 'wb')
    
    line1 = f1.readline()
    line2 = f2.readline()

    while (line1):
        line1 = f1.readline()
        target = line1[8:18]
        
        for barcode in barcode_list:
            mismatch = distance(barcode, target)
            find = False
            
            if (mismatch <= mismatch_rate):
                find = True
                UMI = line1[:8]
                first_line = '@' + barcode + ',' + UMI + ',' + line2[1:]
                f3.write(first_line)

                second_line = f2.readline()
                f3.write(second_line)

                third_line = f2.readline()
                f3.write(third_line)

                four_line = f2.readline()
                f3.write(four_line)

                line2 = f2.readline()
                break
                
        if find == False:
            line2 = f2.readline()
            line2 = f2.readline()
            line2 = f2.readline()
            line2 = f2.readline()

        line1 = f1.readline()
        line1 = f1.readline()
        line1 = f1.readline()

    f1.close()
    f2.close()
    f3.close()

# this function accept an input folder and a output folder and then generate the output file with the index
def attach_UMI_files(input_folder, sampleID, output_folder, barcode_file, core_number):
    
    init_message = '''
    --------------------------start attaching UMI-----------------------------
    input folder: %s
    sample ID: %s
    output_folder: %s
    barcode file: %s
    ___________________________________________________________________________
    ''' %(input_folder, sampleID, output_folder, barcode_file)
    
    print(init_message)
    
    # generate the barcode list:
    barcode_list = []
    barcodes = open(barcode_file)
    for barcode in barcodes:
        barcode_list.append(barcode.strip())
    barcodes.close()
    
    #for each sample in the sample list, use the read1 file, read2 file, output file
    # and barcode_list to run UMI_attach_read2_barcode_list
    sample_file = open(sampleID)
    sample_list = []
    for line in sample_file:
        sample = line.strip()
        sample_list.append(sample)
    sample_file.close()
    
    # parallele for the functions
    p = Pool(processes = int(core_number))
    #print("Processing core number: ", core_number)
    func = partial(UMI_attach_read2_barcode_list, input_folder = input_folder, output_folder=output_folder, barcode_list=barcode_list, mismatch_rate = 1)
    #sciRNAseq_count(sample, input_folder, exons, genes, gene_end)
    result = p.map(func, sample_list)
    p.close()
    p.join()
    
    #print the completion message
    com_message = '''~~~~~~~~~~~~~~~UMI attachment done~~~~~~~~~~~~~~~~~~'''
    print com_message
    
if __name__ == "__main__":
    input_folder = sys.argv[1]
    sampleID = sys.argv[2]
    output_folder = sys.argv[3]
    barcode_file = sys.argv[4]
    core=sys.argv[5]
    attach_UMI_files(input_folder, sampleID, output_folder, barcode_file, core)
```
