## Supplementary File 1 for "A comprehensive view of cell-type-specific temporal dynamics in human and mouse brains"

### TrackerSci protocol

#### Materials used

- EZ lysis Buffer (Millipore Sigma, NUC101)
- SUPERase•In™ RNase Inhibitor (Thermo Fisher Scientific, AM2696)
- cOmplete™, EDTA-free Protease Inhibitor Cocktail (Sigma, 11873580001)
- PBS (VWR, 45000-446)
- BSA (NEB, B90000S)
- 1 M Tris-HCl, pH 7.5 (VWR, 97062-936)
- 5 M NaCl (VWR, 97062-858)
- 1 M $MgCl_2$ (VWR, 97062-848)
- Nuclease-free water (VWR, 45001-044)
- 16% methanol-free formaldehyde (Thermo Fisher Scientific, 28906)
- Click-iT Plus EdU Alexa Fluor™ 647 Flow Cytometry assay Kit (Thermo Fisher Scientific, 10634)
- DAPI (Invitrogen, D1306)
- 10 mM dNTP mix (Thermo Fisher Scientific, R0194)
- 100 mM DTT (Invitrogen, P2325)
- SuperScript™ IV Reverse Transcriptase (Invitrogen, 18090200)
- RNaseOUT™ Recombinant Ribonuclease Inhibitor (Invitrogen, 10777019)
- 0.5 M EDTA (VWR, 19K1956689)
- mRNA Second Strand Synthesis buffer and enzyme (NEB, E6111L)
- Dimethylformamide (Fisher, AC327175000)
- 10% SDS (VWR, E719-100ML)
- Tween-20 (Sigma, P9416)
- NEBNext High-Fidelity 2X PCR Master Mix (NEB, M0541L)
- 100X dsGreen for real-time PCR (Lumiprobe, 41010)
- AMPure XP beads (Beckman Coulter, A63882)
- Ethanol
- EB buffer (Qiagen, 19086)
- DNA Clean & Concentrator kit (Zymoresearch, D4014)
- 2% E-Gel™ EX Agarose Gels (Invitrogen, G402022)
- E-Gel™ 50 bp DNA Ladder (Invitrogen, 10488099)
- Eppendorf™ DNA LoBind Tubes (1.5 mL) (Eppendorf, 22431021)
- Fine scissors (Fine Science Tools, 14060-09)
- pluriStrainer Mini 20 µm filter (Pluriselect, 43-10020-70)
- 40 µm cell strainers (VWR, 470236-276)
- 6-cm/10-cm cell culture dishes(Genesee, 25-260/25-202)
- 5 mL Syringes (Fisher, 309603)
- 96-well plates (BioRad, HSP9601)

#### Required Equipments

- Bioruptor Sonication Device
- Centrifuge (Eppendorf 5702 RH)
- Eppendorf Mastercycler (4x)

- Freezer (-20C, -80C) and Refrigerator (4C)
- Gel Imager
- Ice Buckets
- Multi-channel Pipettes (2-20μL, 20-200μL) (Rainin Instruments)
- Pipettors
- Liquid nitrogen tank for sample storage

#### *TrackerSci-RNA*

#### Buffers Preparation

1. Nuclei Isolation Buffer (NIB):
    a. EZ lysis Buffer + 0.1% SUPERase•In™ RNase Inhibitor + 1X Protease Inhibitor
        i. To make 25X Protease Inhibitor, dissolve 1 tablet of cOmplete™, EDTA-free Protease Inhibitor Cocktail into 2 mL nuclease free water
    b. Made fresh every time, 5 mL per sample, stored on ice
2. Nuclei Buffer (NB)
    a. 10 mM Tris-HCl pH7.5, 10 mM NaCl, 3 mM $MgCl_2$ in nuclease-free water
    b. Made 500 mL and can be stored in 4°C for long term

| Reagent | Stock Con. | Final Con. | Volume (mL) |
|---|---|---|---|
| Tris-HCl pH7.5 | 1 M | 10 mM | 5 |
| NaCl | 5 M | 10 mM | 1 |
| $MgCl_2$ | 1 M | 3 mM | 1.5 |
| Nuclease-free water | | | 492.5 |
| Total | | | 500 |

3. Nuclei Suspension Buffer (NSB-RNA):
    a. NB + 1% BSA + 0.1% SUPERase•In™ RNase Inhibitor
    b. Made fresh every time
4. 1% Formaldehyde
    a. 16% methanol-free formaldehyde in PBS
    b. made fresh every time, 1.2 mL per sample, stored on ice
5. 2X Tagmentation buffer
    a. 20 mM Tris-HCl pH 7.5 + 20 mM $MgCl_2$ + 20% Dimethylformamide (Fisher AC327175000)
    b. Can be stored in -20°C for long term

#### EdU labeling and nuclei isolation on cultured cells

6. Passage cells (HEK293T and NIH/3T3) in 10 cm dishes to 60% confluency at the time of labeling. Incubate cells at 37°C with 5% $CO_2$ in high glucose DMEM supplemented with 10% FBS and 1x Penicillin- Streptomycin.
7. Add EdU at the final concentration of 10 μM and place cells in the incubator for 1 hour.

8. Harvest cells with 2 mL trypsin and incubate cells at 37°C for 30 seconds. Neutralize trypsin with 8 mL of DMEM and resuspend until no visible cell clumps can be observed.
9. Combine HEK293T and 3T3 cells at 1:1 ratio and pellet at 300g for 5 minutes. Dump/aspirate the supernatant.
10. Resuspend the cells in 10 mL ice-cold PBS and pellet at 300g for 5 minutes. Dump/aspirate the supernatant.
11. Resuspend the pellet in 1mL NIB and transfer cells to a 1.5 mL LoBind tube. Pellet the cells at 300g for 5 minutes and dump supernatant.

#### EdU labeling and nuclei isolation on mouse brain tissues

12. Perform i.p. injection on mice with 50 mg/kg of EdU in PBS in 24-hour intervals for five days.
13. Harvest mice brains and flash freeze in liquid nitrogen before storing in -80°C. If used immediately after harvesting, brains may be used directly after.
14. Thawed brains were cut into small pieces (<1mm diameter) with fine scissors in 1 mL ice-cold PBS with 1% SUPERase•In™ RNase Inhibitor and 1% BSA, and transferred to 1.5 mL tubes.
15. Pellet tissue pieces at 500g, 4°C for 5 minutes, and resuspend in 750 µL NIB with a 1 mL pipette with wide bore tips. Incubate samples on ice for 5 minutes.
16. Homogenize the tissue through a 40 µm cell strainer into a 6 cm dish with the rubber tip of a 5 mL syringe. Add an additional 750 µL of NIB to aid with homogenization.
17. Transfer the filtered tissue into a LoBind 1.5ml tube and pellet the nuclei (500g, 5 minutes), then dump the supernatant.

#### Nuclei Fixation

18. Resuspend nuclei pellets in 1mL of freshly prepared 1% formaldehyde and fix cells on ice for 10 minutes. (In a chemical hood)
19. Pellet the nuclei at 500g for 5 minutes, dump supernatant and resuspend in 500 µL NIB.
20. Pellet the nuclei at 500g for 5 minutes, dump supernatant and resuspend in 400 µL NIB.
21. If working with tissue, perform a short sonication (Diagenode, low power mode for 12 seconds) to reduce clumping.
22. Filter all nuclei with a pluriStrainer Mini 20 µm filter with a quick spin down (15 seconds), then pellet the nuclei at 500g for 5 minutes, and resuspend with 100 µL NSB-RNA with a 1 mL pipette.

#### Click-iT Tagging

23. Prepare Click-iT reaction cocktail according to the manufacturer

| Reaction component | Volume (µL) |
|---|---|
| PBS | 433 |
| Copper protectant | 10 |
| Azide solution | 2.5 |
| 1X Reaction Buffer addictive | 50 |

| | |
|---|---|
| SUPERase•In™ RNase Inhibitor | 5 |
| Total volume | 500 |

24. Add 500 μL of Click-iT reaction cocktail to 100 μL of resuspend nuclei and mix well.
25. Incubate the reaction mixture for 30 minutes at room temperature, protected from light.
26. Pellet the nuclei at 500g for 5 minutes and wash the cells once with 500 μl of 1X Click-iT® saponin-based permeabilization and wash reagent
27. Pellet the cells, remove the supernatant, and resuspend the cells in 200 μl NSB-RNA. Add 1:20 dilution of 0.25 mg/mL DAPI to each sample.

#### First round of FACS sorting for EdU enrichment

28. Prepare 96-well plates with 4 μl NSB-RNA per well.
29. Sort 250~500 nuclei per well. Nuclei are gated based on Alexa647 and DAPI staining.
30. Briefly centrifuge the plate after the sort.

#### Reverse Transcription

31. For each well of the 96-well plate, add
    a. 0.5 μL 10 mM dNTP
    b. 1 μl indexed oligo-dT primers (50 μM)
32. Incubate plates at 55°C for 5minutes. Immediately place plates on ice after incubation.
33. Prepare RT mix (prepare for one 96-well plate) and distribute 3.5 μL to each well. Pipet up and down once.

| Reagent | 1X (μL) | 110X (μL) |
|---|---|---|
| 5X SuperScript™ IV Reverse Transcriptase Buffer | 2 | 220 |
| 100 mM DTT | 0.5 | 55 |
| SuperScript™ IV Reverse Transcriptase | 0.5 | 55 |
| RNaseOUT™ Recombinant Ribonuclease Inhibitor | 0.5 | 55 |
| Total | 3.5 | 385 |

34. Start the RT reaction with the following program:
    a. 4°C for 2 minutes
    b. 10°C for 2 minutes
    c. 20°C for 2 minutes
    d. 30°C for 2 minutes
    e. 40°C for 2 minutes
    f. 50°C for 2 minutes
    g. 55°C for 15 minutes
35. After the reaction, briefly centrifuge the plate and add 1μL 18mM EDTA to each well to step the reaction.

36. Pool the nuclei with P20 wide bore tips and 1 mL pipette, and transfer into one 1.5 mL tube. Keep nuclei on ice.

#### Second round of FACS Sort for EdU enrichment

37. Add 1:20 dilution of 0.25 mg/ml DAPI to each sample.
38. Prepare two 96 well plates with 5 µL EB buffer per well.
39. Sort 25 nuclei per well per 96-well plate in the first sort (i.e. if two 96-well plates were sorted and combined in the first sort, then double the cell number per well to 50 nuclei for the second sort). Nuclei are gated based on Alexa647 and DAPI staining.
40. Brief centrifuge the plate
41. Plates can be stored in -80°C or directly proceed to the next step.

#### Second Strand Synthesis

42. Prepare second strand synthesis mix
    a. Each well needs ⅔ µL of Second Strand Synthesis Buffer and ⅓ µL of Second Strand Synthesis Enzyme.
    b. For each 96-well plate, prepare 110X second strand synthesis mix:
        i. 73.3 µL of Second Strand Synthesis Buffer
        ii. 36.7 µL of Second Strand Synthesis Enzyme
43. Add 1 µL reaction mix per well and pipe up and down once.
44. Perform second strand synthesis at 16°C for 1 hour.

#### Tagmentation

45. Prepare the Tagmentation mix: for each 96-well plate, combine 600 µL tagmentation buffer + 1.5 µL self-loaded Tn5. Mix well.
46. Add 6 µL tagmentation mix to each well.
47. Perform tagmentation at 55°C for 5 minutes.

#### SDS treatment

48. Prepare 1% SDS and BSA mixture for one 96-well plate:

| Reagent | 1X (µL) | 110X (µL) |
|---|---|---|
| 1% SDS | 0.4 | 44 |
| BSA | 0.4 | 44 |

Add 0.8 µL to each well. Pipe up and down to mix.

49. Add 2 µL indexed P5 primer into each well, and incubate the mixture at 55°C for 15min.
50. For each well, add a 3.2 µL mixture including 2 µL 10% Tween-20 and 1.2 µL $H_2O$ to quench SDS.
51. For each well, add 2 µL P7 primer.

#### qPCR and PCR

52. Perform qPCR to decide the number of cycles for amplification.
    a. Take out 1 well, add 20 μL NEBNext® High-Fidelity 2X PCR Master Mix and 0.4 μL dsGreen for real-time PCR
    b. qPCR program
       72°C for 5 minutes
       98°C for 30 seconds
       20 cycles of (98°C for 10 seconds, 66°C for 30 seconds, 72°C for 30 seconds)
       72°C for 5 minutes
53. Amplification was carried out using the same program with the determined cycle.
54. Pool the PCR products and stored at -20°C .

#### Ampure beads purification (two rounds of 0.8X)

55. Put the Ampure beads in RT 30 minutes before the experiment.
56. Take 800 μL PCR product for a 0.8X Ampure XP beads purification by adding 640 μL Ampure XP beads.
57. Incubate at room temperature for 5 minutes and briefly centrifuge the tube.
58. Put the tube on a magnetic stand for 5 minutes and remove supernatants.
59. Wash the beads with 500 μL 80% ethanol twice.
60. Centrifuge the tube and carefully remove all supernatants from the tube.
61. Add 105 μL EB buffer, vortex, and incubate the tube at room temperature for 3 minutes.
62. Briefly centrifuge and leave the tube on a magnetic stand for 3 minutes.
63. Transfer 100 μL of the supernatant to a new tube.
64. Add 80 μL Ampure beads and repeat another round of 0.8x Ampure beads purification.
65. Elute the product to 20 μL EB buffer.

#### Prepare Library

66. Determine library concentration by Qubit.
67. Visualized library by electrophoresis on a 2% E-gel.
68. Sequence on Novaseq/NextSeq platform.

#### Representative gel image

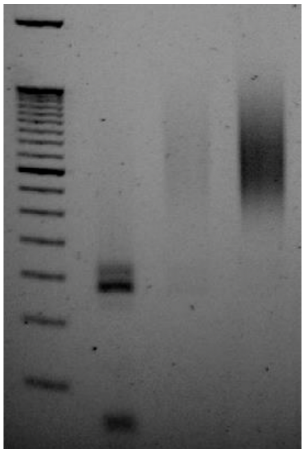

Lanes from left to right:
1. E-Gel™ 50 bp DNA Ladder
2. Pooled PCR product without Ampure beads purification
3. 0.8X Ampure beads purified product from 2.
4. 0.8X Ampure beads purified product from 3. Final library to be sequenced.

*TrackerSci-ATAC*

#### Buffers Preparation

1. Nuclei suspension buffer-ATAC (NSB-ATAC)
    a. NB + 0.1% IGEPAL-CA630 + 0.1% Tween-20 + 1X Protease Inhibitor
    a. Made fresh every time.
2. 2X Stop Buffer (Prepared fresh everytime):

| Reagent | Stock Con. | Volume |
|---|---|---|
| EDTA | 40 mM | 400 uL |
| Spermidine | | 0.78 uL |
| Nuclease free water | | 4.6 mL |

#### EdU labeling and nuclei isolation on cultured cells

Same as documented in *TrackerSci-RNA* protocol

#### EdU labeling and nuclei isolation on mouse brain tissues

Same as documented in *TrackerSci-RNA* protocol, unless skipping the sonication step to avoid chromatin damage

#### Nuclei Fixation

Same as documented in *TrackerSci-RNA* protocol

#### Click -iT Tagging

Same as documented in *TrackeSci-RNA* protocol, unless replacing SUPERase•In™ RNase Inhibitor with 1X Protease Inhibitor

#### First round of FACS sorting for EdU enrichment

3. Prepare 96-well plates with 4 μL NSB-ATAC per well.
4. Sort 250~500 nuclei per well. Nuclei are gated based on Alexa647 and DAPI staining.
5. Briefly centrifuge the plate after the sort.

#### Indexed transposition

6. Add 5 μL TD buffer and 1uL indexed Tn5 enzyme into the sorted plate. Keep the plate on ice.
7. Perform tagmentation at 55°C for 30 minutes on the PCR machine.

8. After tagmentation, place the plate on ice, and add 11 µL 2X stop reaction buffer to each well
9. Pool the nuclei with P20 wide bore tips and 1 mL pipette, and transfer into one 1.5 mL tube. Keep nuclei on ice.

#### Second round of FACS Sort for EdU enrichment

10. Add 1:20 dilution of 0.25 mg/ml DAPI to each sample.
11. Prepare two 96 well plates with 5 µL EB buffer per well.
12. Sort 25 nuclei per well per 96-well plate in the first sort (i.e. if two 96-well plates were sorted and combined in the first sort, then double the cell number per well to 50 nuclei for the second sort). Nuclei are gated based on Alexa647 and DAPI staining.
13. Brief centrifuge the plate

#### Reverse crosslinking

14. After sorting, add 1 µL reverse crosslinking master mix to each well and incubate at 65°C for 16 hours.
15. Spin down the plate, and add 2 µL 10% Tween-20 to each well to quench the SDS.

#### qPCR and PCR

16. For each well, add 1 µL indexed P5 (10 µM) and 1 µL indexed P7 (10 µM) primers
17. Perform qPCR to decide the number of cycles for amplification.
    a. Take out 1 well, add 20 µL NEBNext® High-Fidelity 2X PCR Master Mix and 0.4 µL dsGreen for real-time PCR
    b. qPCR program
       72°C for 5 minutes
       98°C for 30 seconds
       20 cycles of (98°C for 10 seconds, 66°C for 30 seconds, 72°C for 30 seconds)
       72°C for 5 minutes
18. Amplification was carried out using the same program with the determined cycle.
19. Pool the PCR products and stored at -20°C .

#### Sequencing Library Purification

16. Combine PCR products into a 1.5 mL tube.
17. Perform column purification using Zymo DNA Clean & Concentrator kit and elute in 20 µL EB.
18. Determine library concentration by Qubit.
19. Visualized library by electrophoresis on a 2% E-gel.
20. Sequence on Novaseq/NextSeq platform.

#### Representative gel image

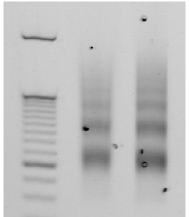

Lanes from left to right:
1. E-Gel™ 50 bp DNA Ladder.
2. Pooled PCR product.

3. Library after column purification from 2. Final library to be sequenced
